# Stony coral tissue loss disease (SCTLD) destabilizes the coral microbiome

**DOI:** 10.1101/2025.06.19.660612

**Authors:** Shrinivas Nandi, Timothy G. Stephens, Kasey Walsh, Rebecca Garcia, Maria F. Villalpando, Rita I. Sellares-Blasco, Ainhoa L Zubillaga, Aldo Croquer, Debashish Bhattacharya

## Abstract

Stony coral tissue loss disease (SCTLD) is a rapidly spreading lethal coral disease, the etiology of which remains poorly understood. In this study, using deep metagenomic sequencing, we investigate microbial and viral community dynamics associated with SCTLD progression in the Caribbean stony coral *Diploria labyrinthiformis*. We assembled 264 metagenome-assembled genomes (MAGs) and correlated their abundance with disease phenotypes, revealing significant shifts in both the prokaryotic microbiome and virome. Our results provide clear evidence of microbial destabilization in diseased corals, suggesting that microbial dysbiosis is an outcome of SCTLD progression. We identified DNA viruses that increase in abundance in infected corals and are present in SCTLD-affected corals at other sites. In addition, we identify the first putative instance of asymptomatic/resistant SCTLD-affected colonies, suggesting potential microbial induced resilience (i.e., beneficial microbiome). Finally, we propose a mechanistic model of SCTLD progression, in which viral dynamics may contribute to a microbiome collapse. These findings provide novel insights into SCTLD pathogenesis and offer consistent molecular signals of disease across diverse geographic sites, presenting new opportunities for disease monitoring and mitigation.

## Background

In the past decade, coral reefs in the Caribbean have emerged as a hotspot of coral diseases [1, 2]. Stony coral tissue loss disease (SCTLD) was first observed in 2014 in the Florida Keys [3]. Sediment resuspension caused by a dredging operation in Miami Harbor is believed to have contributed to the emergence of this disease [4], however the exact cause remains unknown. SCTLD infects 22 different coral species with varying levels of susceptibility [5]. Signs of active SCTLD infection include focal or multifocal lesions that spread at chronic to acute rates, followed by tissue loss [3, 5]. The SCTLD epidemic has decimated affected reefs, with a fatality rate of 73-100% [6] and is transmissible through both direct contact and seawater, even from the ballast of ships [4, 7–11]. Since 2014, SCTLD has spread throughout Florida’s coral reefs, and is now found in Mexico, Turks and Caicos, the U.S. Virgin Islands, the Dominican Republic, and many other sites across the Caribbean [12–15].

No SCTLD-associated vector has yet been identified that satisfies Koch’s postulates [16]. Elucidating the cause of coral diseases has been a challenge, wherein of 18 known coral diseases, Koch’s postulates have been satisfied for only five [17]. A vast proportion of initial SCTLD studies focused on utilizing 16S rRNA sequencing to identify causative prokaryotes [5]. These studies identified many orders of bacteria that are strongly correlated with infection, including Flavobacteriales, Clostridiales, Rhodobacterales, Alteromonadales, Vibrionales, Rhizobiales, and Campylobacterales [5, 18–21]. These findings, paired with an alleviation of the disease upon treatment with antibiotics [22–24] and probiotics [25, 26] initially suggested a bacterial causative agent. However, a single causative bacterial pathogen present in all infected coral species could not be identified, instead, the data suggested that a polymicrobial consortium may be involved in SCTLD [27, 28].

Whereas these 16S rRNA-based studies provided novel insights, they have largely excluded other potential causative agents, such as viruses and eukaryotes. For example, one study identified an increased abundance of the micro-eukaryote *Cillaphora* in SCTLD-infected corals [28]. Viral-like particles (VLPs) have also been observed using microscopy within the algal endosymbionts of SCTLD-infected corals [29]. Which, when paired with metatranscriptomics, led to the hypothesis that a viral agent may induce endosymbiont dysfunction and autophagy [30]. RNA viruses, specifically *coral-associated alphaflexiviruses*, have also been linked to diseased corals [31]. A few studies have assessed the role of DNA viruses in SCTLD, however only a handful of viral contigs were identified in association with the disease phenotype [32]. However, other studies have reported the presence of filamentous viral-like particles in endosymbionts from regions where SCTLD is not present [33], raising questions about their role in the disease. Additionally, histopathological studies have shown discrepancies in lesion severity and cellular involvement, suggesting regional differences and variable disease presentations [34].

In this study, we performed holobiont metagenomic analysis of *Diploria labyrinthiformis*, a SCTLD-susceptible coral present in reefs of the Dominican Republic [16]. Samples were collected from infected and healthy colonies near and far away from the disease lesion and analyzed using Illumina metagenomic sequencing and metagenomic analysis. These data were used to derive a putative model of SCTLD progression, in which a viral infection of the holobiont is hypothesized to cause dysbiosis of the microbiome, making the coral more vulnerable to a secondary infection by opportunistic pathogens. We propose that these biotic shifts result in the visual symptoms of SCTLD.

## Methods

### Field Sampling

Samples of the coral *D. labyrinthiformis* were collected from Playita Reef (18.373081 - 68.853475), Bayahibe, Dominican Republic on June 12, 2024 **(Figure 1A).** The samples were taken from an average depth of 5.82 m using SCUBA. One replicate was taken for each colony. Five healthy colonies (hereinafter “HC”), apparently unaffected by SCTLD, were collected (**Figure 1B**). Five infected colonies (hereinafter “IC”), bearing macroscopic signs of SCTLD [35], were also collected. Two samples were collected from each IC, one from the active lesion (hereinafter “DS” [“Diseased Site”]) and one about 2 inches from the active lesion (hereinafter “AH” [“Apparently Health”]) (**Figure 1C**). After collection, samples were stored in a cooler filled with seawater and brought back to shore and processed. Approximately 0.25 mg of each sample was stored in 2 mL tubes with 2 mm silica beads and preserved using DNA/RNA shield stabilization fluid. Samples were subsequently flown back to Rutgers University within 2 days after sampling where they were stored at -80°C.

**Figure 1:**
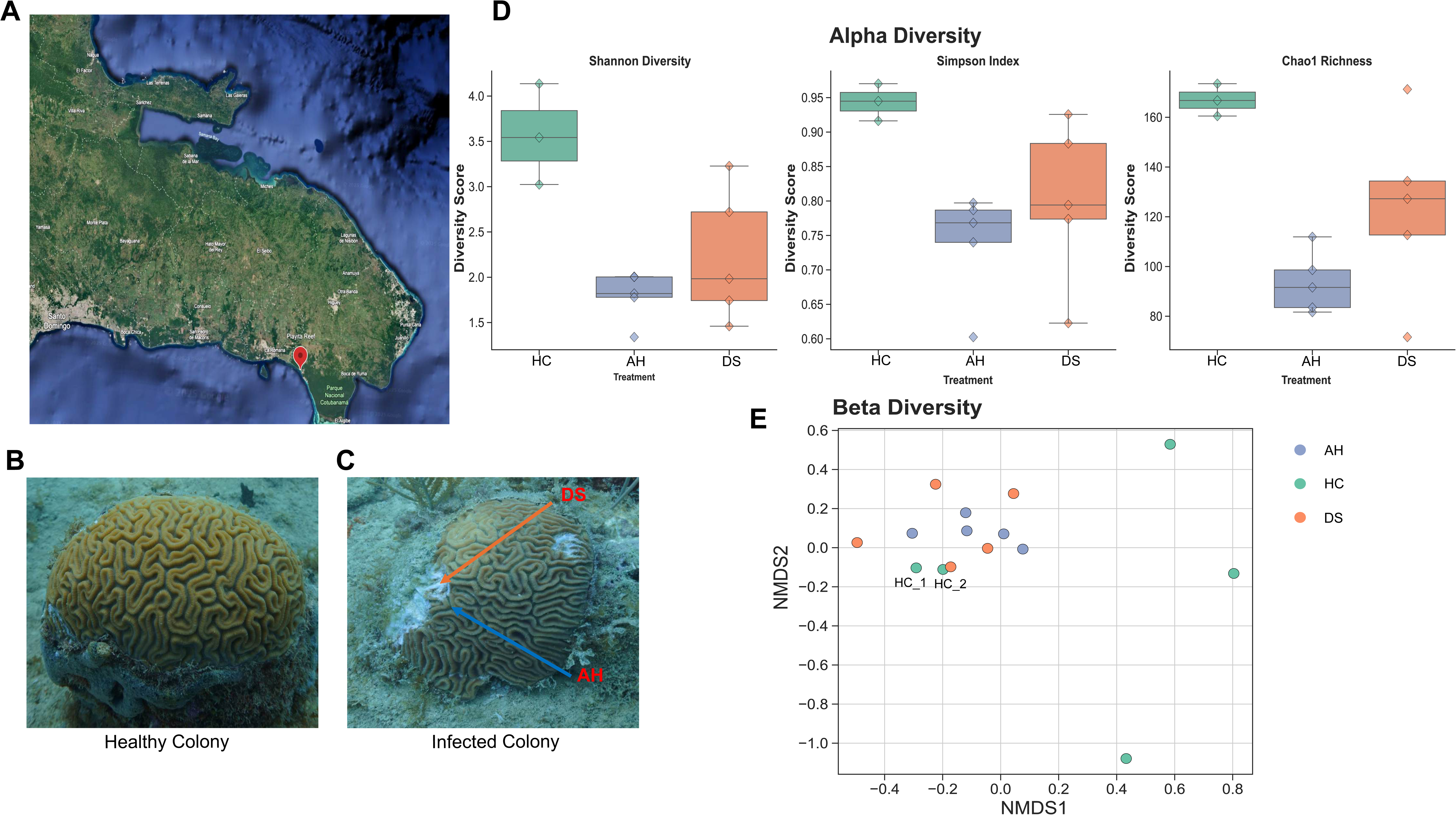
Experimental design and microbiome ecological diversity parameters. **(A)** Satellite image showing the coordinates of sample collection in Playita Reef (18.373081, - 68.853475), Dominican Republic. Image taken from Google Earth. **(B)** Image of a healthy *D. labyrinthiformis* colony, i.e., no signs of SCTLD. Scrapings shown are those caused during sample collection. **(C)** Image of an infected *D. labyrinthiformis* colony, the blue arrow indicates where the apparently healthy (AH) samples were collected, and the orange arrow shows where diseased samples (DS) were collected. **(D)** Alpha diversity metrics (Shannon Diversity, Simpson index, and Chao1 richness) for all MAGs from the three sample types (Healthy [H]: Green, Apparently Healthy [AH]: Blue, and Diseased [DS]: Orange). Asymptomatic colonies N31 and N37 were removed from the healthy samples for this plot. **(E)** Beta diversity metric Aitchison distance was computed for all samples. The color code is the same as (D). A clear clustering of “healthy” samples HC_1 and HC_2 (labeled) with infected colonies is observed.

Ideally, a more extensive sampling of different colony genotypes exhibiting macroscopic signs of SCTLD-related health states would have been done to strengthen the results described in this study. However, the dire situation faced by reefs in the Dominican Republic, with low population sizes for many species, led us to reduce our effort to minimize damage to surviving corals. Our overarching goal was to identify potential biomarkers of SCTLD for future analyses that could be tested more broadly using non-destructive sampling approaches, such as analysis of coral mucus.

### DNA extraction and Sequencing

Previous coral microbiome studies have focused on microbe-rich mucus excreted by the host. However, this approach may fail to enrich for intracellular microbes and viruses, which are one of the hypothesized infectious agents in SCTLD [29, 32]. We therefore homogenized and extracted DNA from the collected coral samples (host and mucus combined) using the ZymoBIOMICS DNA extraction kit, based on the protocol provided by the manufacturer, with the average yield being 1900 ng of total DNA. Samples were sent in two batches for sequencing, the first six samples were sent to assess the extraction procedure, and once validated, the remaining samples were sent for processing. Sequencing was done by Azenta Life Sciences using paired-end (2×150 bp) Illumina shotgun metagenomic sequencing reagents on a NovaSeq X+, targeting 90 Gbp of read data for each sample.

### Construction of Metagenome assembled genomes (MAGs)

Metagenome Assembled Genomes (MAGs) were constructed for prokaryotes, eukaryotes, plasmids, and viruses using a custom work termed Naïve ATLAS (https://github.com/TimothyStephens/naive_atlas). This workflow combined the highly automated ATLAS workflow (https://github.com/metagenome-atlas/atlas; only designed for prokaryotes) [36] with the taxonomically agnostic (but less automated) VEBA v2 (https://github.com/jolespin/veba) workflow [37]. Whereas both workflows have been extensively described in their respective publications and GitHub pages, a detailed description of the Naïve ATLAS workflow is presented in the supplements (see **Supplemental Text**).

Additionally, read mapping was used to assess the relative abundance and putative taxonomy of the algal endosymbionts present in each sample (see **Supplemental Text**). All downstream analyses were performed using Python v3.9.7 with the base packages, pandas (v2.2.3), and numpy (v1.26.4). Plots were generated using matplotlib (v3.6.2) and seaborn (v0.12.1). Statistical tests were performed using SciPy (v1.13.1) unless specified otherwise.

### Species Diversity, Relative Abundance, and Differential Abundance analysis

Alpha (Shannon diversity, Simpson evenness index, and Chao1 richness) and beta diversity metrics (Aitchison Distance) were calculated using the approaches described in the **Supplemental Text**. Non-metric multidimensional scaling (NMDS) was used to assess sample clustering. Finally, PERMANOVA and PERMDISP were used to assess the effect of sample variation. DESeq2 was used for differential abundance analysis (see **Supplemental Text** for a detailed justification), with additional TPM-based foldchange assessment (log_2_[DS or AH TPM] – log_2_[HC TPM]; see below) and visualization to confirm the trends observed in the data. TPM values were recalculated separately for prokaryotes (pMAGs) and viruses (vMAGs) in each sample to prevent shifts in one affecting the apparent relative abundance of MAGs in the other. This allowed independent assessment of the prokaryotic microbiome (hereinafter *microbiome*) and the viral microbiome (hereinafter *virome*). Microbiome shifts were evaluated at the class level, while virome shifts were assessed at the phylum level (based on the Baltimore classification). Statistical significance was determined using the Mann-Whitney U test.

### Selecting MAGs associated with SCTLD

MAGs were postulated to be associated with SCTLD if they were differentially abundant (|FC| > 2.0 and *p*-adjusted value < 0.05) in AH and DS samples compared to HC samples. Additionally, MAGs were only considered to be biologically relevant if they had a median relative abundance > 5% in either healthy or infected colonies. Using these thresholds, a set of pMAGs and vMAGs were selected that appeared to have a significant shift in infected colonies (AH and DS) compared to healthy colonies (HC). TPM based box plots were generated for each selected MAG to visualize and interrogate the apparent shifts in abundance.

### Verifying vMAGs and putative host identification

To confirm that all scaffolds assigned to each of the selected vMAGs are of viral provenance, and to identify putative hosts for each vMAG, a phylogenetic approach was applied to each protein predicted in each vMAG. This analysis used the nr (2022_07) and IMG/VR (v4.1) databases and the DIAMOND blastp (v2.1.10.164; --ultra-sensitive --max-target-seqs 0 -- evalue 1e-05) [38], MAFFT (v7.520; --auto) [39], trimAl (v1.5.rev0; --automated1) [40], iqtree (v2.3.6; -m TEST -bb 1000) programs. A detailed description of the methods is presented in the **Supplemental Text**.

### Downstream assessment of prokaryotic MAGs

The taxonomy and quality of pMAGs was assessed using the Naïve ATLAS workflow (described above). Functional annotation of proteins from selected pMAGs was performed using GhostKOALA [41]. Furthermore, virulence capabilities of these pMAGs were assessed using a DIAMOND blastp search against the virulence factor database (VFDB 2022) [42].

### Assessing other SCTLD metagenomic studies

To test the association of our selected MAGs with SCTLD, we assessed their presence in the only other SCTLD metagenomic dataset available, NCBI BioProject PRJNA576217 [28]. These authors generated 58 short-read metagenomic datasets from four Floridian coral species: *Diploria labyrinthiformis*, *Stephanocoenia intersepta*, *Meandrina meandrites*, and *Dichocoenia stokesii*. These samples were annotated as Diseased Lesion (originally “DL”, but hereinafter “DS” to match the format of the current study), Disease Unaffected (“DU”, hereinafter “AH”) and lastly, Apparently Healthy (“AH”, hereinafter “HC”). Briefly, reads were mapped against all our MAGs using bowtie2 (--local) (v2.5.4) [43], with coverage calculated using CoverM (--genome) (v.0.6.1) [44]. A MAG was present in a species if it had a TPM > 1.0 in at least 5 samples (out of ∼15 samples per species). See **Supplemental Text** for a full description.

## Results

### General metagenome analysis

A detailed description of the sequencing and MAG construction results is presented in the **Supplemental Text**. Briefly, a total of 264 MAGs were identified, two are eukaryotic (although highly incomplete and not used for downstream analysis), 85 are prokaryotic MAGs (pMAGs; **Supplemental Table 1**), 36 are plasmid MAGs, and 141 are viral MAGs (vMAGs; **Supplemental Table 2**). Of the 141 vMAGs, 109 were classified as Caudoviricetes, which are widely distributed bacteriophages. Endosymbiont composition was also assessed in the colonies, indicating the presence of *Breviolum minutum*, *Durusdinium trenchii* and *Symbiodinium necroappetens* (only in IC) [for details see **Supplemental Text**].

### Microbial species metrics

Alpha diversity was assessed using the Simpson index (evenness), Shannon diversity (diversity), and Chao1 score (richness). In all three indices, HC samples had the highest diversity and richness (0.944 ± 0.027, 3.568 ± 0.557, and 166.917 ± 6.502, respectively), followed by DS samples (0.8 ± 0.117, 2.227 ± 0.729, and 123.431 ± 36.127), and lastly, AH samples (0.739 ± 0.079, 1.79 ± 0.272, and 93.46 ± 12.37) [**Supplemental Table 3, Figure 1D**]. Kruskal-Wallis tests revealed significant differences in Simpson (*p*-value = 0.037) and Chao1 (*p*-value = 0.034) metrics, whereas the Shannon index was not statistically significant (*p*-value = 0.054). Pairwise comparisons using the Mann-Whitney U test showed a significant difference (*p*-value = 0.035) in all three-diversity metrics between AH and HC samples. However, no statistically significant differences were observed between DS and HC samples.

An NMDS plot **(Figure 1E)** of the MAG beta diversity between each sample showed that the infected colonies (AH and DS) clustered tightly together irrespective of genotype, suggesting they had similar microbiome profiles. Of the healthy colonies, three were distributed along both axes and did not group with other samples, suggesting they had diverse and unique microbiome compositions. Two of the healthy colonies, which had no visible lesions at the time of collection (HC_1 and HC_2 in **Figure 1E**), clustered tightly with the infected colonies. Furthermore, upon reinspection of these two “apparently” healthy colonies in the field nine months after initial sampling, both were still healthy, with no visible symptoms of SCTLD [**Supplemental Figure 1**]. Therefore, HC_1 and HC_2 were removed from subsequent analyses, unless otherwise specified. PERMANOVA was used to evaluate the significance of microbial beta-diversity across the three sample groups, i.e., HC, AH, and DS (pseudo-F = 1.63, n = 13, *p-value* = 0.008, permutations = 999). To confirm these findings, PERMDISP was also calculated (F-value = 13.57, n = 13, *p-value* = 0.005, permutations = 999). These results imply that the dispersion of samples are driving the results, which may be attributed to microbial taxon variation in the healthy microbiome [**Figure 1E**].

### Microbiome profile shifts under infection

Class-level relative abundance analysis of the microbiome identified Alphaproteobacteria and Gammaproteobacteria as being highly prevalent across all samples [**Supplemental Table 1; Figure 2A**; see **Supplemental Text**]. In AH and DS (but not HC) samples, Leptospirales was also highly prevalent, reaching statistical significance (*p*-value = 0.035 for both) compared to HC. Differential abundance analysis of the 85 pMAGs using DESeq2 (|log[JFC| > 2.0 and an adjusted *p*-value < 0.05) identified three which significantly increased in infected colonies (AH and DS samples). Eight and seven pMAGs showed significantly decreased abundance in DS and AH samples (respectively, with the later a subset of the former). Fold change analysis (not statistically significant) identified 16 pMAGs with increased abundance and 69 pMAGs with decreased abundance in infected colonies [**Figure 2B**]. Differentially abundant pMAGs with a median relative abundance > 0.1 in at least one health sample were considered of biologically interesting. As a result, MAG_prokaryotic_02 (hereinafter pMAG02) and MAG_prokaryotic_01 (hereinafter pMAG01) were selected for downstream analysis.

**Figure 2:**
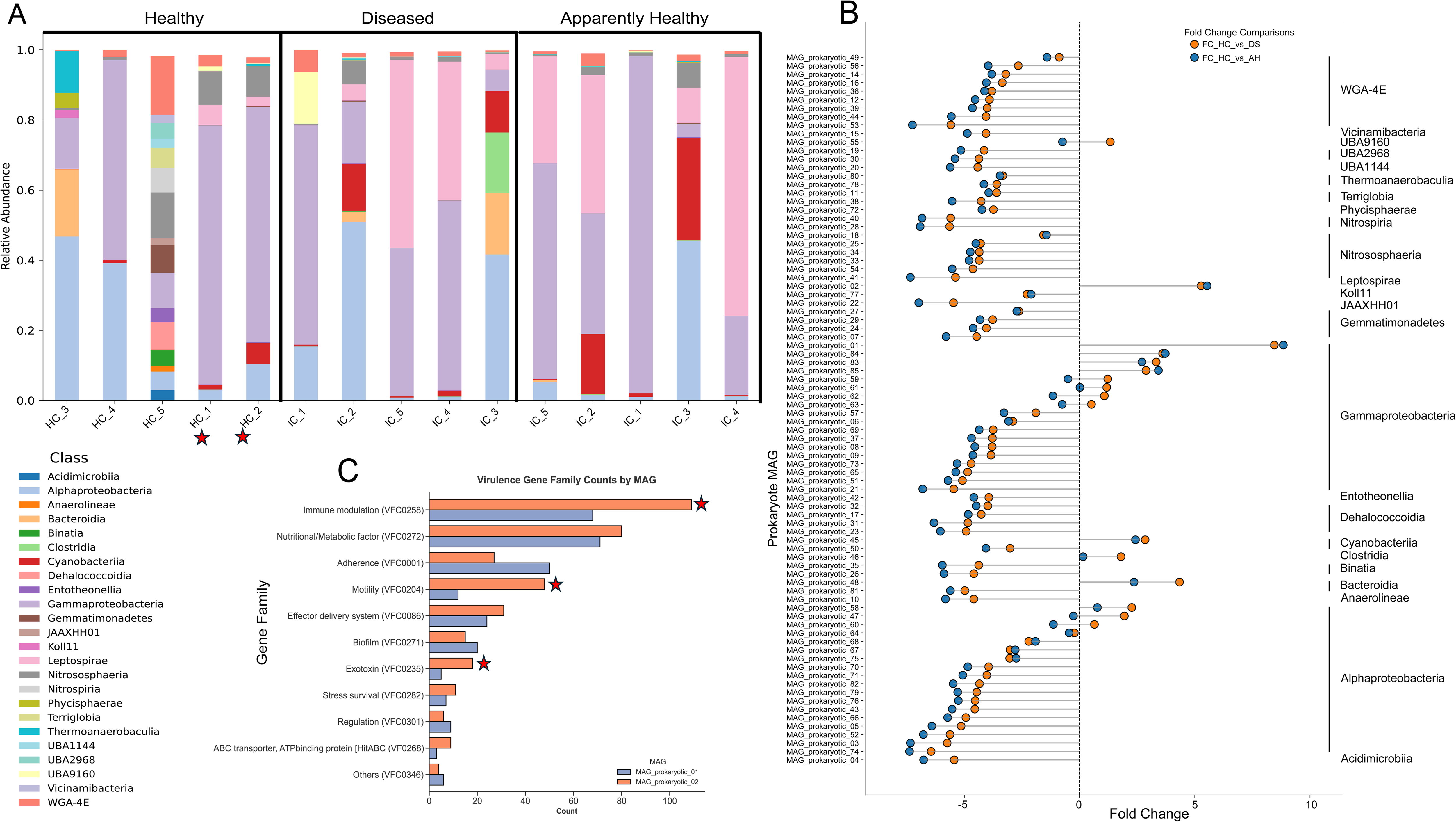
Shifts in prokaryotic component of the microbiome. **(A)** Relative abundance (Y-axis) at the class level of the prokaryotic microbiome plotted as a stacked bar graph. The X-axis represents all samples, with the respective colony number, either HC (healthy colony) or IC (Infected Colony). The graph is divided into three parts, headers provided on the top, each representing the health status, i.e., healthy (H), apparently healthy (AH), or diseased (DS). The red stars show the two putative asymptomatic colonies in this study. A legend with the color used for each class is shown on the bottom left of the image. Some samples may not add up to 1.0 because the sample has some pMAGs that do not have class level taxonomic classification. **(B)** Lollipop plot showing the log_2_FC (X-axis) for each prokaryotic MAG (Y-axis). MAGs are grouped by taxonomic class, shown on the right of the panel. The orange points represent the change in abundance between the H and DS samples, and the blue points represent the change in abundance between the H and AH samples. **(C)** Gene abundance plot of virulence factor genes from the two pMAGs of interest: Leptospirales (orange) and Gammaproteobacteria (blue). For each gene family (Y-axis) the respective VFDB ID is shown, with the number of genes identified in those families in each pMAG shown on the X-axis. The red stars show key virulence pathways, wherein pMAG02 has more genes than pMAG01.

### pMAGs involved in disease phenotype

pMAG02 had a CheckM completeness of 96.82% and a contamination of 1.05%, and pMAG01 had a completeness of 72.68% and a contamination of 0.54%. The lowest GTDB taxonomy level annotated to pMAG02 was in the order Leptospirales. pMAG01 was annotated as a Gammaproteobacteria, with a genus level annotation of JANQNX01 (an uncultured Gammaproteobacterium, previously identified in Australia). We assessed the virulence proteins in each of these pMAGs using VFDB. A total of 393 proteins were identified as virulence factors in pMAG02 and 305 proteins in pMAG01 [**Supplemental Table 4**]. Both MAGs had similar nutrient/metabolic factor (VFC0272) at 80 and 71 [**Figure 2C**], respectively. pMAG02 contained 109 immune modulation genes (VFC0258), whereas pMAG01 contained 69. Furthermore, pMAG02 had more motility (48) genes (VFC0204) compared to pMAG01 (12). Lastly pMAG02 also contained more (18) exotoxin genes (VFC0235), compared to pMAG01 (12) [red stars in **Figure 2C**]. pMAG02 also contained key exotoxins that have been previously associated with virulence in other species, such as hemolysins *tlyA* and *tlyC*. Furthermore, we assessed prokaryotes that have been previously associated with SCTLD, namely Rhodobacterales, Rhizobiales, and *Vibrio* spp. We identified three pMAGs annotated as Rhodobacterales, which had a median relative abundance in the order of 1e^−05^. No pMAGs were annotated as Rhizobiales or *Vibrio* spp.

### Virome profile shifts under infection

An initial relative abundance analysis of the virome was performed at the “phylum” taxonomic level. Out of the 141 viral MAGs identified in our dataset, 109 were classified as Caudoviricetes (phylum Uroviricota) [**Supplemental Table 2]**. Seven other phyla were detected in our samples, namely, Duplodnaviricota, Lenarviricota, Nucleocytoviricota, Peploviricota, Phixviricota, Pisuviricota, and Preplasmiviricota [for in-depth phylum level analyses see **Supplemental Text, Supplemental Figure 2**].

Differential abundance analysis of vMAGs identified 17 with significantly increased abundance in AH samples (compared to HC samples) and 16 in DS samples, with the former overlapping completely with the latter [see **Supplemental Text, Supplemental Figure 3**]. To assess the biological relevance of the differentially abundant vMAGs, we examined their relative abundances in colonies with different health states. Among the 17 vMAGs with increased abundance in infected colonies, only five contributed substantially to the overall virome composition based on total median relative abundance, accounting for 91.3% in AH samples and 76.5% in DS samples [**Figure 3A**, **Supplemental Table 2**]. These five dominant vMAGs were MAG_viral_001, MAG_viral_043, MAG_viral_055, MAG_viral_058, and MAG_viral_060 (hereafter known as SCTLD-associated vMAGs). In HC samples, the median abundances of these five vMAGs had a total median abundance of 15.2% of the virome. None of the other 12 vMAGs that had an increased abundance in AH or DS samples had a relative abundance greater than 0.1 and were therefore not considered for downstream analyses.

**Figure 3:**
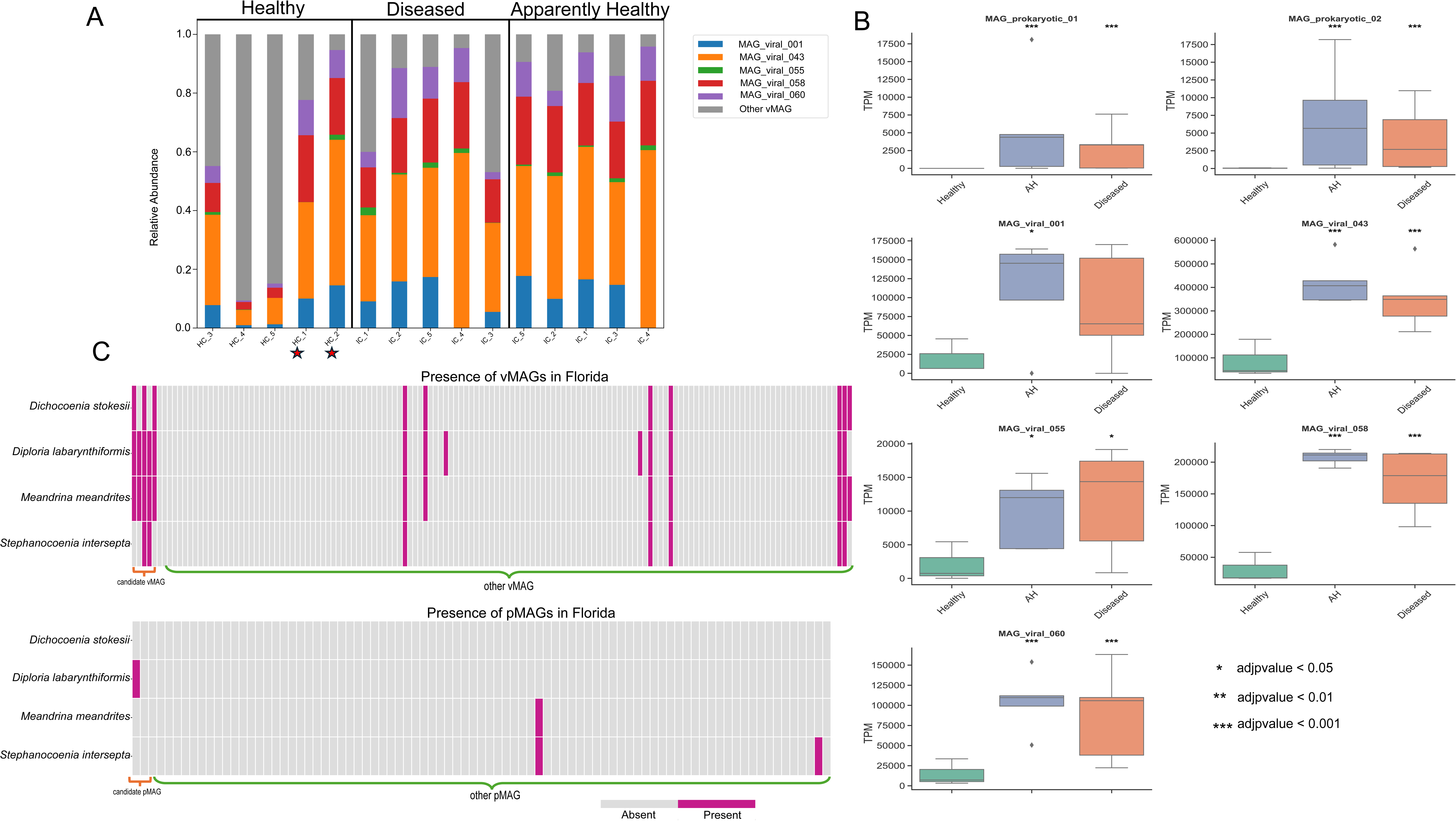
Viral compositional analysis and MAGs correlated with SCTLD. **(A)** Relative abundance (Y-axis) of the SCTLD-associated vMAGs (color) and all other vMAGs have been represented in gray. The X-axis represents all samples, with the respective colony number, either HC (healthy colony) or IC (Infected Colony). The graph is divided into three parts, headers provided on the top, each representing the health status, i.e., healthy (H), apparently healthy, or diseased. The red stars show the two putative asymptomatic colonies in this study. **(B)** Box plots showing the TPM (Y-axis) of the MAGs of interest, across the different health conditions (X-axis). For this analysis, the putatively asymptomatic colonies were removed from the H condition (green); all five samples were used for the AH (blue) and DS (orange) conditions. Stars on the top of each box plot show the statistical significance generated by DeSeq2, compared to the H condition. A legend is presented on the bottom right of the panel. **(C)** Presence/absence heatmap of vMAGs and pMAGs that were generated in this study in SCTLD samples from Florida, USA. Pink indicates presence of signal in more than five samples (an average of 15 samples per species were collected in Florida). Grey indicates a signal was found in <5 samples per species. Orange lines on the x-axis indicate those MAGs associated with SCTLD infection in our study, whereas green indicates MAGs that were not associated with SCTLD infection in this study.

### vMAGs involved in disease phenotype

In the five SCTLD-associated vMAGs, 41 proteins were predicted with only four having functional annotations predicted using GhostKOALA. Three of the proteins are from vMAG055 and were annotated as dcm (K26510), a DNA methyltransferase ERCC 2 (K10844), and a kinesin family member 21 protein KIF21 (K24185). One protein was from vMAG060 and was annotated as an adiponectin receptor ADIPOR. Of the 41 proteins from these five vMAGs, 25 (from four vMAGs [vMAG058 had no hits]) had hits to enough sequences in the nr and IMG/VR databases for phylogenetic analysis [**Supplemental File 1; Supplemental Table 5**]. This analysis identified vMAG001 (Adintoviridae), vMAG043 (Imitevirales), and vMAG060 (unknown taxonomy) as putatively host infecting, and vMAG055 (Megaviricetes) as putatively coral endosymbiont infecting.

### Assessing asymptomatic/resistant colonies

We assessed the relative abundance of our selected vMAGs and pMAGs in the putative asymptomatic healthy colonies (i.e., HC_1 and HC_2). Combined, these vMAGs made up 91.04% of the virome in HC_2 and 77.64% in colony HC_1. Similarly, we observe that pMAG01 had a relative abundance of 0.6387 and 0.6195 (respectively), and pMAG02 had a relative abundance of 0.0252 and 0.0576.

### Validating MAG signatures using data from the Florida SCTLD outbreak

SCTLD metagenomic data from Rosalses et al. [28] was analyzed for the presence of our selected SCTLD-associated MAGs, across the four targeted species: *Stephanocoenia intersepta*, *Diploria labyrinthiformis*, *Dichocoenia stokesii*, and *Meandrina meandrites*. 14 vMAGs (generated in this study) were present in the samples from Florida, which included all five of our SCTLD-associated vMAGs [**Figure 3C]**. Two vMAGs were exclusive to *D. labyrinthiformis* (from Florida), two were detected in two coral species (including vMAG043), four were detected in three coral species (including vMAG001, vMAG058, and vMAG060), and six were detected in all four coral species (including vMAG055). Note, the vMAGs identified in the Florida samples that do form not part of our selected set had low relative abundances (∼ 0.001) in the data generated from the Dominican Republic (this study). A log_2_TPM based trend analysis was performed to assess variation based on health states but was deemed unreliable due to the unknown outcomes of the healthy colonies [**Supplemental Figure 4; Supplemental Table 6**].

Of the generated 85 pMAGs in this study, only three were present in the Florida data. No pMAG was detected in more than two coral species. Of the SCTLD-associated pMAGs (generated in this study), pMAG01 was present only in *D. labyrinthiformis*, whereas pMAG02 was not detected in any species [**Figure 3C]**. pMAG84, annotated as *Bacterioplanoides* sp024397975 (class: Gammaproteobacteria), was detected only in *S. intersepta,* and pMAG50, annotated as *Parasynechococcus* sp002724845 (class: Cyanobacteria), was detected in *S. intersepta* and *M. meandrites*.

## Discussion

### Marked microbiome shifts exist, independent of coral health state

Significant efforts have been made to identify a causative agent of SCTLD, yet fulfilling Koch’s postulates in corals remains a challenge. Of the 18 known coral diseases, only five have satisfied Koch’s postulates, highlighting the challenges facing disease studies in this group [17]. Our research shows that SCTLD infection drives substantial shifts in the prokaryotic microbiome and virome. We observe a clear dispersion of microbial beta-diversity with the clustering of infected colonies in both AH and DS samples, which has been documented in the literature [20, 45]. Alpha-diversity metrics (Shannon diversity, chao1 richness, and Simpson index) demonstrate that healthy colonies maintain rich and diverse microbial profiles which are potentially tailored to their specific micro-environments [46–48]. Consistent with previous studies, we also observe that upon infection, these microbiomes become more uniform, with reduced diversity, evenness, and richness *via* the loss of prokaryotic and bacteriophage communities [18, 45]. We also observe that upon lesion formation (i.e., the DS samples), there is a slight increase in alpha diversity, suggesting a new (potentially opportunistic or pathogenic) microbiome has colonized the infected colonies.

Our results suggest that microbiome uniformity is observable, even in the absence of visible SCTLD lesions. This outcome is found in samples HC_1 and HC_2 which were initially targeted as healthy colonies because no visible symptoms of SCTLD were observed during field sampling. Follow-up observations nine months after initial sample collection showed that these colonies were still alive with no visible signs of SCTLD. In addition, these HC samples had a high load of SCTLD-associated vMAGs, but a different pMAG composition when compared to the infected colonies. These colonies may present a novel case of asymptomatic/resistant SCTLD infections, which has been suggested in previous studies [11, 29]. The presence of viruses, in both asymptomatic and symptomatic samples has been noted in sea-stars [49], however additional life history studies need to be performed to validate these results in SCTLD infected corals. The HC_1 and HC_2 colonies may have had an innate immunity/resistance to the infection, despite containing SCTLD-associated vMAGs. The role of genotype has been well characterized in conferring thermal stress induced bleaching resilience in corals [50–52], but this has not yet been translated to coral diseases. However, preliminary results have shown variations in genotype resilience and susceptibility in other coral diseases such as White Syndrome and White Pox Disease [53–56]. Regardless, this finding demonstrates the potential for treatment of SCTLD and the selection of resistant genotypes. These observations underscore the urgent need to develop diagnostic biomarkers for SCTLD across various coral species to facilitate informed reef management practices.

### DNA viruses appear to play a key role in infection

We observe two clear trends in the virome, first, similar to the microbiome, there are phylum-level shifts in the virome (See **Supplement Text**). This is primarily marked by loss of the bacteriophage Caudoviricetes, potentially linked to the concurrent loss of numerous pMAGs that may serve as hosts, highlighting the complex interplay between microbiome, virome, and coral host [46, 57, 58]. The second trend we observe is the significant increase in five selected SCTLD-associated vMAGs in infected colonies. These vMAGs make up a small component of the virome of HC colonies (∼ 15%). However, a significant increase is observed in the AH and DS samples, ∼ 91% and ∼ 76%, respectively. Furthermore, these vMAGs comprise the majority of the virome in the asymptomatic/resistant colonies and are also observed across four SCTLD-affected corals in Florida, suggesting a connection between these vMAGs and SCTLD.

Phylogenetic analysis of the proteins predicted in each SCTLD-associated vMAG was used to infer a putative viral host. Three of the five SCTLD-associated vMAGs contained genes whose phylogenies included coral- or hydra-associated viral sequences, supporting the possibility of direct infection of the coral host. One vMAG also included endosymbiont sequences (e.g., *Symbiodinium* spp.) in its gene trees, suggesting a potential interaction with the algal symbiont community. However, the frequent presence of prokaryotic sequences in many phylogenies complicates host prediction, as does the uncertain association of the viral sequences from IMG/VR which are interspersed throughout each tree, making it difficult to determine whether the viruses primarily infect the coral, its symbionts, or members of the surrounding microbiome. Furthermore, the limited functional annotation of viral genes in these vMAGs limits our ability to infer infection mechanisms.

### Prokaryotic composition confers resilience or susceptibility

Compared to the healthy colonies, most pMAGs had decreased abundance in infected colonies. In healthy colonies, Alphaproteobacteria had the highest abundance followed by Gammaproteobacteria, both of which have been identified in previous SCTLD studies [20, 27]. However, as the disease progressed, Alphaproteobacteria abundances decline in the AH and DS samples. We observed a decline in the majority of pMAGs in infected colonies relative to healthy, suggesting microbiome destabilization. Two candidate pMAGs had significant increases in infected colonies: a Leptospirales and a Gammaproteobacteria. In HC samples, these bacteria have extremely low (0.013%) median abundance in the microbiome, however combined, in AH samples these two MAGs comprise up to ∼ 48% of the microbiome (Leptospirales ∼ 30% and Gammaproteobacteria MAG ∼ 18%). However, these pMAGs have a clear decline in abundance in DS samples, i.e., upon tissue loss, suggesting a shift in the microbiome at this stage of disease progression. Gene content analysis of these two pMAGs revealed that Leptospirales had the highest relative abundance of virulence factor genes, which is unsurprising given that pathogenic species from the genus *Leptospira* are known to cause leptospirosis in humans and utilize long-chain fatty acids (LFAs) as their sole carbon and energy sources [59]. The Leptospirales MAG identified in this study contained several genes associated with pathogenicity, notably, *tlyA* and *tlyC*, which are hemolysin genes implicated in erythrocyte lysis in model organisms. In addition, key *fli* genes encoding components of the flagellar apparatus, structures that facilitate motility and contribute to host infection, were also present in our Leptospirales MAG [59]. Based on these findings, we hypothesize that this pMAG may be pathogenic, acquiring LFAs from the coral in the AH samples, but upon tissue loss (DS samples), declines in abundance with its host.

Our results point to three distinct microbiome phases: the first, a rich and diverse community found in healthy colonies, the second, a drop in microbiome diversity upon SCTLD infection (tissue still present), and third, colonization by opportunistic/pathogenic microbes (increasing diversity) as the disease progresses. We suggest that many of the microbes involved in this community shift are always associated with the coral, however upon infection, the decline of the health dominant microbiome creates an open niche that opportunistic/pathogenic species exploit. Interestingly, in the two asymptomatic/resistant colonies (HC_1 and HC_2), we observed only marginal levels of Leptospirales. Instead, these colonies were dominated by pMAG01 (Gammaproteobacteria) [**Figure 2A and Supplemental Figure 5**], which had fewer virulence genes (relative to pMAG02). The relatively higher abundance of Gammaproteobacteria, when compared to the Leptospirales in these colonies may thus prevent the progression to symptomatic disease. Moreover, we speculate that the dominant Gammaproteobacteria present in these colonies may function as a natural probiotic (i.e., beneficial microbiome), offering colonization resistance against Leptospirales and thereby mitigating disease progression. We further highlight that Leptopsirales has not been previously associated to SCTLD, supporting the role of prokaryotes as opportunistic pathogens during disease progression [5, 16, 19, 21, 45]. Other prokaryotes associated with SCTLD (e.g., Rhodobacterales, Rhizobiales, and *Vibrio* spp.) were not detected in our study, or were detected at extremely low abundances. This suggests that SCTLD-associated opportunistic pathogens may reflect local conditions [5, 56].

### Pathogenic MAG presence in other SCTLD datasets

The presence of our selected MAGs in other SCTLD-affected coral species was assessed using data from Rosales *et al.* (BioProject PRJNA576217) [28]. All our selected vMAGs were detected in this analysis [**Figure 3C**], with several found across multiple species, consistent with their role in SCTLD. This analysis turned up our selected vMAGs in the HC samples [see **Supplemental Table 6**]. It is unclear whether the fate of healthy colonies was studied by the authors: i.e., whether any of them later developed SCTLD, preventing further assessment of these colonies as putative asymptomatic/resistant. We only detected three pMAGs across any of the four Floridian coral species, of which two were specific to a single species, and one (Cyanobacteria) was present in two species. Overall, we see a more conserved viral pattern across SCTLD-affected coral species than vis-à-vis prokaryotes. This finding suggests that the prokaryote communities associated with this disease are highly geography and host species specific, aligning with the opportunistic pathogen role previously attributed to SCTLD associated prokaryotes [5, 28, 45, 56]. Nonetheless, the detection of SCTLD-associated vMAGs in multiple coral species, collected over 1,000 miles away and several years apart in time, highlights their potential significance in coral disease dynamics.

### A working multi-kingdom model for SCTLD progression

Based on our findings and the results of previous studies, we propose a multi-kingdom model for the progression of SCTLD [**Figure 4**]. The detection and significant enrichment of vMAGs in SCTLD-affected samples across different studies, geographic locations, and coral species suggest a pivotal role for viruses in disease progression. We hypothesize that these viruses infect microbial members of the coral holobiont, triggering dysbiosis and a collapse of the native microbiome, perhaps through the activation of the coral host immune response [5, 56, 60, 61]. This destabilization creates an open ecological niche which opportunistic microbes can occupy. Leptospirales and Gammaproteobacteria performed this role in our dataset, clearly dominating the microbiomes of infected and asymptomatic colonies. These opportunistic microbes may harbor virulence factors (e.g., hemolysins) which cause tissue degradation and contribute to coral mortality. We note that the Leptospirales was not detected in the Florida study, and the Gammaproteobacteria was limited to *D. labyrinthiformis.* Consistent with previous work, we suggest that opportunistic pathogens may comprise polymicrobial consortia that reflect geographic location and coral host [5, 11, 18, 20, 28, 45].

**Figure 4:**
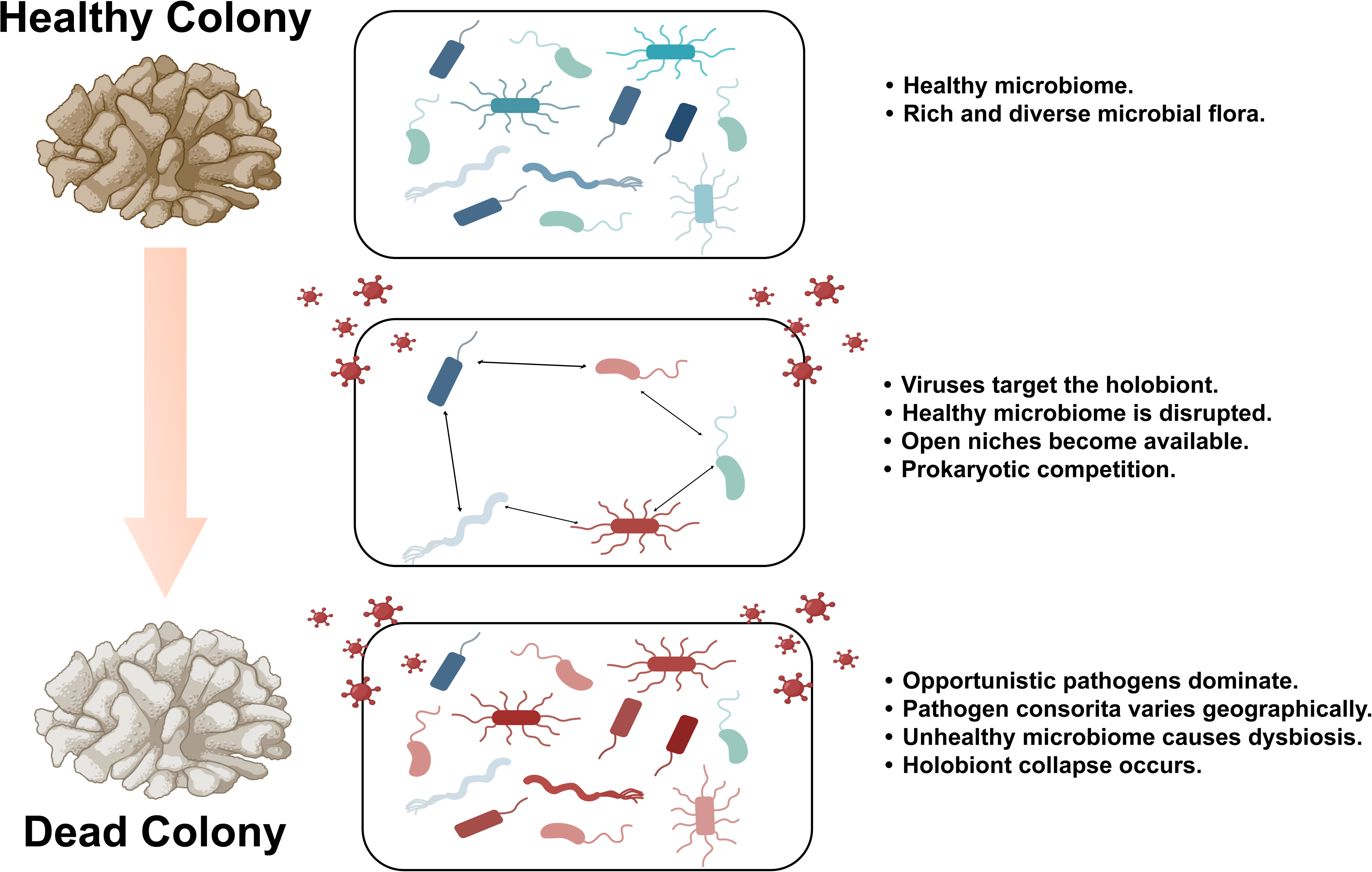
Working model for SCTLD infection. **(A)** A healthy coral holobiont with a rich and diverse microbiome. **(B)** Viruses target components of the healthy holobiont, which either directly or indirectly kill most prokaryotes, opening a niche. (**C**) Previously low abundance bacteria and other opportunistic pathogens flourish and dominate the coral holobiont post-viral infection. Microbiome dysbiosis, caused by the primary viral infection, results in a significant shift in the prokaryotic microbiome towards one dominated by pathogenic bacteria, which is the predominant cause of lesion formation and visual SCTLD symptoms.

Lastly, we highlight the presence of potentially SCTLD asymptomatic/resistant coral colonies, which has been noted in other coral diseases [56, 60]. Apart from putative host resistance, we suggest a microbial resilience hypothesis. Upon microbiome destabilization, if a non-pathogenic taxon (e.g., Gammaproteobacteria in this study) comes to dominate, then it may act beneficially, inhibiting colonization of pathogenic species and promoting disease resilience [58, 62]. This working model aligns with the observed success of probiotic and antibiotic treatments in halting lesion progression in infected colonies [22–25], despite the lack of clear causative prokaryotic agents. That is, anti-microbial interventions may mitigate secondary bacterial infections, allowing the coral host to recover, or the microbiome to stabilize despite an ongoing viral presence.

### Study Limitations and Future Directions

We recognize several important limitations in this study. First, because the viruses of interest have not yet been isolated, it is impossible to assess host specificity. In addition, the small genome size of many vMAGs suggests they may be incomplete, limiting functional interpretation. Nevertheless, the consistent increase in abundance of these vMAGs in infected colonies and their consistent presence in other SCTLD datasets suggests they likely play an important role in disease dynamics. Additional research is needed to better resolve the host range and infection mechanisms of these viruses, ideally through targeted isolation and improved genome completeness. Despite these limitations, our initial findings support the idea that DNA viruses are likely to be significant contributors to SCTLD infection.

Another important outcome of our work is the identification of putative asymptomatic/resistant colonies, which is a novel finding for SCTLD. However, this inference requires validation through long-term life history monitoring, which was beyond the scope of this study and challenging, given the ubiquity of the disease throughout the Caribbean. Future SCTLD and coral disease studies should account for the potential presence of such asymptomatic/resistant states and investigate outcomes for both infected and healthy colonies. Finally, we emphasize that this study does not fulfil Koch’s postulates but rather we propose a working model of SCTLD as a multi-kingdom infection involving both viral and prokaryote components. Further experimental work is needed to establish causality, identify putative viral targets, and generate complete genome data for characterization. Nonetheless, the vMAGs identified here serve as valuable early biomarkers for predicting infection risk or resistance in coral colonies.

## Data Availability

Raw data has been made available on NCBI under the project number PRJNA1278695. All code used in this study is available on https://github.com/shrinivas-nandi/shrinivas-nandi/tree/main/Projects/Dominican_Republic_SCLTD. The Naïve ATLAS pipeline and code is available on https://github.com/TimothyStephens/naive_atlas

## Acknowledgments

Primary funding for this study was provided by the National Philanthropic Trust (23-7825575) in a grant awarded to DB. This work was also supported by grants from the National Science Foundation (2128073) and the USDA National Institute of Food and Agriculture Hatch Formula (NJ01180) awarded to DB. Samples were collected and exported to the US under permit number DJ-CON-1-2025-0005 We wish to acknowledge the help of research interns in the Dominican Republic Marine Innovation Hub for aid in processing samples and the captains of the research vessels who led the different sampling trips.

