## Supplemental Text for "Stony coral tissue loss disease (SCTLD) destabilizes the coral microbiome"

**Methodology**

*Naïve-ATLAS workflow*

Raw metagenomic reads were deduplicated, trimmed, error corrected, and removed if they were putatively from the draft *D. labyrinthiformis* genome (see below) using the bbmap v39.13 package ([https://sourceforge.net/projects/bbmap](https://sourceforge.net/projects/bbmap/)) tools clumpify.sh (dedupe=t dupesubs=2 optical=false), bbduk.sh (qout=33 trd=t hdist=1 k=27 ktrim=r mink=8 trimq=10 qtrim=rl minlength=51 maxns=-1 minbasefrequency=0.05 ecco=t prealloc=t), tadpole.sh (mode=correct aggressive=false tossjunk=f tossdepth=1 merge=t shave=f rinse=f), and bbsplit.sh (maxindel=20 minratio=0.65 minhits=1 ambiguous=best k=13 local=t machineout=t), respectively. The cleaned read pairs were merged using bbmerge.sh (k=62), before being assembled by megahit v1.2 (--k-min 21 --k-max 121 --k-step 20 --min-contig-len 1000 --min-count 2 --merge-level 20,0.98 --prune-level 2 --low-local-ratio 0.2) [1]. The cleaned reads from each sample were mapped against each megahit assembly using minimap2 v1.19.0 (-x sr) [2], with contig abundance calculated using the jgi_summarize_bam_contig_depths (--percentIdentity 95) tool from metabat2 v2.15 [3]. MAG construction was performed in three sequential stages.

(1) Prokaryotic MAGs were identified and extracted from each assembly first. Initially, bins were constructed using the contig abundance information and raw assemblies by MetaBAT2 v2.15 (-m 1500 --minClsSize 150000 --seed 1), MaxBin v2.2.7 (-min_contig_length 1500 -markerset 107) [4], and MaxBin again using the alternative marker set (-min_contig_length 1500 -markerset 40). DAS_Tool v1.1.2 (--search_engine diamond --score_threshold 0.1 --write_bins 1 --create_plots 1) [5] was used to combine the different bins into a unified non-redundant set. Non-prokaryotic contigs were removed from the DAS tools bins using whokaryote.py v1.1.2 (--minsize 1500) [6] and mdmcleaner v0.8.7 (using the “clean” command) [7]. CheckM2 v1.0.2 (using the “predict” command) [8, 9] was used to assess the integrity of the cleaned bins, keeping only those with completeness >50% and contamination <10%. The cleaned and filtered bins from each sample were clustered into a final non-redundant set using skani v0.1 (triangle --sparse --ci --min-af 20) [10] and the representative selection script provided by VEBA (pre-cluster threshold 92.5% and ANI threshold 95%). Taxonomy was assigned using GTDB-Tk v2.4 (using the “identify”, “align”, and “classify” commands; default parameters) [11]. The bins produced by this process constitute the final prokaryotic MAGs (pMAGs) used for downstream analysis.

(2) Next, the contigs not binned in the previous step (i.e., non-prokaryotic) were assessed and used to construct eukaryotic MAGs. Bins were constructed using MetaBAT2 v2.15 (-m 1500 --minClsSize 2000000 --seed 1), with putative non-eukaryotic contigs removed from each bin using whokaryote.py v1.1.2 (--minsize 1500). BUSCO v5.4 (-m genome --auto-lineage-euk --evalue 0.001) [12] was used to assign taxonomy and completeness to each bin; only bins with completeness >10% were retained for downstream analysis. A low completeness cutoff was used due to the challenges associated with binning eukaryotic MAGs. It is difficult to assemble and assess the completeness of short read-based eukaryotic genomes, thus a lower threshold is required, if only to gain a view of which taxa are present across the samples. The eukaryotic bins from each sample were combined into a non-redundant set (95%) of MAGs (eMAGs) using the same approach and cutoffs as for the prokaryotes (skani and the custom VEBA script).

(3) Finally, the contigs not binned in the previous steps (i.e., non-prokaryotic and non-eukaryotic) were used to construct viral and plasmid MAGs. Bins were constructed using MetaBAT2 v2.15 (-m 1500 --minClsSize 2500 --seed 1), with bin completeness and taxonomy assessed using geNomad v1.9.0 (end-to-end --cleanup --verbose --enable-score-calibration --disable-find-proviruses --sensitivity 4.0 --splits 0 --composition auto --min-score 0.7 --max-fdr 1.0 --min-plasmid-marker-enrichment -100 --min-virus-marker-enrichment -100 --min-plasmid-hallmarks 0 --min-virus-hallmarks 0 --max-uscg 100) [13]. Only bins with a False Discovery Rate < 0.05 were retained. Bins from each sample were combined (viruses and plasmids separately) into a non-redundant set (95%) of MAGs (vMAGs and pMAGs, respectively) using the same approach and cutoffs as for the prokaryotes and eukaryotes (skani and the custom VEBA script).

Gene prediction was conducted separately for each type of non-redundant MAG. Of the pMAGs, those taxonomically assigned as bacteria had genes predicted using bakta v1.10.4 (--meta --keep-contig-headers) [14], with those assigned as archaea predicted using prokka v1.14.6 (--addgenes --addmrna --metagenome --kingdom Archaea) [15]. Genes were predicted in eMAGs using the “eukaryotic_gene_modeling_wrapper.py” script from VEBA (using the VEBA custom MicroEuk50 database; --metaeuk_sensitivity 4.0 --metaeuk_evalue 0.01 --pyrodigal_minimum_gene_length 90 --pyrodigal_minimum_edge_gene_length 60 --pyrodigal_maximum_gene_overlap_length 60 --pyrodigal_mitochondrial_genetic_code 4 --pyrodigal_plastid_genetic_code 11 --barrnap_length_cutoff 0.8 --barrnap_reject 0.25 --barrnap_evalue 1e-06 --trnascan_mitochondrial_searchmode='-O' --trnascan_plastid_searchmode='-O'), in vMAGs using prokka v1.14.6 (--addgenes --addmrna --metagenome --kingdom Viruses), and in plMAGs using bakta v1.10.4 (--meta --keep-contig-headers).

To aid MAG construction and reduce overall computational load, a preliminary short-read-only *D. labrynthiformis* reference genome was constructed using the first round of sequencing data. Naïve ATLAS was utilized to clean the read data and assemble contigs. BLASTN (v.2.13.0) was performed against the nt database (2022_07), to assess taxonomy. Blobtools (v 1.1) [16] was used to visualize the taxonomy and subsequently to filter for cnidarian contigs. The resulting cnidarian contigs were filtered for viruses using geNOMAD. The final cleaned cnidarian contigs were used as an internal reference for Naïve ATLAS.

**Assessing algal endosymbiont population**

A database was generated of endosymbionts nuclear genomes, from seven genera [17–24]. Additionally, plastid genome, from Symbiodinium Clade C3 [25] and mitochondrial genome from *B. minutum* [26] were also included in the analyses. All genomes were combined and indexed using Bowtie2 (v2.5.4) . The metagenome cleaned reads were mapped to this combined dataset using bowtie2 (--end-to-end). CoverM (--genome, v.0.6.1) was utilized to generate TPM counts for the algal endosymbionts.

*Species Diversity*

Alpha diversity was assessed using Shannon diversity, Simpson evenness index, and Chao1 richness (scikit-bio v0.6.3 using function alpha) metrics. A Shapiro-Wilk test was utilized to assess the normality of the data. Due to the data not being normally distributed, the non-parametric Kruskal Wallis test was performed to assess significant shifts between health conditions. Furthermore, the Mann Whitney U test was used to assess pairwise community shifts across the three health conditions. Beta diversity between the samples was assessed using Aitchison distance, by first performing a centered log ratio transformation and subsequently generating a dissimilarity matrix (scipy.spatial.distance: pdist, squareform, and skbio.stats.composition: clr) [27]. To further assess the sample clustering profile, we used the same dissimilarity matrix in a non-metric multidimensional scaling (NMDS) analysis using sklearn.manifold MDS (scikit-learn v1.6.1). We further performed a PERMANOVA analysis (999 permutations) and PERMDISP to assess the effect of sample variation.

*Differential Abundance Analysis*

Numerous tools exist to assess differential abundance in metagenomic data. Non-normally distributed data, paired with its heterogenous nature and the large number of zero values made it challenging to select a single best tool [28]. Conservative tools like ANCOM-BC [29] and Aldex2 [30] have been championed as the most reliable for microbiome analyses. However, these tools often require many samples per group, a minimum of *n* = 10 and *n* = 8 (respectively), whereas our study only had *n* = 5. Furthermore, studies have also assessed the utilization of tools designed for RNA-seq data analysis, like edgeR [31] and limma-voom, however these are not recommended due to high false positive rates [32, 33] Another such tool recommended for metagenomic analysis is DESeq2 (or pyDESeq2 v0.4.8) [34] which can run with fewer samples per group (*n* = 3). Normalization factors were calculated (fit_size_function()), followed by genewise dispersions (fit_genewise_dispersion()), fitting dispersion trend coefficients (fit_dispersion_trend()), lastly log FC was calculated (fit_LFC). Whereas DESeq2 was observed to still yield some false positive results, in this study we paired DESeq2 results with that of a basic TPM-based foldchange assessment (log_2_[DS or AH TPM] – log_2_[HC TPM]; see below) and visualization of the data to confirm the trends observed in the data.

*Verifying vMAGs and putative host identification*

A common challenge when isolating vMAGs in metagenomic data is the mis-annotation of host contigs as viral. To confirm viral origin and putative hosts, proteins from vMAGs associated with SCTLD were assessed using phylogenetics. DIAMOND blastp (v2.1.10.164; --ultra-sensitive --max-target-seqs 0 --evalue 1e-05) [35] was used to search the vMAG proteins against two databases, nr (2022_07) and IMG/VR (v4.1) [36]. The hits returned by this analysis were too voluminous for phylogenetic analysis (computationally intractable or too large to visualize), necessitating the application of downsampling. For each query protein, hits to the **nr** database were downsampled, retaining only the **top hit for each taxonomic class (i.e., for each taxonomic class in the resulting hits, just the top hit with the lowest *e*-value was retained)**. For hits to the IMG/VR database, only the top 50 hits (based on bitscore) were selected for phylogenetic analysis (due to the absence of reliable taxonomy information for many of the sequences in IMG/VR). For each query, the protein sequences of the hits which remained after downsampling were combined, along with the query protein itself, into a file for downstream analysis. For each of the resulting protein files, MAFFT (v7.520; --auto) [37] was used to generate an alignment and trimAl (v1.5.rev0; --automated1) [38] was used to trim poor quality sequences and positions. Maximum Likelihood (ML) phylogenetic trees were generated for each trimmed alignment using iqtree (v2.3.6; -m TEST -bb 1000) [39]. The placement of the query protein in each tree was assessed relative to all viral and non-viral proteins to verify true viral origin and potentially identify putative hosts. The function of each viral protein was assessed using **GhostKOALA (v3.1.0) [40]**.

*Assessing other SCTLD metagenomic studies*

Reads were downloaded from NCBI using Kingfisher (v0.4.1) [41]. Reads were assessed for quality using fastqc (v 0.12.1) [42]; this analysis confirmed that the reads were of high quality, with only small amounts of adapter contamination, and thus did not require additional processing before mapping. Obtained reads were mapped to all 264 MAGs generated in this study using bowtie2 (--local) (v2.5.4) [43], with the coverage of each MAG in each sample calculated using CoverM (--genome) (v.0.6.1) [44]. Due to only a small number of MAGs having mapped reads, we were unable to use relative abundance or pyDESeq2 to analyze the results. Furthermore, a comparison between HC, AH, and DS samples was deemed unreliable due the putative presence of asymptomatic colonies, and the unknown outcomes of the healthy colonies (infected later or disease resistant). Therefore, we only performed a presence/absence analysis using this data: i.e., a MAG was considered present in a species if it had a TPM > 1.0 in at least 5 samples (out of the 58 total).

**Results and Discussion**

*General metagenome analysis*

Metagenomic short read sequencing produced 249-321 million read pairs per sample, which constituted 74.8-113.5 Gbp of data per sample [**Supplemental Table 7**]. As expected, read deduplication, quality filtering, and mapping to the preliminary *D. labyrinthiformis* genome generated from the first round of sequencing significantly reduced the total amount of data for binning, with many samples reducing in size by >90%. This is an expected result because most of the read data will be of host provenance and is removed during the mapping stage. Removal of this data improves both the runtime and accuracy of metagenomic workflows. After read processing, binning, and quality assessment we obtained a total of 264 MAGs which exceeded the minimum MAG quality cutoffs used for each taxonomic domain (**See Supplemental Text**). Of the 264 MAGs, two were eukaryotic, a sponge and a diatom, which had BUSCO completeness scores of 23.9% and 11% (respectively). Owing to their low completeness these eukaryotes were not included in further downstream analysis. A total of 85 MAGs were prokaryotic (pMAGs), 23 were classified as “Complete” (completeness > 90% and contamination < 5%, as assessed by CheckM), 39 were classified as “High-quality” (completeness > 70% and contamination < 10%), and 23 were classified as “Medium-quality” (completeness > 50% and contamination < 10%) [**Supplemental Table 1**]. Additionally, we identified 36 plasmid MAGs, with a geNOMAD plasmid score > 0.85 and a false discovery rate (FDR) < 0.05, and 141 viral MAGs (vMAGs), with a geNOMAD score > 0.90 and an FDR < 0.05. Of the 141 vMAGs, 109 were classified as Caudoviricetes, which are widely distributed bacteriophages [**Supplemental Table 2**].

*Algal endosymbiont composition*

After mapping the cleaned metagenomic data against a combined database of all available endosymbiont genomes it was observed that four species were detected across the collected samples: *Breviolum minutum*, *Cladocopium goreaui*, *Durusdinium trenchii* (CCMP2556 and SCF082_1), and *Symbiodinium necroappetens* [**Supplemental Figure 6**, **Supplemental Table 8**]. In the HC samples we observed a mixed infection of *D. trenchii* (0.21 ± 0.20) and *B. minutum* (0.57 ± 0.46). One sample also had the endosymbiont *C. goreaui* at a relative abundance of 0.06. The presence of these endosymbionts is consistent with previous findings of endosymbiont populations in the Caribbean [45]. *D. trenchii and B. minutum* are also highly prevalent in infected colonies (AH and DS), however, the opportunistic *S. necroappetens*, which has been previously isolated from Yellow Band Disease infected corals (active lesions) and bleached colonies in the Caribbean is also present [46]. In the AH samples, IC5_AH had no reads mapped to any of the endosymbiont genomes, however, in the remaining samples, *S. necroappetens* (0.44 ± 0.28) was the most abundant, followed by *D. trenchii* (0.20 ± 0.17) and *B. minutum* (0.00 ± 0.06). Lastly in the DS samples, *D. trenchii* (0.33 ± 0.18) was the most abundant, followed by *S. necroappetens* (0.11 ± 0.37) and *B. minutum* (0.05 ± 0.09).

*Virome phylum level analysis*

Phylum-level taxonomic analysis of the virome was performed to assess the overall shifts in the viral community between the different health conditions. In HC samples, Uroviricota was the most abundant phylum (0.5094 ± 0.2083), followed by Nucleocytoviricota (0.2054 ± 0.1410). In AH samples, the most abundant phylum shifts to Nucleocytoviricota (0.4496 ± 0.1002); this shift was large enough to reach statistical significance (*p*-value = 0.035). A decrease of Uroviricota (0.3636 ± 0.0429) was observed in the AH samples, although it was not statistically significant. Lastly in DS samples, the opposite pattern was observed: i.e., Uroviricota was the most abundant (0.4083 ± 0.0432), followed by Nucleocytoviricota (0.3762 ± 0.1176). Log₂TPM fold change values for each of the vMAGs were also examined to capture general trends in the virome [**Supplemental Figure 3**]. We observe an increased abundance of 18 vMAGs in AH samples and 33 vMAGs in DS samples (all compared to HC samples), of which 16 were shared. Furthermore, 97 vMAGs have a decreased absence in AH samples and 95 vMAGs in DS samples.

These results show that there is a notable shift in viral community composition across health conditions. Specifically, HC samples were dominated by Uroviricota, particularly Caudoviricetes, which are bacteriophages typically associated with environmental microbiomes. In contrast, AH samples were dominated by Nucleocytoviricota, a phylum of double-stranded DNA viruses. Interestingly, Uroviricota abundance showed a partial resurgence in DS samples, where lesions were already established, and significant tissue loss had occurred. The decline of Uroviricota in AH samples aligns with shifts in the prokaryotic microbiome, their primary host, suggesting a tight coupling between viral and bacterial community dynamics. The subsequent resurgence in DS samples likely results from bacteriophage infection of newly colonizing bacteria. Moreover, the high diversity of vMAGs in healthy colonies makes it challenging to attribute a biologically significant contribution to any single vMAG. However, general virome trends can be utilized, and show that a majority of vMAGs decrease in relative abundance in samples from infected colonies (IC).

*Prokaryote abundance results*

It is important to note that in one infected colony, samples IC_1 (AH) and IC_1 (DS), we observe low Leptospirales and Gammaproteobacteria abundance. All other infected colonies showed an increased prevalence of these pMAGs, suggesting an alternative opportunistic pathogen was present in samples N35 and N33. This idea is corroborated by other SCTLD studies that have identified different putatively pathogenic bacteria associated with infection. These other studies have found Rhodobacterales, Rhizobiales*,* and *Vibrio* spp. [47, 48] to be enriched in diseased corals, however, our analysis did not identify these bacteria, suggesting that the secondary infection agent may differ across coral species and locations.

*Microbiome profile shifts under infection*

An initial relative abundance analysis of the microbiome was performed at the “class” taxonomic level [**Supplemental Table 1**]. In HC samples, Alphaproteobacteria was the most abundant class (0.3918 ± 0.2209), followed by Gammaproteobacteria (0.1461 ± 0.2585) [**Figure 2A**]. In AH samples, Gammaproteobacteria was the most abundant (0.3432 ± 0.3603) and Leptospirales was the second most abundant (0.3047 ± 0.2873), and Alphaproteobacteria was the third (0.0159 ± 0. 1945). In DS samples, Gammaproteobacteria was the most abundant (0.4207 ± 0.2397), followed by Alphaproteobacteria (0.1538 ± 0.2317) and *Leptospirae* (0.0453 ± 0.2444). The increase in the class *Leptospirae* was statistically significant in both AH and DS samples compared to the HC samples (*p*-value = 0.035 for both).

We assessed **differential abundance** of individual pMAGs using DESeq2 (|log₂FC| > 2.0 and an adjusted *p*-value < 0.05). Out of the 85 pMAGs, three showed a significant increase in infected colonies, in both AH and DS samples. In addition, seven pMAGs showed significantly decreased abundance in AH samples, and eight in DS samples. Only one pMAG was not decreased in AH but decreased in DS. To evaluate broader patterns beyond statistical significance, we calculated log₂TPM-based fold changes. Although this analysis did not assess statistical significance, it revealed that 16 pMAGs had increased abundance in infected colonies, whereas 69 pMAGs showed decreased abundance, suggesting a general trend of microbial decline across infected colonies (in either HC_vs_DS or HC_vs_AH) [**Figure 2B**]. To interpret biological significance of these shifts, we looked at the relative abundance of each of the statistically significant pMAGs. In infected colonies, three pMAGs had an increased abundance: MAG_prokaryotic_02, MAG_prokaryotic_01, and MAG_prokaryotic_45. In HC samples, we observe that the relative abundance of these MAGs was 0.000127 ± 0.000117, 0.000003 ± 0.000003, and 0.000264 ± 0.000103, respectively. In AH samples, an increase in relative abundance (compared to HC samples) was observed, 0.304739 ± 0.287329, 0.177501 ± 0.240849, and 0.009025 ± 0.129915. In the DS samples, the relative abundance was higher than in the HC samples but lower than the AH samples, 0.045344 ± 0.244371, 0.055040 ± 0.158030, and 0.015674 ± 0.062248. pMAGs that had a decreased abundance, i.e., abundant in healthy but less abundant in infected colonies, were more challenging to assess due to the high microbiome richness and diversity in the HC samples. That is, pMAGs in healthy colonies tended to be specific to a single colony and not broadly shared across other colonies, confounding statistical analysis. Given these considerations, pMAGs were considered of biological interest if they were differentially abundant and had a median relative abundance > 0.1 in at least one health condition. As a result, MAG_prokaryotic_02 and MAG_prokaryotic_01 were selected for downstream analyses.

*vMAG phylogenetic analysis*

Of the 41 proteins from the five SCTLD-associated vMAGs, we were able to generate phylogenetic trees for 25 **[Supplemental File 1]**. vMAG058 had no genes with sufficient hits to nr or IMGV for phylogenetic reconstruction and thus could not be assessed by this analysis. For vMAG001 (geNOMAD taxonomy: Adintoviridae), 11/15 genes had phylogenies constructed, all of which contained hydra or coral sequences intermixed with known viral and bacterial sequences [**Supplemental Table 5**]. Furthermore, 10 of the trees contained hydra adintovirus sequences and 2 contained *Stylophora* adintovirus sequences, supporting vMAG001 as putatively host infecting. For vMAG043 (geNOMAD taxonomy: Imitevirales), 3/7 genes had phylogenies constructed. Two of the constructed phylogenies contained the coral virus *Rhodactis* coral adintovirus and the coral *Acropora millepora*. For vMAG055 (geNOMAD taxonomy: Megaviricetes), 5/9 genes had phylogenies constructed. None of the phylogenies contained any corals, however 2 trees contained coral endosymbiont sequences from *Symbiodinium microadriaticum* and *Symbiodinium natans*. Lastly, for vMAG060 (geNOMAD taxonomy: Caudoviricites), 7/8 genes had phylogenies constructed. All seven trees contained either corals or hydras sequences, and only one gene contained *Corynactis* coral adintovirus, suggesting that vMAG060 may be a putative host infecting virus. However, across all vMAGs discussed, the majority of trees also included other bacteria, making it difficult to unambiguously identify the host.
