## Supplementary figures and images for "Stony coral tissue loss disease (SCTLD) destabilizes the coral microbiome"

### Supplemental Figure 1

# HC_1

# HC_2

**June 12, 2024**

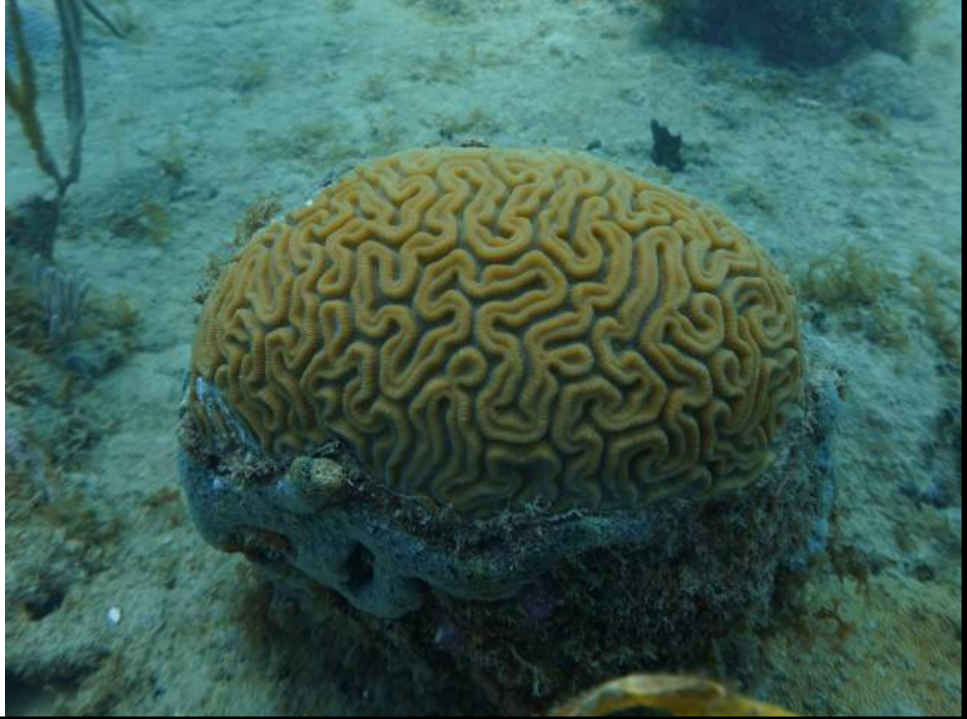

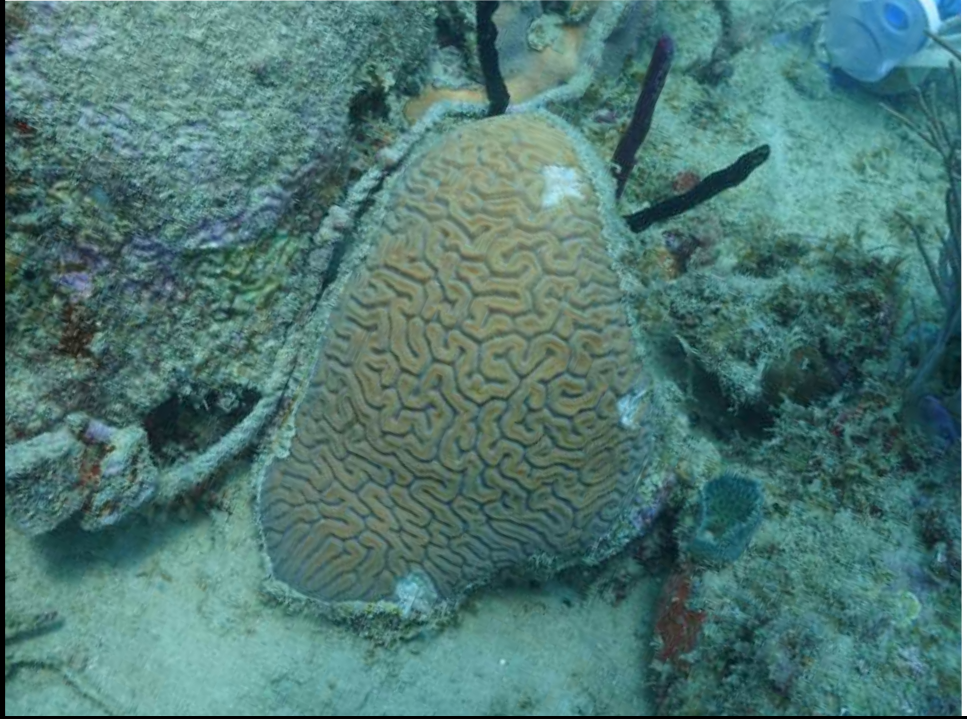

**March 28, 2025**

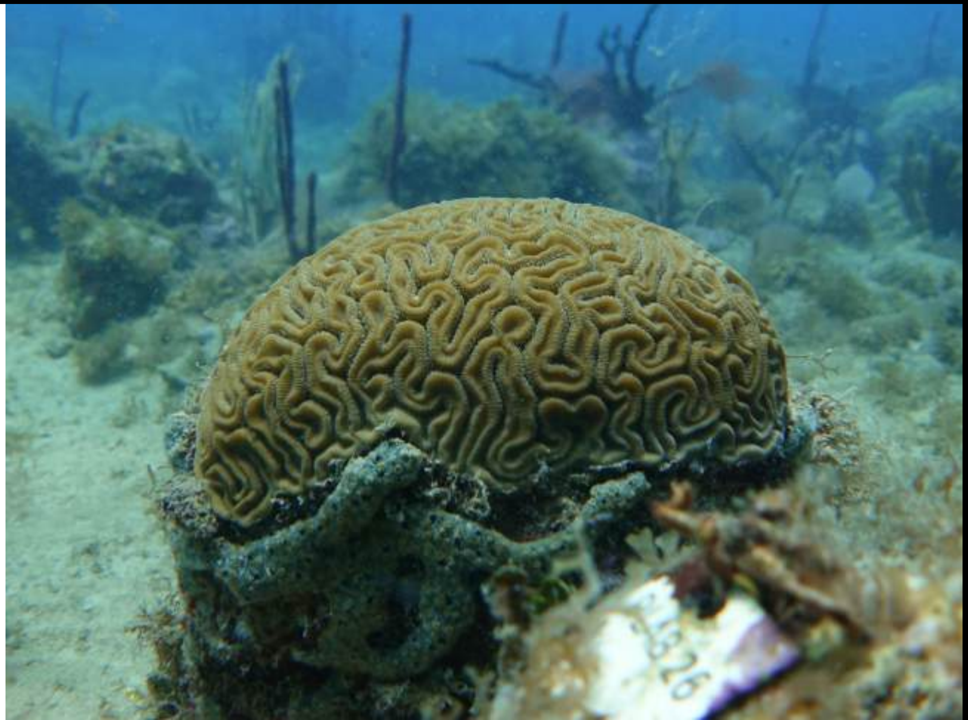

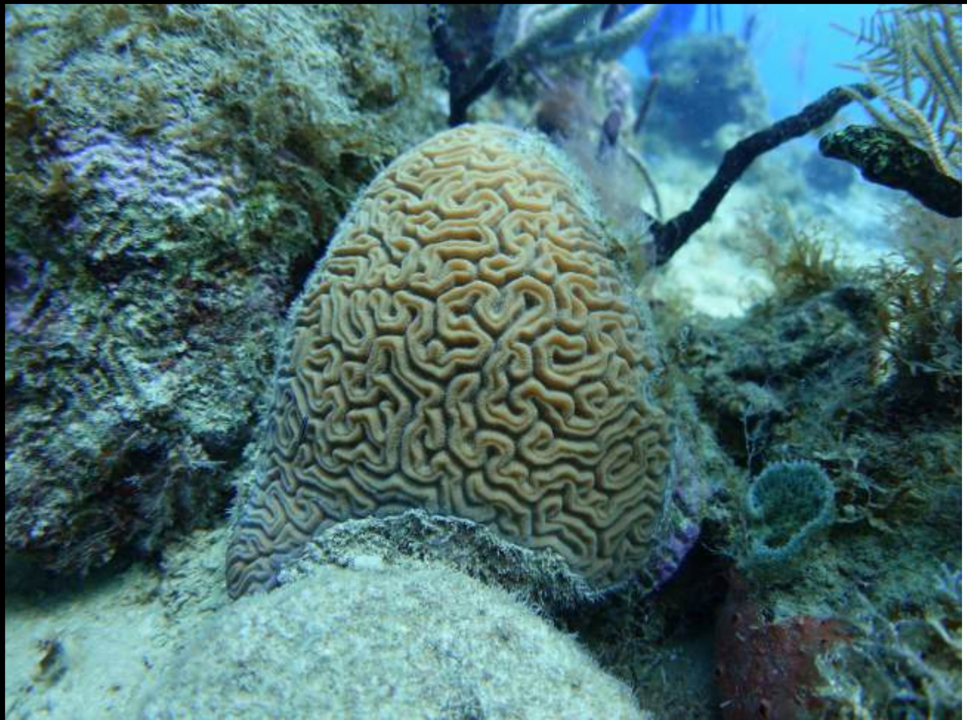

### Supplemental Figure 2

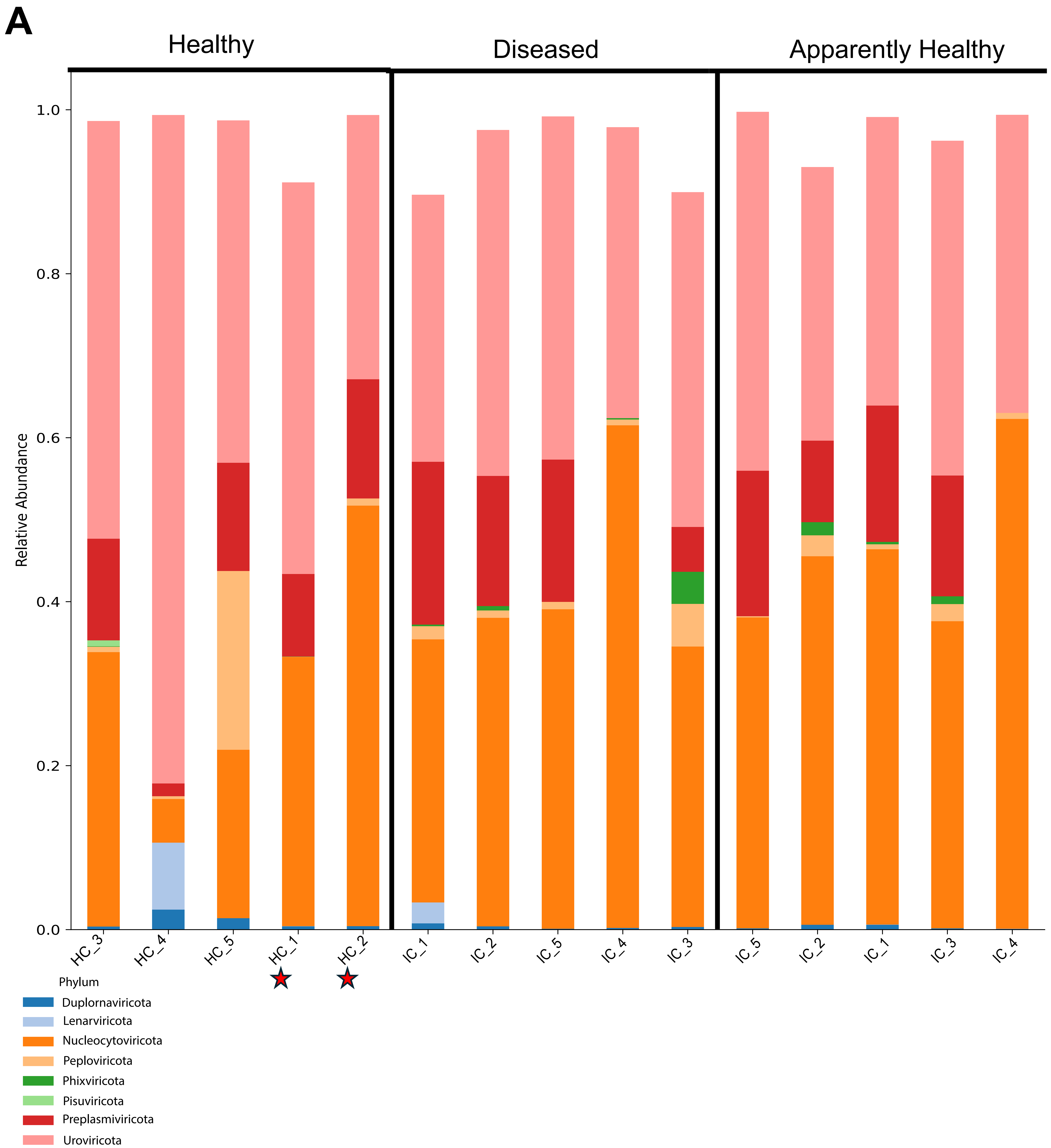

### Supplemental Figure 3

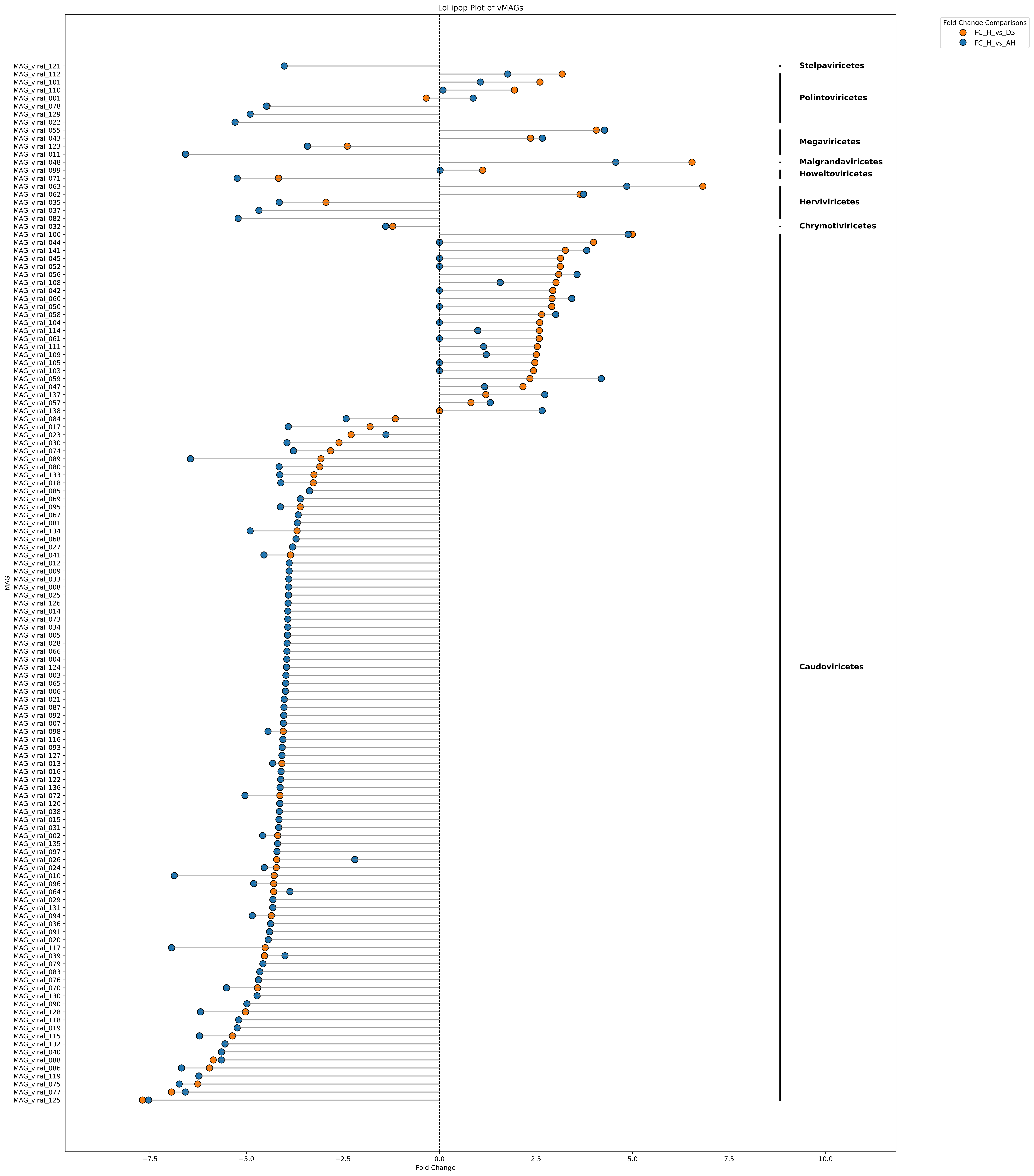

### Supplemental Figure 4

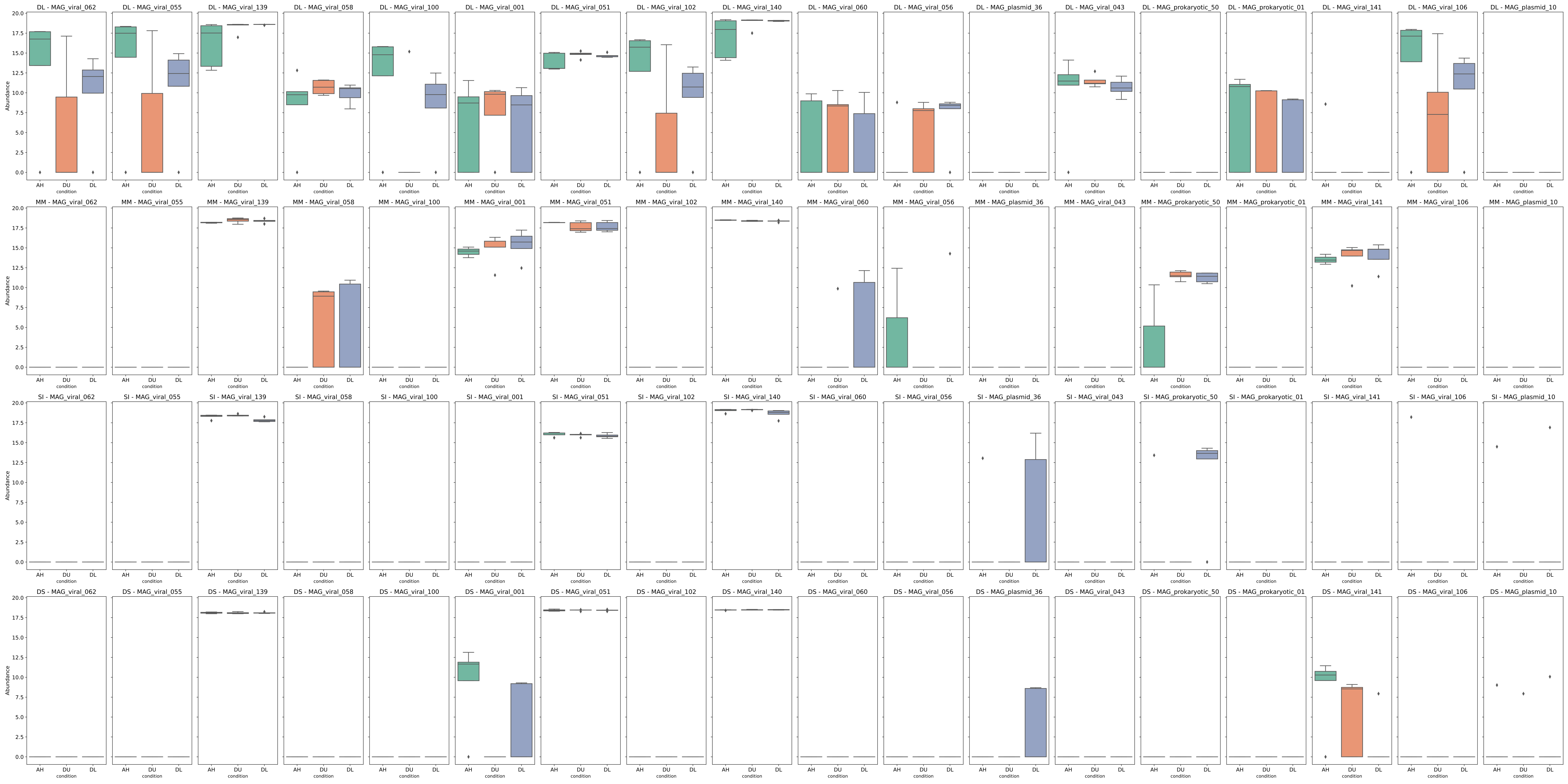

### Supplemental Figure 5

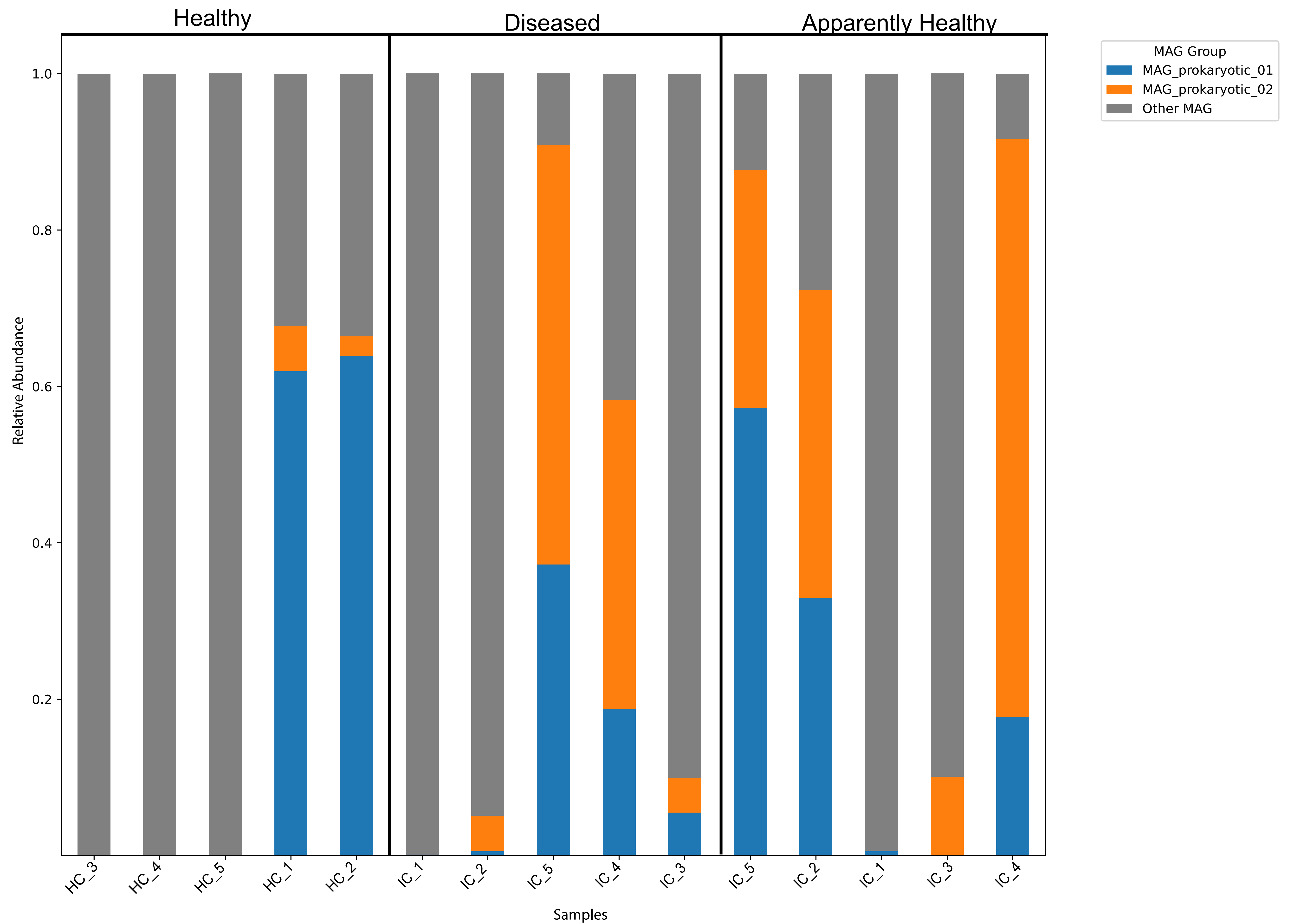

### Supplemental Figure 6

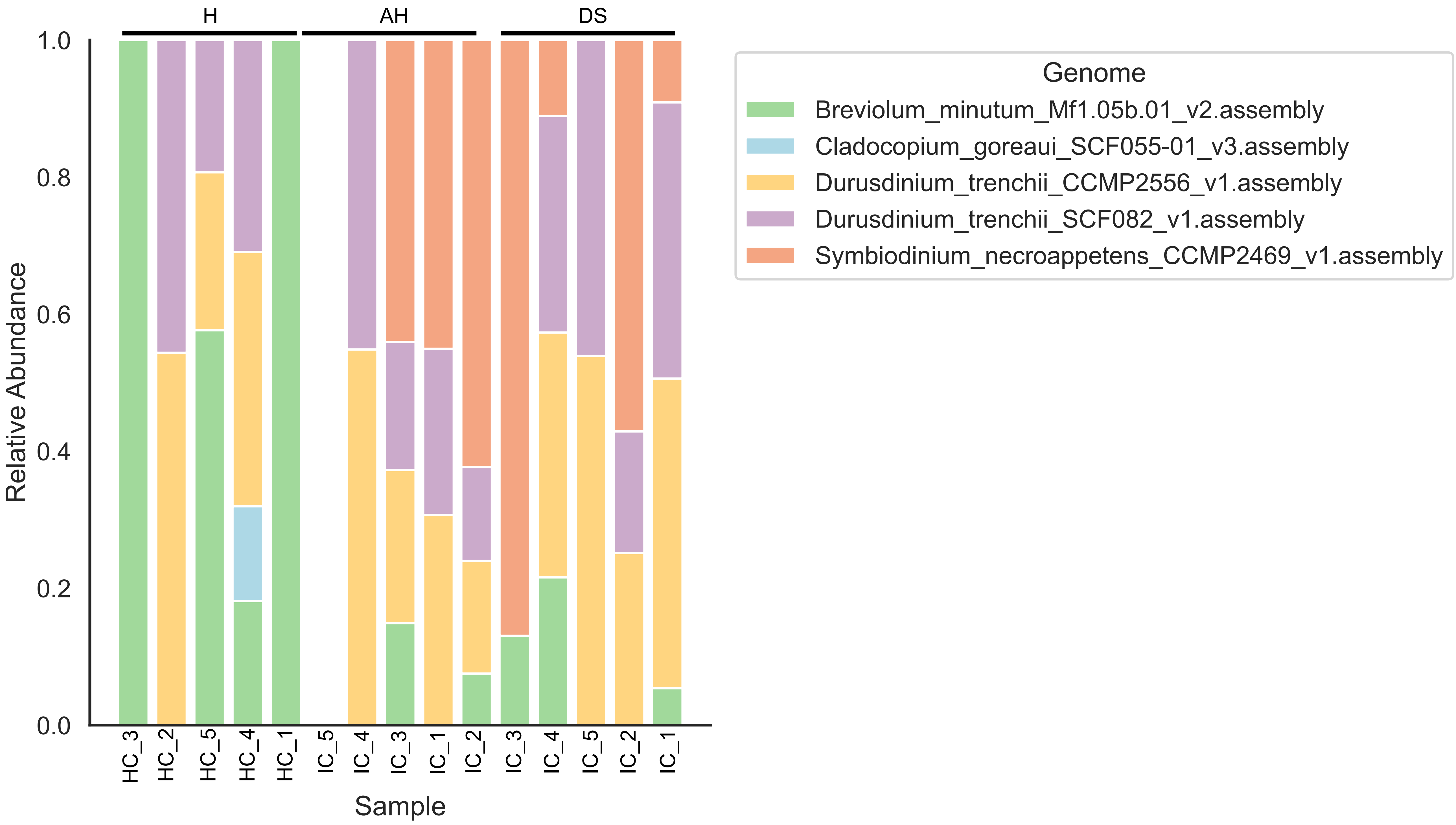
