## Supplemental File 1 for "Stony coral tissue loss disease (SCTLD) destabilizes the coral microbiome"

| viral_protein | Model |
|---|---|
| MAG_viral_001-00000001_1 | LG+F+I+G4 chosen according to BIC |
| MAG_viral_001-00000002_1 | LG+F+I+G4 chosen according to BIC |
| MAG_viral_001-00000002_10 | LG+F+G4 chosen according to BIC |
| MAG_viral_001-00000002_11 | LG+I+G4 chosen according to BIC |
| MAG_viral_001-00000002_12 | LG+I+G4 chosen according to BIC |
| MAG_viral_001-00000002_14 | VT+G4 chosen according to BIC |
| MAG_viral_001-00000002_2 | LG+F+G4 chosen according to BIC |
| MAG_viral_001-00000002_4 | LG+F+I+G4 chosen according to BIC |
| MAG_viral_001-00000002_5 | LG+G4 chosen according to BIC |
| MAG_viral_001-00000002_6 | LG+I+G4 chosen according to BIC |
| MAG_viral_001-00000002_8 | VT chosen according to BIC |
| MAG_viral_001-00000002_9 | LG+G4 chosen according to BIC |
| MAG_viral_043-00000001_2 | LG+G4 chosen according to BIC |
| MAG_viral_043-00000001_3 | LG+I+G4 chosen according to BIC |
| MAG_viral_043-00000001_4 | LG+I+G4 chosen according to BIC |
| MAG_viral_055-00000001_1 | LG+I chosen according to BIC |
| MAG_viral_055-00000001_3 | LG+G4 chosen according to BIC |
| MAG_viral_055-00000001_4 | LG+G4 chosen according to BIC |
| MAG_viral_055-00000001_5 | LG+G4 chosen according to BIC |
| MAG_viral_055-00000001_6 | VT+G4 chosen according to BIC |
| MAG_viral_055-00000001_7 | LG+I+G4 chosen according to BIC |
| MAG_viral_055-00000001_8 | LG+F+I+G4 chosen according to BIC |
| MAG_viral_058-00000001_1 | LG chosen according to BIC |
| MAG_viral_060-00000001_1 | LG+G4 chosen according to BIC |
| MAG_viral_060-00000001_2 | JTT+G4 chosen according to BIC |
| MAG_viral_060-00000001_3 | LG+G4 chosen according to BIC |
| MAG_viral_060-00000001_4 | LG+I+G4 chosen according to BIC |
| MAG_viral_060-00000001_5 | LG+G4 chosen according to BIC |
| MAG_viral_060-00000001_6 | LG chosen according to BIC |
| MAG_viral_060-00000001_7 | LG+G4 chosen according to BIC |
| MAG_viral_060-00000001_8 | LG+I chosen according to BIC |

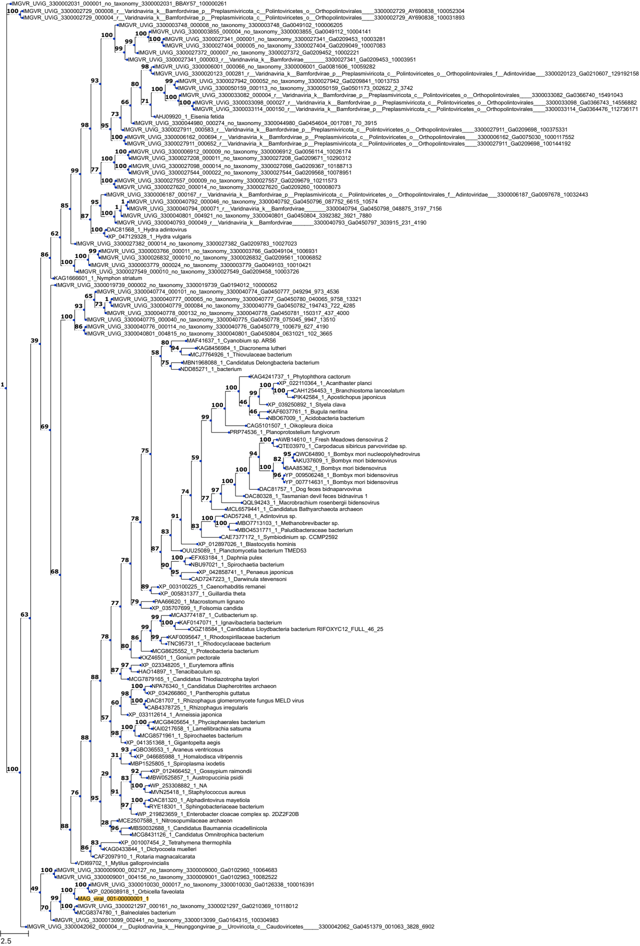
IMGVR_UViG_3300002031_000001_no_taxonomy_3300002031_BBAY57_100000261
IMGVR_UViG_3300002729_000008_r__Varidnaviria_k__Bamfordvirae_p__Preplasmiviricota_c__Polintoviricetes_o__Orthopolintovirales____3300002729_AY690838_100052304
IMGVR_UViG_3300002729_000004_r__Varidnaviria_k__Bamfordvirae_p__Preplasmiviricota_c__Polintoviricetes_o__Orthopolintovirales____3300002729_AY690838_100031893
IMGVR_UViG_3300003748_000008_no_taxonomy_3300003748_Ga0049102_100006205
IMGVR_UViG_3300003855_000004_no_taxonomy_3300003855_Ga0049112_10004141
IMGVR_UViG_3300027341_000001_no_taxonomy_3300027341_Ga0209453_10003281
IMGVR_UViG_3300027404_000005_no_taxonomy_3300027404_Ga0209049_10007083
IMGVR_UViG_3300027372_000007_no_taxonomy_3300027372_Ga0209452_10002221
IMGVR_UViG_3300027341_000003_r__Varidnaviria_k__Bamfordvirae________3300027341_Ga0209453_10003951
IMGVR_UViG_3300006001_000066_no_taxonomy_3300006001_Ga0081606_10059282
IMGVR_UViG_3300020123_000281_r__Varidnaviria_k__Bamfordvirae_p__Preplasmiviricota_c__Polintoviricetes_o__Orthopolintovirales_f__Adintoviridae___3300020123_Ga0210607_129192158
IMGVR_UViG_3300027942_000052_no_taxonomy_3300027942_Ga0209841_10013753
IMGVR_UViG_3300050159_000113_no_taxonomy_3300050159_Ga0501173_002622_2_3742
IMGVR_UViG_3300033082_000004_r__Varidnaviria_k__Bamfordvirae_p__Preplasmiviricota_c__Polintoviricetes_o__Orthopolintovirales____3300033082_Ga0366740_15491043
IMGVR_UViG_3300033098_000027_r__Varidnaviria_k__Bamfordvirae_p__Preplasmiviricota_c__Polintoviricetes_o__Orthopolintovirales____3300033098_Ga0366743_14556882
IMGVR_UViG_3300033114_000150_r__Varidnaviria_k__Bamfordvirae_p__Preplasmiviricota_c__Polintoviricetes_o__Orthopolintovirales____3300033114_Ga0364476_112736171
AHJ09920_1_Eisenia fetida
IMGVR_UViG_3300044980_000274_no_taxonomy_3300044980_Ga0454604_0017081_70_3915
IMGVR_UViG_3300027911_000583_r__Varidnaviria_k__Bamfordvirae_p__Preplasmiviricota_c__Polintoviricetes_o__Orthopolintovirales____3300027911_Ga0209698_100375331
IMGVR_UViG_3300006162_000694_r__Varidnaviria_k__Bamfordvirae_p__Preplasmiviricota_c__Polintoviricetes_o__Orthopolintovirales____3300006162_Ga0075030_1000117552
IMGVR_UViG_3300027911_000652_r__Varidnaviria_k__Bamfordvirae_p__Preplasmiviricota_c__Polintoviricetes_o__Orthopolintovirales____3300027911_Ga0209698_100144192
IMGVR_UViG_3300006912_000009_no_taxonomy_3300006912_Ga0056114_10026174
IMGVR_UViG_3300027208_000011_no_taxonomy_3300027208_Ga0209671_10290312
IMGVR_UViG_3300027098_000014_no_taxonomy_3300027098_Ga0209367_10188713
IMGVR_UViG_3300027544_000022_no_taxonomy_3300027544_Ga0209568_10078951
IMGVR_UViG_3300027557_000009_no_taxonomy_3300027557_Ga0209679_10211573
IMGVR_UViG_3300027620_000014_no_taxonomy_3300027620_Ga0209260_100008073
IMGVR_UViG_3300006187_000167_r__Varidnaviria_k__Bamfordvirae_p__Preplasmiviricota_c__Polintoviricetes_o__Orthopolintovirales_f__Adintoviridae___3300006187_Ga0097678_10032443
IMGVR_UViG_3300040792_000046_no_taxonomy_3300040792_Ga0450796_087752_6615_10574
IMGVR_UViG_3300040794_000071_r__Varidnaviria_k__Bamfordvirae________3300040794_Ga0450798_048875_3197_7156
IMGVR_UViG_3300040801_004921_no_taxonomy_3300040801_Ga0450804_3392382_3921_7880
IMGVR_UViG_3300040793_000049_r__Varidnaviria_k__Bamfordvirae________3300040793_Ga0450797_303915_231_4190
DAC81568_1_Hydra adintovirus
XP_047129328_1_Hydra vulgaris
IMGVR_UViG_3300027382_000014_no_taxonomy_3300027382_Ga0209783_10027023
IMGVR_UViG_3300003766_000011_no_taxonomy_3300003766_Ga0049104_1006931
IMGVR_UViG_3300026832_000010_no_taxonomy_3300026832_Ga0209561_10006852
IMGVR_UViG_3300003779_000024_no_taxonomy_3300003779_Ga0049103_10010421
IMGVR_UViG_3300027549_000010_no_taxonomy_3300027549_Ga0209458_10003726
KAG1666601_1_Nymphon striatum
IMGVR_UViG_3300019739_000002_no_taxonomy_3300019739_Ga0194012_10000052
IMGVR_UViG_3300040774_000101_no_taxonomy_3300040774_Ga0450777_049294_973_4536
IMGVR_UViG_3300040777_000065_no_taxonomy_3300040777_Ga0450780_040065_9758_13321
IMGVR_UViG_3300040779_000084_no_taxonomy_3300040779_Ga0450782_194743_722_4285
IMGVR_UViG_3300040778_000132_no_taxonomy_3300040778_Ga0450781_150317_437_4000
IMGVR_UViG_3300040775_000040_no_taxonomy_3300040775_Ga0450778_075045_9947_13510
IMGVR_UViG_3300040776_000114_no_taxonomy_3300040776_Ga0450779_100679_627_4190
IMGVR_UViG_3300040801_004815_no_taxonomy_3300040801_Ga0450804_0631021_102_3665
MAF41637_1_Cyanobium sp. ARS6
KAG8456984_1_Diacronema lutheri
MCJ7764926_1_Thiovulaceae bacterium
MBN1968088_1_Candidatus Delongbacteria bacterium
NDD85271_1_bacterium
KAG4241737_1_Phytophthora cactorum
XP_022110364_1_Acanthaster planci
CAH1254453_1_Branchiostoma lanceolatum
PIK42584_1_Apostichopus japonicus
XP_039250892_1_Styela clava
KAF6037761_1_Bugula neritina
NBO67009_1_Acidobacteria bacterium
CAG5101507_1_Oikopleura dioica
PRP74536_1_Planoprotostelium fungivorum
AWB14610_1_Fresh Meadows densovirus 2
QTE03970_1_Carpodacus sibiricus parvoviridae sp.
QWC64890_1_Bombyx mori nucleopolyhedrovirus
AKU37609_1_Bombyx mori bidensovirus
BAA85362_1_Bombyx mori bidensovirus
YP_009506248_1_Bombyx mori bidensovirus
YP_007714631_1_Bombyx mori bidensovirus
DAC81757_1_Dog feces bidnaparvovirus
DAC80328_1_Tasmanian devil feces bidnavirus 1
QQL94243_1_Macrobrachium rosenbergii bidensovirus
MCL6579441_1_Candidatus Bathyarchaeota archaeon
DAD57248_1_Adintovirus sp.
MBO7713103_1_Methanobrevibacter sp.
MBO4531771_1_Paludibacteraceae bacterium
CAE7377172_1_Symbiodinium sp. CCMP2592
XP_012897026_1_Blastocystis hominis
OUU25089_1_Planctomycetia bacterium TMED53
EFX63184_1_Daphnia pulex
NBU97021_1_Spirochaetia bacterium
XP_042858741_1_Penaeus japonicus
CAD7247223_1_Darwinula stevensoni
XP_003100225_1_Caenorhabditis remanei
XP_005831377_1_Guillardia theta
PAA66620_1_Macrostomum lignano
XP_035707699_1_Folsomia candida
MCA3774187_1_Cutibacterium sp.
KAF0147071_1_Ignavibacteria bacterium
OGZ18584_1_Candidatus Lloydbacteria bacterium RIFOXYC12_FULL_46_25
KAF0095647_1_Rhodospirillaceae bacterium
TNC95731_1_Rhodocyclaceae bacterium
MCG8625552_1_Proteobacteria bacterium
KXZ46501_1_Gonium pectorale
XP_023348205_1_Eurytemora affinis
HAO14897_1_Tenacibaculum sp.
MCG7879165_1_Candidatus Thiodiazotropha taylori
NPA76340_1_Candidatus Diapherotrites archaeon
XP_034266860_1_Pantherophis guttatus
DAC81707_1_Rhizophagus glomeromycete fungus MELD virus
CAB4378725_1_Rhizophagus irregularis
XP_033112614_1_Anneissia japonica
MCG8405654_1_Phycisphaerales bacterium
KAI0217658_1_Lamellibrachia satsuma
MCG8571961_1_Spirochaetes bacterium
XP_041351368_1_Gigantopelta aegis
GBO36553_1_Araneus ventricosus
XP_046685988_1_Homalodisca vitripennis
MBP1525805_1_Spiroplasma ixodetis
XP_012466452_1_Gossypium raimondii
MBW0525857_1_Austropuccinia psidii
WP_253308882_1_NA
MVN25418_1_Staphylococcus aureus
DAC81320_1_Alphadintovirus mayetiola
RYE18301_1_Sphingobacteriaceae bacterium
WP_219823659_1_Enterobacter cloacae complex sp. 2DZ2F20B
MCE2507588_1_Nitrosopumilaceae archaeon
MBS0032688_1_Candidatus Baumannia cicadellinicola
MCG8431126_1_Candidatus Omnitrophica bacterium
XP_001007454_2_Tetrahymena thermophila
KAG0433844_1_Dictyocoela muelleri
CAF2097910_1_Rotaria magnacalcarata
VDI69702_1_Mytilus galloprovincialis
IMGVR_UViG_3300009000_002127_no_taxonomy_3300009000_Ga0102960_10064683
IMGVR_UViG_3300009001_004156_no_taxonomy_3300009001_Ga0102963_10082522
IMGVR_UViG_3300010030_000017_no_taxonomy_3300010030_Ga0126338_100016391
XP_020608918_1_Orbicella faveolata
MAG_viral_001-00000001_1
IMGVR_UViG_3300021297_000161_no_taxonomy_3300021297_Ga0210369_10118012
MCG8374780_1_Balneolales bacterium
IMGVR_UViG_3300013099_002441_no_taxonomy_3300013099_Ga0164315_100304983
IMGVR_UViG_3300042062_000004_r__Duplodnaviria_k__Heunggongvirae_p__Uroviricota_c__Caudoviricetes______3300042062_Ga0451379_001063_3828_6902
2.5

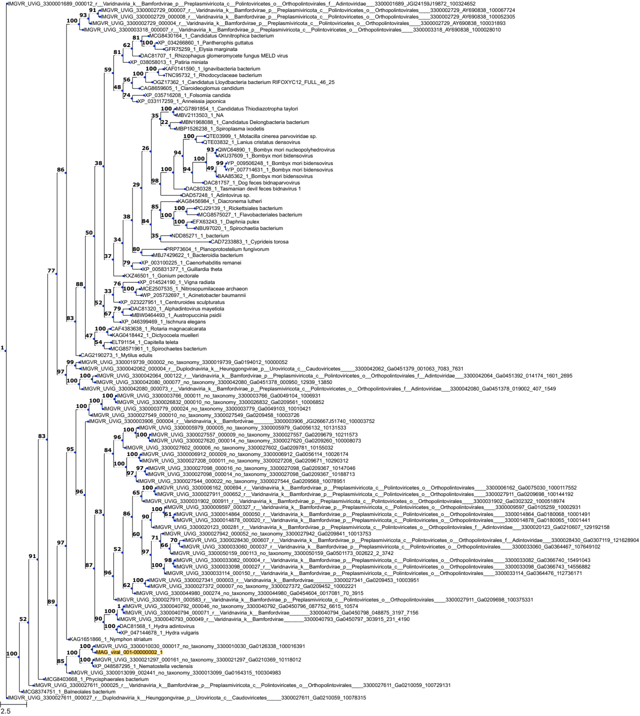

IMGVR_UViG_3300001689_000012_r__Varidnaviria_k__Bamfordvirae_p__Preplasmiviricota_c__Polintoviricetes_o__Orthopolintovirales_f__Adintoviridae___3300001689_JGI24159J19872_100324652
IMGVR_UViG_3300002729_000007_r__Varidnaviria_k__Bamfordvirae_p__Preplasmiviricota_c__Polintoviricetes_o__Orthopolintovirales____3300002729_AY690838_100067724
IMGVR_UViG_3300002729_000008_r__Varidnaviria_k__Bamfordvirae_p__Preplasmiviricota_c__Polintoviricetes_o__Orthopolintovirales____3300002729_AY690838_100052305
IMGVR_UViG_3300002729_000004_r__Varidnaviria_k__Bamfordvirae_p__Preplasmiviricota_c__Polintoviricetes_o__Orthopolintovirales____3300002729_AY690838_100031893
IMGVR_UViG_3300003318_000007_r__Varidnaviria_k__Bamfordvirae_p__Preplasmiviricota_c__Polintoviricetes_o__Orthopolintovirales____3300003318_AY690838_1000028010
MCG8430164_1_Candidatus Omnitrophica bacterium
XP_034266860_1_Pantherophis guttatus
GFR75259_1_Elysia marginata
DAC81707_1_Rhizophagus glomeromycete fungus MELD virus
XP_038058013_1_Patiria miniata
KAF0141590_1_Ignavibacteria bacterium
TNC95732_1_Rhodocyclaceae bacterium
OGZ17362_1_Candidatus Lloydbacteria bacterium RIFOXYC12_FULL_46_25
CAG8659605_1_Claroideoglomus candidum
XP_035716208_1_Folsomia candida
XP_033117259_1_Anneissia japonica
MCG7891854_1_Candidatus Thiodiazotropha taylori
MBV2113503_1_NA
MBN1968088_1_Candidatus Delongbacteria bacterium
MBP1526238_1_Spiroplasma ixodetis
QTE03999_1_Motacilla cinerea parvoviridae sp.
QTE03832_1_Lanius cristatus densovirus
QWC64890_1_Bombyx mori nucleopolyhedrovirus
AKU37609_1_Bombyx mori bidensovirus
YP_009506248_1_Bombyx mori bidensovirus
YP_007714631_1_Bombyx mori bidensovirus
BAA85362_1_Bombyx mori bidensovirus
DAC81757_1_Dog feces bidnaparvovirus
DAC80328_1_Tasmanian devil feces bidnavirus 1
DAD57248_1_Adintovirus sp.
KAG8456984_1_Diacronema lutheri
PCJ29139_1_Rickettsiales bacterium
MCG8575027_1_Flavobacteriales bacterium
EFX63243_1_Daphnia pulex
NBU97020_1_Spirochaetia bacterium
NDD85271_1_bacterium
CAD7233883_1_Cyprideis torosa
PRP73604_1_Planoprotostelium fungivorum
MBJ7429622_1_Bacteroidia bacterium
XP_003100225_1_Caenorhabditis remanei
XP_005831377_1_Guillardia theta
KXZ46501_1_Gonium pectorale
XP_014524190_1_Vigna radiata
MCE2507535_1_Nitrosopumilaceae archaeon
WP_205732697_1_Acinetobacter baumannii
XP_023227951_1_Centruroides sculpturatus
DAC81320_1_Alphadintovirus mayetiola
MBW0464493_1_Austropuccinia psidii
XP_046399469_1_Ischnura elegans
CAF4383638_1_Rotaria magnacalcarata
KAG0418442_1_Dictyocoela muelleri
ELT91154_1_Capitella teleta
MCG8571961_1_Spirochaetes bacterium
CAG2190273_1_Mytilus edulis
IMGVR_UViG_3300019739_000002_no_taxonomy_3300019739_Ga0194012_10000052
IMGVR_UViG_3300042062_000004_r__Duplodnaviria_k__Heunggongvirae_p__Uroviricota_c__Caudoviricetes_____3300042062_Ga0451379_001063_7083_7631
IMGVR_UViG_3300042064_000122_r__Varidnaviria_k__Bamfordvirae_p__Preplasmiviricota_c__Polintoviricetes_o__Orthopolintovirales_f__Adintoviridae___3300042064_Ga0451392_014174_1601_2695
IMGVR_UViG_3300042080_000077_no_taxonomy_3300042080_Ga0451378_000950_12939_13850
IMGVR_UViG_3300042080_000073_r__Varidnaviria_k__Bamfordvirae_p__Preplasmiviricota_c__Polintoviricetes_o__Orthopolintovirales_f__Adintoviridae___3300042080_Ga0451378_019002_407_1549
IMGVR_UViG_3300003766_000011_no_taxonomy_3300003766_Ga0049104_1006931
IMGVR_UViG_3300026832_000010_no_taxonomy_3300026832_Ga0209561_10006852
IMGVR_UViG_3300003779_000024_no_taxonomy_3300003779_Ga0049103_10010421
IMGVR_UViG_3300027549_000010_no_taxonomy_3300027549_Ga0209458_10003726
IMGVR_UViG_3300003906_000004_r__Varidnaviria_k__Bamfordvirae_______3300003906_JGI26667J51740_100003752
IMGVR_UViG_3300005979_000005_no_taxonomy_3300005979_Ga0056132_10131533
IMGVR_UViG_3300027557_000009_no_taxonomy_3300027557_Ga0209679_10211573
IMGVR_UViG_3300027620_000014_no_taxonomy_3300027620_Ga0209260_100008073
IMGVR_UViG_3300027602_000006_no_taxonomy_3300027602_Ga0209781_10155032
IMGVR_UViG_3300006912_000009_no_taxonomy_3300006912_Ga0056114_10026174
IMGVR_UViG_3300027208_000011_no_taxonomy_3300027208_Ga0209671_10290312
IMGVR_UViG_3300027098_000016_no_taxonomy_3300027098_Ga0209367_10147046
IMGVR_UViG_3300027098_000014_no_taxonomy_3300027098_Ga0209367_10188713
IMGVR_UViG_3300027544_000022_no_taxonomy_3300027544_Ga0209568_10078951
IMGVR_UViG_3300006162_000694_r__Varidnaviria_k__Bamfordvirae_p__Preplasmiviricota_c__Polintoviricetes_o__Orthopolintovirales____3300006162_Ga0075030_1000117552
IMGVR_UViG_3300027911_000652_r__Varidnaviria_k__Bamfordvirae_p__Preplasmiviricota_c__Polintoviricetes_o__Orthopolintovirales____3300027911_Ga0209698_100144192
IMGVR_UViG_3300031902_000911_r__Varidnaviria_k__Bamfordvirae_p__Preplasmiviricota_c__Polintoviricetes_o__Orthopolintovirales____3300031902_Ga0302322_1000518974
IMGVR_UViG_3300009597_000327_r__Varidnaviria_k__Bamfordvirae_p__Preplasmiviricota_c__Polintoviricetes_o__Orthopolintovirales____3300009597_Ga0105259_10002931
IMGVR_UViG_3300014864_000050_r__Varidnaviria_k__Bamfordvirae_p__Preplasmiviricota_c__Polintoviricetes_o__Orthopolintovirales____3300014864_Ga0180068_10001491
IMGVR_UViG_3300014878_000020_r__Varidnaviria_k__Bamfordvirae_p__Preplasmiviricota_c__Polintoviricetes_o__Orthopolintovirales____3300014878_Ga0180065_10001441
IMGVR_UViG_3300020123_000281_r__Varidnaviria_k__Bamfordvirae_p__Preplasmiviricota_c__Polintoviricetes_o__Orthopolintovirales_f__Adintoviridae___3300020123_Ga0210607_129192158
IMGVR_UViG_3300027942_000052_no_taxonomy_3300027942_Ga0209841_10013753
IMGVR_UViG_3300028430_000607_r__Varidnaviria_k__Bamfordvirae_p__Preplasmiviricota_c__Polintoviricetes_o__Orthopolintovirales_f__Adintoviridae___3300028430_Ga0307119_121628904
IMGVR_UViG_3300033060_000037_r__Varidnaviria_k__Bamfordvirae_p__Preplasmiviricota_c__Polintoviricetes_o__Orthopolintovirales____3300033060_Ga0364467_107649102
IMGVR_UViG_3300050159_000113_no_taxonomy_3300050159_Ga0501173_002622_2_3742
IMGVR_UViG_3300033082_000004_r__Varidnaviria_k__Bamfordvirae_p__Preplasmiviricota_c__Polintoviricetes_o__Orthopolintovirales____3300033082_Ga0366740_15491043
IMGVR_UViG_3300033098_000027_r__Varidnaviria_k__Bamfordvirae_p__Preplasmiviricota_c__Polintoviricetes_o__Orthopolintovirales____3300033098_Ga0366743_14556882
IMGVR_UViG_3300033114_000150_r__Varidnaviria_k__Bamfordvirae_p__Preplasmiviricota_c__Polintoviricetes_o__Orthopolintovirales____3300033114_Ga0364476_112736171
IMGVR_UViG_3300027341_000003_r__Varidnaviria_k__Bamfordvirae_______3300027341_Ga0209453_10003951
IMGVR_UViG_3300027372_000007_no_taxonomy_3300027372_Ga0209452_10002221
IMGVR_UViG_3300044980_000274_no_taxonomy_3300044980_Ga0454604_0017081_70_3915
IMGVR_UViG_3300027911_000583_r__Varidnaviria_k__Bamfordvirae_p__Preplasmiviricota_c__Polintoviricetes_o__Orthopolintovirales____3300027911_Ga0209698_100375331
IMGVR_UViG_3300040792_000046_no_taxonomy_3300040792_Ga0450796_087752_6615_10574
IMGVR_UViG_3300040794_000071_r__Varidnaviria_k__Bamfordvirae_______3300040794_Ga0450798_048875_3197_7156
IMGVR_UViG_3300040793_000049_r__Varidnaviria_k__Bamfordvirae_______3300040793_Ga0450797_303915_231_4190
DAC81568_1_Hydra adintovirus
XP_047144678_1_Hydra vulgaris
KAG1651866_1_Nymphon striatum
IMGVR_UViG_3300010030_000017_no_taxonomy_3300010030_Ga0126338_100016391
MAG_viral_001-00000002_1
IMGVR_UViG_3300021297_000161_no_taxonomy_3300021297_Ga0210369_10118012
XP_048587295_1_Nematostella vectensis
IMGVR_UViG_3300013099_002441_no_taxonomy_3300013099_Ga0164315_100304983
MCG8403668_1_Phycisphaerales bacterium
IMGVR_UViG_3300027611_000025_r__Varidnaviria_k__Bamfordvirae_p__Preplasmiviricota_c__Polintoviricetes_o__Orthopolintovirales____3300027611_Ga0210059_100729131
MCG8374751_1_Balneolales bacterium
IMGVR_UViG_3300027611_000027_r__Duplodnaviria_k__Heunggongvirae_p__Uroviricota_c__Caudoviricetes_____3300027611_Ga0210059_10078315
2.5

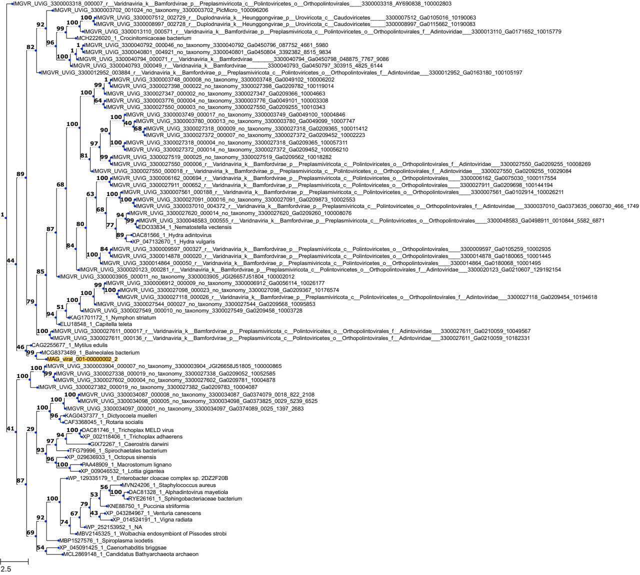
IMGVR_UViG_3300003318_000007_r__Varidnaviria_k__Bamfordvirae_p__Preplasmiviricota_c__Polintoviricetes_o__Orthopolintovirales____3300003318_AY690838_100002803
IMGVR_UViG_3300003702_001024_no_taxonomy_3300003702_PicMicro_100096206
IMGVR_UViG_3300007512_002729_r__Duplodnaviria_k__Heunggongvirae_p__Uroviricota_c__Caudoviricetes_____3300007512_Ga0105016_10190063
IMGVR_UViG_3300008997_002728_r__Duplodnaviria_k__Heunggongvirae_p__Uroviricota_c__Caudoviricetes_____3300008997_Ga0115662_10190083
IMGVR_UViG_3300013110_000571_r__Varidnaviria_k__Bamfordvirae_p__Preplasmiviricota_c__Polintoviricetes_o__Orthopolintovirales_f__Adintoviridae___3300013110_Ga0171652_10015779
MCH2226020_1_Crocinitomicaceae bacterium
IMGVR_UViG_3300040792_000046_no_taxonomy_3300040792_Ga0450796_087752_4661_5980
IMGVR_UViG_3300040801_004921_no_taxonomy_3300040801_Ga0450804_3392382_8515_9834
IMGVR_UViG_3300040794_000071_r__Varidnaviria_k__Bamfordvirae_______3300040794_Ga0450798_048875_7767_9086
IMGVR_UViG_3300040793_000049_r__Varidnaviria_k__Bamfordvirae_______3300040793_Ga0450797_303915_4825_6144
IMGVR_UViG_3300012952_003884_r__Varidnaviria_k__Bamfordvirae_p__Preplasmiviricota_c__Polintoviricetes_o__Orthopolintovirales_f__Adintoviridae___3300012952_Ga0163180_100105197
IMGVR_UViG_3300003748_000008_no_taxonomy_3300003748_Ga0049102_100006202
IMGVR_UViG_3300027398_000022_no_taxonomy_3300027398_Ga0209782_100119014
IMGVR_UViG_3300027347_000002_no_taxonomy_3300027347_Ga0209366_10004663
IMGVR_UViG_3300003776_000004_no_taxonomy_3300003776_Ga0049101_100003308
IMGVR_UViG_3300027550_000003_no_taxonomy_3300027550_Ga0209255_10010343
IMGVR_UViG_3300003749_000017_no_taxonomy_3300003749_Ga0049100_10004846
IMGVR_UViG_3300003780_000013_no_taxonomy_3300003780_Ga0049099_10007747
IMGVR_UViG_3300027318_000009_no_taxonomy_3300027318_Ga0209365_100011412
IMGVR_UViG_3300027372_000007_no_taxonomy_3300027372_Ga0209452_10002223
IMGVR_UViG_3300027318_000004_no_taxonomy_3300027318_Ga0209365_100057311
IMGVR_UViG_3300027372_000014_no_taxonomy_3300027372_Ga0209452_100056210
IMGVR_UViG_3300027519_000025_no_taxonomy_3300027519_Ga0209562_10018282
IMGVR_UViG_3300027550_000006_r__Varidnaviria_k__Bamfordvirae_p__Preplasmiviricota_c__Polintoviricetes_o__Orthopolintovirales_f__Adintoviridae___3300027550_Ga0209255_10008269
IMGVR_UViG_3300027550_000018_r__Varidnaviria_k__Bamfordvirae_p__Preplasmiviricota_c__Polintoviricetes_o__Orthopolintovirales_f__Adintoviridae___3300027550_Ga0209255_10029084
IMGVR_UViG_3300006162_000694_r__Varidnaviria_k__Bamfordvirae_p__Preplasmiviricota_c__Polintoviricetes_o__Orthopolintovirales____3300006162_Ga0075030_1000117554
IMGVR_UViG_3300027911_000652_r__Varidnaviria_k__Bamfordvirae_p__Preplasmiviricota_c__Polintoviricetes_o__Orthopolintovirales____3300027911_Ga0209698_100144194
IMGVR_UViG_3300007561_000188_r__Varidnaviria_k__Bamfordvirae_p__Preplasmiviricota_c__Polintoviricetes_o__Orthopolintovirales____3300007561_Ga0102914_100026211
IMGVR_UViG_3300027091_000016_no_taxonomy_3300027091_Ga0209873_10002553
IMGVR_UViG_3300037010_004372_r__Varidnaviria_k__Bamfordvirae_p__Preplasmiviricota_c__Polintoviricetes_o__Orthopolintovirales_f__Adintoviridae___3300037010_Ga0373635_0060730_466_1749
IMGVR_UViG_3300027620_000014_no_taxonomy_3300027620_Ga0209260_100008076
IMGVR_UViG_3300048583_000555_r__Varidnaviria_k__Bamfordvirae_p__Preplasmiviricota_c__Polintoviricetes_o__Orthopolintovirales____3300048583_Ga0498911_0010844_5582_6871
EDO33834_1_Nematostella vectensis
DAC81566_1_Hydra adintovirus
XP_047132670_1_Hydra vulgaris
IMGVR_UViG_3300009597_000327_r__Varidnaviria_k__Bamfordvirae_p__Preplasmiviricota_c__Polintoviricetes_o__Orthopolintovirales____3300009597_Ga0105259_10002935
IMGVR_UViG_3300014878_000020_r__Varidnaviria_k__Bamfordvirae_p__Preplasmiviricota_c__Polintoviricetes_o__Orthopolintovirales____3300014878_Ga0180065_10001445
IMGVR_UViG_3300014864_000050_r__Varidnaviria_k__Bamfordvirae_p__Preplasmiviricota_c__Polintoviricetes_o__Orthopolintovirales____3300014864_Ga0180068_10001495
IMGVR_UViG_3300020123_000281_r__Varidnaviria_k__Bamfordvirae_p__Preplasmiviricota_c__Polintoviricetes_o__Orthopolintovirales_f__Adintoviridae___3300020123_Ga0210607_129192154
IMGVR_UViG_3300003905_000011_no_taxonomy_3300003905_JGI26657J51804_100002012
IMGVR_UViG_3300006912_000009_no_taxonomy_3300006912_Ga0056114_10026177
IMGVR_UViG_3300027098_000023_no_taxonomy_3300027098_Ga0209367_10176574
IMGVR_UViG_3300027118_000026_r__Varidnaviria_k__Bamfordvirae_p__Preplasmiviricota_c__Polintoviricetes_o__Orthopolintovirales_f__Adintoviridae___3300027118_Ga0209454_10194618
IMGVR_UViG_3300027544_000027_no_taxonomy_3300027544_Ga0209568_10095853
IMGVR_UViG_3300027549_000010_no_taxonomy_3300027549_Ga0209458_10003728
KAG1701172_1_Nymphon striatum
ELU18548_1_Capitella teleta
IMGVR_UViG_3300027611_000017_r__Varidnaviria_k__Bamfordvirae_p__Preplasmiviricota_c__Polintoviricetes_o__Orthopolintovirales_f__Adintoviridae___3300027611_Ga0210059_10049567
IMGVR_UViG_3300027611_000136_r__Varidnaviria_k__Bamfordvirae_p__Preplasmiviricota_c__Polintoviricetes_o__Orthopolintovirales_f__Adintoviridae___3300027611_Ga0210059_10182331
CAG2255677_1_Mytilus edulis
MCG8373489_1_Balneolales bacterium
MAG_viral_001-0000002_2
IMGVR_UViG_3300003904_000007_no_taxonomy_3300003904_JGI26658J51805_100000865
IMGVR_UViG_3300027338_000019_no_taxonomy_3300027338_Ga0209052_10052585
IMGVR_UViG_3300027602_000004_no_taxonomy_3300027602_Ga0209781_10004878
IMGVR_UViG_3300027382_000019_no_taxonomy_3300027382_Ga0209783_10004087
IMGVR_UViG_3300034087_000008_no_taxonomy_3300034087_Ga0374079_0018_822_2108
IMGVR_UViG_3300034098_000005_no_taxonomy_3300034098_Ga0373825_0029_5239_6525
IMGVR_UViG_3300034097_000001_no_taxonomy_3300034097_Ga0374089_0025_1397_2683
KAG0437377_1_Dictyocoela muelleri
CAF3368045_1_Rotaria socialis
DAC81746_1_Trichoplax MELD virus
XP_002118406_1_Trichoplax adhaerens
GIX72267_1_Caerostris darwini
TFG79996_1_Spirochaetales bacterium
XP_029636933_1_Octopus sinensis
PAA48909_1_Macrostomum lignano
XP_009046532_1_Lottia gigantea
WP_129335179_1_Enterobacter cloacae complex sp. 2DZ2F20B
MVN24206_1_Staphylococcus aureus
DAC81328_1_Alphadintovirus mayetiola
RYE26161_1_Sphingobacteriaceae bacterium
KNE88750_1_Puccinia striiformis
XP_043284967_1_Venturia canescens
XP_014524191_1_Vigna radiata
WP_252153952_1_NA
MBV2145325_1_Wolbachia endosymbiont of Pissodes strobi
MBP1527576_1_Spiroplasma ixodetis
XP_045091425_1_Caenorhabditis briggsae
MCL2869148_1_Candidatus Bathyarchaeota archaeon
2.5

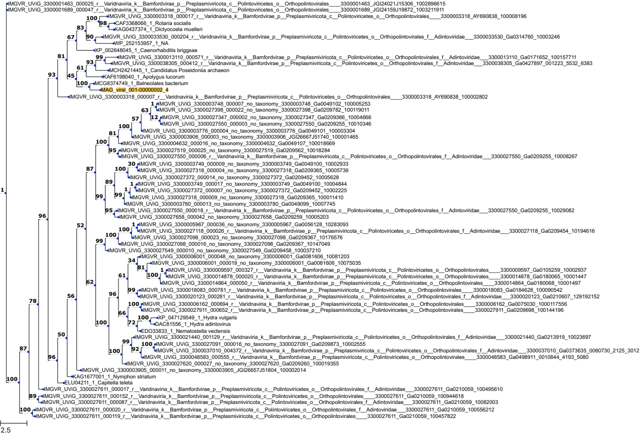
IMGVR_UViG_3300001463_000025_r__Varidnaviria_k__Bamfordvirae_p__Preplasmiviricota_c__Polintoviricetes_o__Orthopolintovirales____3300001463_JGI24021J15306_1002896615
IMGVR_UViG_3300001689_000047_r__Varidnaviria_k__Bamfordvirae_p__Preplasmiviricota_c__Polintoviricetes_o__Orthopolintovirales____3300001689_JGI24159J19872_1003211911
IMGVR_UViG_3300003318_000017_r__Varidnaviria_k__Bamfordvirae_p__Preplasmiviricota_c__Polintoviricetes_o__Orthopolintovirales____3300003318_AY690838_100008196
CAF3368066_1_Rotaria socialis
KAG0437374_1_Dictyocoela muelleri
IMGVR_UViG_3300033530_000204_r__Varidnaviria_k__Bamfordvirae_p__Preplasmiviricota_c__Polintoviricetes_o__Orthopolintovirales_f__Adintoviridae___3300033530_Ga0314760_10003246
WP_252153957_1_NA
XP_002648045_1_Caenorhabditis briggsae
IMGVR_UViG_3300013110_000571_r__Varidnaviria_k__Bamfordvirae_p__Preplasmiviricota_c__Polintoviricetes_o__Orthopolintovirales_f__Adintoviridae___3300013110_Ga0171652_100157711
IMGVR_UViG_3300038305_000412_r__Varidnaviria_k__Bamfordvirae_p__Preplasmiviricota_c__Polintoviricetes_o__Orthopolintovirales_f__Adintoviridae___3300038305_Ga0427897_001223_5532_6383
MCH2421445_1_Candidatus Poseidoniia archaeon
KAF6198040_1_Apolygus lucorum
MCG8374749_1_Balneolales bacterium
MAG_viral_001-00000002_4
IMGVR_UViG_3300003318_000007_r__Varidnaviria_k__Bamfordvirae_p__Preplasmiviricota_c__Polintoviricetes_o__Orthopolintovirales____3300003318_AY690838_100002802
IMGVR_UViG_3300003748_000007_no_taxonomy_3300003748_Ga0049102_100005253
IMGVR_UViG_3300027398_000022_no_taxonomy_3300027398_Ga0209782_100119011
IMGVR_UViG_3300027347_000002_no_taxonomy_3300027347_Ga0209366_10004666
IMGVR_UViG_3300027550_000003_no_taxonomy_3300027550_Ga0209255_10010346
IMGVR_UViG_3300003776_000004_no_taxonomy_3300003776_Ga0049101_100003304
IMGVR_UViG_3300003906_000003_no_taxonomy_3300003906_JGI26667J51740_100001465
IMGVR_UViG_3300004632_000016_no_taxonomy_3300004632_Ga0049107_100018669
IMGVR_UViG_3300027519_000025_no_taxonomy_3300027519_Ga0209562_10018284
IMGVR_UViG_3300027550_000006_r__Varidnaviria_k__Bamfordvirae_p__Preplasmiviricota_c__Polintoviricetes_o__Orthopolintovirales_f__Adintoviridae___3300027550_Ga0209255_10008267
IMGVR_UViG_3300003749_000009_no_taxonomy_3300003749_Ga0049100_10002933
IMGVR_UViG_3300027318_000004_no_taxonomy_3300027318_Ga0209365_10005739
IMGVR_UViG_3300027372_000014_no_taxonomy_3300027372_Ga0209452_10005628
IMGVR_UViG_3300003749_000017_no_taxonomy_3300003749_Ga0049100_10004844
IMGVR_UViG_3300027372_000007_no_taxonomy_3300027372_Ga0209452_10002225
IMGVR_UViG_3300027318_000009_no_taxonomy_3300027318_Ga0209365_100011410
IMGVR_UViG_3300003780_000013_no_taxonomy_3300003780_Ga0049099_10007745
IMGVR_UViG_3300027550_000018_r__Varidnaviria_k__Bamfordvirae_p__Preplasmiviricota_c__Polintoviricetes_o__Orthopolintovirales_f__Adintoviridae___3300027550_Ga0209255_10029082
IMGVR_UViG_3300027658_000042_no_taxonomy_3300027658_Ga0209259_10005203
IMGVR_UViG_3300005967_000036_no_taxonomy_3300005967_Ga0056128_10283093
IMGVR_UViG_3300027118_000026_r__Varidnaviria_k__Bamfordvirae_p__Preplasmiviricota_c__Polintoviricetes_o__Orthopolintovirales_f__Adintoviridae___3300027118_Ga0209454_10194616
IMGVR_UViG_3300027098_000023_no_taxonomy_3300027098_Ga0209367_10176576
IMGVR_UViG_3300027098_000016_no_taxonomy_3300027098_Ga0209367_10147049
IMGVR_UViG_3300027549_000010_no_taxonomy_3300027549_Ga0209458_100037210
IMGVR_UViG_3300006001_000048_no_taxonomy_3300006001_Ga0081606_10081203
IMGVR_UViG_3300006001_000019_no_taxonomy_3300006001_Ga0081606_10075035
IMGVR_UViG_3300009597_000327_r__Varidnaviria_k__Bamfordvirae_p__Preplasmiviricota_c__Polintoviricetes_o__Orthopolintovirales____3300009597_Ga0105259_10002937
IMGVR_UViG_3300014878_000020_r__Varidnaviria_k__Bamfordvirae_p__Preplasmiviricota_c__Polintoviricetes_o__Orthopolintovirales____3300014878_Ga0180065_10001447
IMGVR_UViG_3300014864_000050_r__Varidnaviria_k__Bamfordvirae_p__Preplasmiviricota_c__Polintoviricetes_o__Orthopolintovirales____3300014864_Ga0180068_10001497
IMGVR_UViG_3300018083_000781_r__Varidnaviria_k__Bamfordvirae_p__Preplasmiviricota_c__Polintoviricetes_o__Orthopolintovirales____3300018083_Ga0184628_100090542
IMGVR_UViG_3300020123_000281_r__Varidnaviria_k__Bamfordvirae_p__Preplasmiviricota_c__Polintoviricetes_o__Orthopolintovirales_f__Adintoviridae___3300020123_Ga0210607_129192152
IMGVR_UViG_3300006162_000694_r__Varidnaviria_k__Bamfordvirae_p__Preplasmiviricota_c__Polintoviricetes_o__Orthopolintovirales____3300006162_Ga0075030_1000117556
IMGVR_UViG_3300027911_000652_r__Varidnaviria_k__Bamfordvirae_p__Preplasmiviricota_c__Polintoviricetes_o__Orthopolintovirales____3300027911_Ga0209698_100144196
XP_047129549_1_Hydra vulgaris
DAC81556_1_Hydra adintovirus
EDO33833_1_Nematostella vectensis
IMGVR_UViG_3300021440_001129_r__Varidnaviria_k__Bamfordvirae_p__Preplasmiviricota_c__Polintoviricetes_o__Orthopolintovirales_f__Adintoviridae___3300021440_Ga0213919_10023697
IMGVR_UViG_3300027091_000016_no_taxonomy_3300027091_Ga0209873_10002555
IMGVR_UViG_3300037010_004372_r__Varidnaviria_k__Bamfordvirae_p__Preplasmiviricota_c__Polintoviricetes_o__Orthopolintovirales_f__Adintoviridae___3300037010_Ga0373635_0060730_2125_3012
IMGVR_UViG_3300048583_000555_r__Varidnaviria_k__Bamfordvirae_p__Preplasmiviricota_c__Polintoviricetes_o__Orthopolintovirales____3300048583_Ga0498911_0010844_4193_5080
IMGVR_UViG_3300027620_000027_no_taxonomy_3300027620_Ga0209260_100019355
IMGVR_UViG_3300003905_000011_no_taxonomy_3300003905_JGI26657J51804_100002014
KAG1677001_1_Nymphon striatum
ELU04211_1_Capitella teleta
IMGVR_UViG_3300027611_000017_r__Varidnaviria_k__Bamfordvirae_p__Preplasmiviricota_c__Polintoviricetes_o__Orthopolintovirales_f__Adintoviridae___3300027611_Ga0210059_100495610
IMGVR_UViG_3300027611_000152_r__Varidnaviria_k__Bamfordvirae_p__Preplasmiviricota_c__Polintoviricetes_o__Orthopolintovirales____3300027611_Ga0210059_100944618
IMGVR_UViG_3300027611_000087_r__Varidnaviria_k__Bamfordvirae_p__Preplasmiviricota_c__Polintoviricetes_o__Orthopolintovirales_f__Adintoviridae___3300027611_Ga0210059_10082003
IMGVR_UViG_3300027611_000020_r__Varidnaviria_k__Bamfordvirae_p__Preplasmiviricota_c__Polintoviricetes_o__Orthopolintovirales_f__Adintoviridae___3300027611_Ga0210059_100556212
IMGVR_UViG_3300027611_000119_r__Varidnaviria_k__Bamfordvirae_p__Preplasmiviricota_c__Polintoviricetes_o__Orthopolintovirales____3300027611_Ga0210059_100457822
2.5

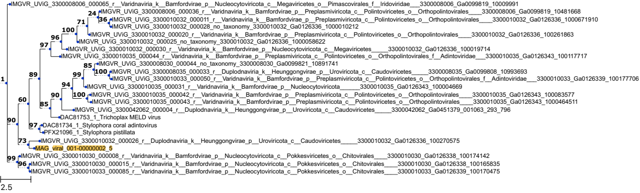
IMGVR_UViG_3300008006_000065_r__Varidnaviria_k__Bamfordvirae_p__Nucleocytoviricota_c__Megaviricetes_o__Pimascovirales_f__Iridoviridae____3300008006_Ga0099819_10009991
IMGVR_UViG_3300008006_000036_r__Varidnaviria_k__Bamfordvirae_p__Preplasmiviricota_c__Polintoviricetes_o__Orthopolintovirales____3300008006_Ga0099819_10481668
IMGVR_UViG_3300010032_000011_r__Varidnaviria_k__Bamfordvirae_p__Preplasmiviricota_c__Polintoviricetes_o__Orthopolintovirales____3300010032_Ga0126336_1000671910
IMGVR_UViG_3300010032_000028_no_taxonomy_3300010032_Ga0126336_1000010212
IMGVR_UViG_3300010032_000020_r__Varidnaviria_k__Bamfordvirae_p__Preplasmiviricota_c__Polintoviricetes_o__Orthopolintovirales____3300010032_Ga0126336_100261863
IMGVR_UViG_3300010032_000025_no_taxonomy_3300010032_Ga0126336_1000058622
IMGVR_UViG_3300010032_000030_r__Varidnaviria_k__Bamfordvirae_p__Nucleocytoviricota_c__Megaviricetes____3300010032_Ga0126336_100019714
IMGVR_UViG_3300010035_000044_r__Varidnaviria_k__Bamfordvirae_p__Preplasmiviricota_c__Polintoviricetes_o__Orthopolintovirales_f__Adintoviridae____3300010035_Ga0126343_100117717
IMGVR_UViG_3300008030_000044_no_taxonomy_3300008030_Ga0099821_10891741
IMGVR_UViG_3300008035_000033_r__Duplodnaviria_k__Heunggongvirae_p__Uroviricota_c__Caudoviricetes____3300008035_Ga0099808_10993693
IMGVR_UViG_3300010033_000050_r__Varidnaviria_k__Bamfordvirae_p__Preplasmiviricota_c__Polintoviricetes_o__Orthopolintovirales_f__Adintoviridae____3300010033_Ga0126339_100177706
IMGVR_UViG_3300010035_000031_r__Varidnaviria_k__Bamfordvirae_p__Nucleocytoviricota______3300010035_Ga0126343_100004669
IMGVR_UViG_3300010035_000042_r__Varidnaviria_k__Bamfordvirae_p__Preplasmiviricota_c__Polintoviricetes_o__Orthopolintovirales____3300010035_Ga0126343_100083577
IMGVR_UViG_3300010035_000043_r__Varidnaviria_k__Bamfordvirae_p__Preplasmiviricota_c__Polintoviricetes_o__Orthopolintovirales____3300010035_Ga0126343_1000464511
IMGVR_UViG_3300042062_000004_r__Duplodnaviria_k__Heunggongvirae_p__Uroviricota_c__Caudoviricetes____3300042062_Ga0451379_001063_293_796
DAC81753_1_Trichoplax MELD virus
DAC81734_1_Stylophora coral adintovirus
PFX21096_1_Stylophora pistillata
IMGVR_UViG_3300010032_000026_r__Duplodnaviria_k__Heunggongvirae_p__Uroviricota_c__Caudoviricetes____3300010032_Ga0126336_100270575
MAG_viral_001-00000002_5
IMGVR_UViG_3300010030_000008_r__Varidnaviria_k__Bamfordvirae_p__Nucleocytoviricota_c__Pokkesviricetes_o__Chitovirales____3300010030_Ga0126338_100174142
IMGVR_UViG_3300010030_000015_r__Varidnaviria_k__Bamfordvirae_p__Nucleocytoviricota_c__Pokkesviricetes_o__Chitovirales____3300010030_Ga0126338_100165835
IMGVR_UViG_3300010033_000085_r__Varidnaviria_k__Bamfordvirae_p__Nucleocytoviricota_c__Pokkesviricetes_o__Chitovirales____3300010033_Ga0126339_100170475
24
36
71
100
96
97
85
100
99
94
100
90
89
85
60
97
1
90
73
99
96
2.5

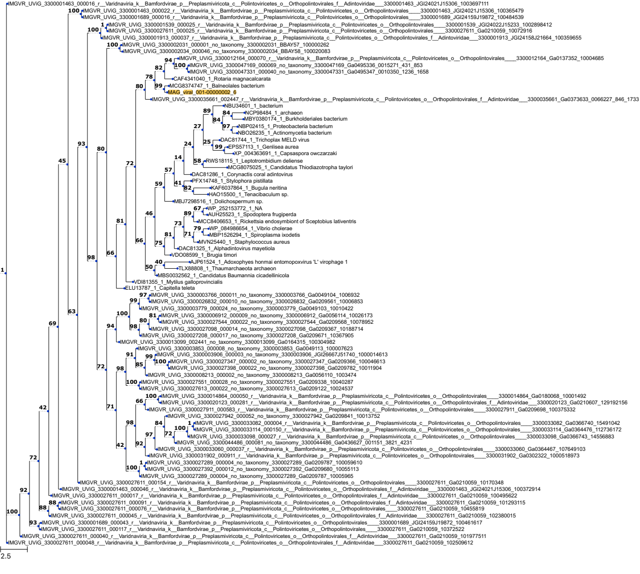

IMGVR_UViG_3300001463_000016_r__Varidnaviria_k__Bamfordvirae_p__Preplasmiviricota_c__Polintoviricetes_o__Orthopolintovirales____3300001463_JGI24021J15306_1003697111
IMGVR_UViG_3300001463_000022_r__Varidnaviria_k__Bamfordvirae_p__Preplasmiviricota_c__Polintoviricetes_o__Orthopolintovirales____3300001463_JGI24021J15306_100365479
IMGVR_UViG_3300001689_000016_r__Varidnaviria_k__Bamfordvirae_p__Preplasmiviricota_c__Polintoviricetes_o__Orthopolintovirales____3300001689_JGI24159J19872_100484539
IMGVR_UViG_3300001539_000025_r__Varidnaviria_k__Bamfordvirae_p__Preplasmiviricota_c__Polintoviricetes_o__Orthopolintovirales____3300001539_JGI24022J15233_1002898412
IMGVR_UViG_3300027611_000025_r__Varidnaviria_k__Bamfordvirae_p__Preplasmiviricota_c__Polintoviricetes_o__Orthopolintovirales____3300027611_Ga0210059_10072916
IMGVR_UViG_3300001913_000037_r__Varidnaviria_k__Bamfordvirae_p__Preplasmiviricota_c__Polintoviricetes_o__Orthopolintovirales_f__Adintoviridae____3300001913_JGI24158J21664_100359655
IMGVR_UViG_3300002031_000001_no_taxonomy_3300002031_BBAY57_100000262
IMGVR_UViG_3300002034_000046_no_taxonomy_3300002034_BBAY58_100020083
IMGVR_UViG_3300012164_000070_r__Varidnaviria_k__Bamfordvirae_p__Preplasmiviricota_c__Polintoviricetes_o__Orthopolintovirales____3300012164_Ga0137352_10004685
IMGVR_UViG_3300047169_000069_no_taxonomy_3300047169_Ga0495336_0015271_431_853
IMGVR_UViG_3300047331_000040_no_taxonomy_3300047331_Ga0495347_0010350_1236_1658
CAF4341040_1_Rotaria magnacalcarata
MCG8374747_1_Balneolales bacterium
MAG_viral_001-00000002_6
IMGVR_UViG_3300035661_002447_r__Varidnaviria_k__Bamfordvirae_p__Preplasmiviricota_c__Polintoviricetes_o__Orthopolintovirales_f__Adintoviridae____3300035661_Ga0373633_0066227_846_1733
NBU34601_1_bacterium
NCP98484_1_archaeon
MBY0380174_1_Burkholderiales bacterium
NBP02415_1_Proteobacteria bacterium
NBO26235_1_Actinomycetia bacterium
DAC81744_1_Trichoplax MELD virus
EPS57113_1_Genlisea aurea
XP_004363691_1_Capsaspora owczarzaki
RWS18115_1_Leptotrombidium deliense
MCG8075025_1_Candidatus Thiodiazotropha taylori
DAC81286_1_Corynactis coral adintovirus
PFX14748_1_Stylophora pistillata
KAF6037864_1_Bugula neritina
HAO15500_1_Tenacibaculum sp.
MBJ7298516_1_Dolichospermum sp.
WP_252153772_1_NA
AUH25523_1_Spodoptera frugiperda
MCC8406653_1_Rickettsia endosymbiont of Sceptobius lativentris
WP_084986654_1_Vibrio cholerae
MBP1526294_1_Spiroplasma ixodetis
MVN25440_1_Staphylococcus aureus
DAC81325_1_Alphadintovirus mayetiola
VDO08599_1_Brugia timori
AJP61524_1_Adoxophyes honmai entomopoxvirus 'L' virophage 1
TLX88808_1_Thaumarchaeota archaeon
MBS0032562_1_Candidatus Baumannia cicadellinicola
VDI81355_1_Mytilus galloprovincialis
ELU13787_1_Capitella teleta
IMGVR_UViG_3300003766_000011_no_taxonomy_3300003766_Ga0049104_1006932
IMGVR_UViG_3300026832_000010_no_taxonomy_3300026832_Ga0209561_100006853
IMGVR_UViG_3300003779_000024_no_taxonomy_3300003779_Ga0049103_10010422
IMGVR_UViG_3300006912_000009_no_taxonomy_3300006912_Ga0056114_10026173
IMGVR_UViG_3300027544_000022_no_taxonomy_3300027544_Ga0209568_10078952
IMGVR_UViG_3300027098_000014_no_taxonomy_3300027098_Ga0209367_10188714
IMGVR_UViG_3300027208_000017_no_taxonomy_3300027208_Ga0209671_10367905
IMGVR_UViG_3300013099_002441_no_taxonomy_3300013099_Ga0164315_100304982
IMGVR_UViG_3300003853_000008_no_taxonomy_3300003853_Ga0049113_100007623
IMGVR_UViG_3300003906_000003_no_taxonomy_3300003906_JGI26667J51740_1000014613
IMGVR_UViG_3300027347_000002_no_taxonomy_3300027347_Ga0209366_100046613
IMGVR_UViG_3300027398_000022_no_taxonomy_3300027398_Ga0209782_10011904
IMGVR_UViG_3300008213_000002_no_taxonomy_3300008213_Ga0056110_1003474
IMGVR_UViG_3300027551_000028_no_taxonomy_3300027551_Ga0209338_10040287
IMGVR_UViG_3300027613_000022_no_taxonomy_3300027613_Ga0209122_10024537
IMGVR_UViG_3300014864_000050_r__Varidnaviria_k__Bamfordvirae_p__Preplasmiviricota_c__Polintoviricetes_o__Orthopolintovirales____3300014864_Ga0180068_10001492
IMGVR_UViG_3300020123_000281_r__Varidnaviria_k__Bamfordvirae_p__Preplasmiviricota_c__Polintoviricetes_o__Orthopolintovirales_f__Adintoviridae____3300020123_Ga0210607_129192156
IMGVR_UViG_3300027911_000583_r__Varidnaviria_k__Bamfordvirae_p__Preplasmiviricota_c__Polintoviricetes_o__Orthopolintovirales____3300027911_Ga0209698_10375332
IMGVR_UViG_3300027942_000052_no_taxonomy_3300027942_Ga0209841_10013752
IMGVR_UViG_3300033082_000004_r__Varidnaviria_k__Bamfordvirae_p__Preplasmiviricota_c__Polintoviricetes_o__Orthopolintovirales____3300033082_Ga0366740_15491042
IMGVR_UViG_3300033114_000150_r__Varidnaviria_k__Bamfordvirae_p__Preplasmiviricota_c__Polintoviricetes_o__Orthopolintovirales____3300033114_Ga0364476_112736172
IMGVR_UViG_3300033098_000027_r__Varidnaviria_k__Bamfordvirae_p__Preplasmiviricota_c__Polintoviricetes_o__Orthopolintovirales____3300033098_Ga0366743_14556883
IMGVR_UViG_3300044486_000081_no_taxonomy_3300044486_Ga0436627_001151_3821_4231
IMGVR_UViG_3300033060_000037_r__Varidnaviria_k__Bamfordvirae_p__Preplasmiviricota_c__Polintoviricetes_o__Orthopolintovirales____3300033060_Ga0364467_107649103
IMGVR_UViG_3300031902_000911_r__Varidnaviria_k__Bamfordvirae_p__Preplasmiviricota_c__Polintoviricetes_o__Orthopolintovirales____3300031902_Ga0302322_1000518973
IMGVR_UViG_3300027289_000004_no_taxonomy_3300027289_Ga0209787_100059610
IMGVR_UViG_3300027392_000012_no_taxonomy_3300027392_Ga0209680_10055113
IMGVR_UViG_3300027289_000004_no_taxonomy_3300027289_Ga0209787_10005965
IMGVR_UViG_3300027611_000154_r__Varidnaviria_k__Bamfordvirae_p__Preplasmiviricota_c__Polintoviricetes_o__Orthopolintovirales____3300027611_Ga0210059_10170348
IMGVR_UViG_3300001463_000046_r__Varidnaviria_k__Bamfordvirae_p__Preplasmiviricota_c__Polintoviricetes_o__Orthopolintovirales_f__Adintoviridae____3300001463_JGI24021J15306_100372914
IMGVR_UViG_3300027611_000017_r__Varidnaviria_k__Bamfordvirae_p__Preplasmiviricota_c__Polintoviricetes_o__Orthopolintovirales_f__Adintoviridae____3300027611_Ga0210059_100495622
IMGVR_UViG_3300027611_000091_r__Varidnaviria_k__Bamfordvirae_p__Preplasmiviricota_c__Polintoviricetes_o__Orthopolintovirales_f__Adintoviridae____3300027611_Ga0210059_101293115
IMGVR_UViG_3300027611_000076_r__Varidnaviria_k__Bamfordvirae_p__Preplasmiviricota_c__Polintoviricetes_o__Orthopolintovirales____3300027611_Ga0210059_10455819
IMGVR_UViG_3300027611_000045_r__Varidnaviria_k__Bamfordvirae_p__Preplasmiviricota_c__Polintoviricetes_o__Orthopolintovirales_f__Adintoviridae____3300027611_Ga0210059_102380015
IMGVR_UViG_3300001689_000043_r__Varidnaviria_k__Bamfordvirae_p__Preplasmiviricota_c__Polintoviricetes_o__Orthopolintovirales_f__Adintoviridae____3300001689_JGI24159J19872_100461617
IMGVR_UViG_3300027611_000117_r__Varidnaviria_k__Bamfordvirae_p__Preplasmiviricota_c__Polintoviricetes_o__Orthopolintovirales____3300027611_Ga0210059_10372522
IMGVR_UViG_3300027611_000040_r__Varidnaviria_k__Bamfordvirae_p__Preplasmiviricota_c__Polintoviricetes_o__Orthopolintovirales_f__Adintoviridae____3300027611_Ga0210059_101977511
IMGVR_UViG_3300027611_000048_r__Varidnaviria_k__Bamfordvirae_p__Preplasmiviricota_c__Polintoviricetes_o__Orthopolintovirales_f__Adintoviridae____3300027611_Ga0210059_102509612
2.5

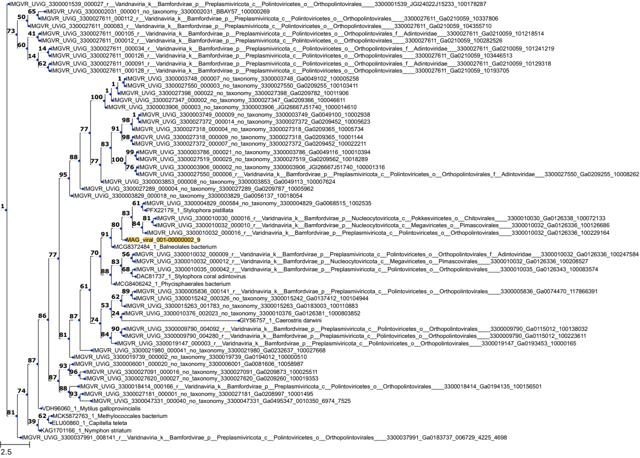
IMGVR_UViG_3300001539_000027_r__Varidnaviria_k__Bamfordvirae_p__Preplasmiviricota_c__Polintoviricetes_o__Orthopolintovirales____3300001539_JGI24022J15233_100178287
IMGVR_UViG_3300002031_000001_no_taxonomy_3300002031_BBAY57_100000269
IMGVR_UViG_3300027611_000112_r__Varidnaviria_k__Bamfordvirae_p__Preplasmiviricota_c__Polintoviricetes_o__Orthopolintovirales____3300027611_Ga0210059_10337806
IMGVR_UViG_3300027611_000083_r__Varidnaviria_k__Bamfordvirae_p__Preplasmiviricota_c__Polintoviricetes_o__Orthopolintovirales____3300027611_Ga0210059_104355710
IMGVR_UViG_3300027611_000105_r__Varidnaviria_k__Bamfordvirae_p__Preplasmiviricota_c__Polintoviricetes_o__Orthopolintovirales_f__Adintoviridae___3300027611_Ga0210059_101218514
IMGVR_UViG_3300027611_000012_r__Varidnaviria_k__Bamfordvirae_p__Preplasmiviricota_c__Polintoviricetes_o__Orthopolintovirales____3300027611_Ga0210059_100282526
IMGVR_UViG_3300027611_000034_r__Varidnaviria_k__Bamfordvirae_p__Preplasmiviricota_c__Polintoviricetes_o__Orthopolintovirales_f__Adintoviridae___3300027611_Ga0210059_101241219
IMGVR_UViG_3300027611_000126_r__Varidnaviria_k__Bamfordvirae_p__Preplasmiviricota_c__Polintoviricetes_o__Orthopolintovirales____3300027611_Ga0210059_103446513
IMGVR_UViG_3300027611_000091_r__Varidnaviria_k__Bamfordvirae_p__Preplasmiviricota_c__Polintoviricetes_o__Orthopolintovirales_f__Adintoviridae___3300027611_Ga0210059_10129318
IMGVR_UViG_3300027611_000128_r__Varidnaviria_k__Bamfordvirae_p__Preplasmiviricota_c__Polintoviricetes_o__Orthopolintovirales____3300027611_Ga0210059_10193705
IMGVR_UViG_3300003748_000007_no_taxonomy_3300003748_Ga0049102_100005258
IMGVR_UViG_3300027550_000003_no_taxonomy_3300027550_Ga0209255_100103411
IMGVR_UViG_3300027398_000022_no_taxonomy_3300027398_Ga0209782_10011906
IMGVR_UViG_3300027347_000002_no_taxonomy_3300027347_Ga0209366_100046611
IMGVR_UViG_3300003906_000003_no_taxonomy_3300003906_JGI26667J51740_1000014610
IMGVR_UViG_3300003749_000009_no_taxonomy_3300003749_Ga0049100_10002938
IMGVR_UViG_3300027372_000014_no_taxonomy_3300027372_Ga0209452_10005623
IMGVR_UViG_3300027318_000004_no_taxonomy_3300027318_Ga0209365_10005734
IMGVR_UViG_3300027318_000009_no_taxonomy_3300027318_Ga0209365_10001144
IMGVR_UViG_3300027372_000007_no_taxonomy_3300027372_Ga0209452_100022211
IMGVR_UViG_3300003786_000021_no_taxonomy_3300003786_Ga0049116_100010394
IMGVR_UViG_3300027519_000025_no_taxonomy_3300027519_Ga0209562_10018289
IMGVR_UViG_3300003906_000002_no_taxonomy_3300003906_JGI26667J51740_100001316
IMGVR_UViG_3300027550_000006_r__Varidnaviria_k__Bamfordvirae_p__Preplasmiviricota_c__Polintoviricetes_o__Orthopolintovirales_f__Adintoviridae___3300027550_Ga0209255_10008262
IMGVR_UViG_3300003853_000008_no_taxonomy_3300003853_Ga0049113_100007624
IMGVR_UViG_3300027289_000004_no_taxonomy_3300027289_Ga0209787_10005962
IMGVR_UViG_3300003829_000018_no_taxonomy_3300003829_Ga0056137_10018054
IMGVR_UViG_3300004829_000584_no_taxonomy_3300004829_Ga0068515_1002535
PFX22179_1_Stylophora pistillata
IMGVR_UViG_3300010030_000016_r__Varidnaviria_k__Bamfordvirae_p__Nucleocytoviricota_c__Pokkesviricetes_o__Chitovirales____3300010030_Ga0126338_100072133
IMGVR_UViG_3300010032_000010_r__Varidnaviria_k__Bamfordvirae_p__Nucleocytoviricota_c__Megaviricetes_o__Pimascovirales____3300010032_Ga0126336_100126686
IMGVR_UViG_3300010032_000016_r__Varidnaviria_k__Bamfordvirae_p__Preplasmiviricota_c__Polintoviricetes_o__Orthopolintovirales____3300010032_Ga0126336_100229164
MAG_viral_001-00000002_9
MCG8372484_1_Balneolales bacterium
IMGVR_UViG_3300010032_000009_r__Varidnaviria_k__Bamfordvirae_p__Preplasmiviricota_c__Polintoviricetes_o__Orthopolintovirales_f__Adintoviridae___3300010032_Ga0126336_100247584
IMGVR_UViG_3300010032_000012_r__Varidnaviria_k__Bamfordvirae_p__Nucleocytoviricota_c__Megaviricetes_o__Pimascovirales____3300010032_Ga0126336_100208527
IMGVR_UViG_3300010035_000042_r__Varidnaviria_k__Bamfordvirae_p__Preplasmiviricota_c__Polintoviricetes_o__Orthopolintovirales____3300010035_Ga0126343_100083574
DAC81737_1_Stylophora coral adintovirus
MCG8406242_1_Phycisphaerales bacterium
IMGVR_UViG_3300005836_000141_r__Varidnaviria_k__Bamfordvirae_p__Preplasmiviricota_c__Polintoviricetes_o__Orthopolintovirales____3300005836_Ga0074470_117866391
IMGVR_UViG_3300015242_000326_no_taxonomy_3300015242_Ga0137412_100104944
IMGVR_UViG_3300015263_001783_no_taxonomy_3300015263_Ga0183003_100110883
IMGVR_UViG_3300010376_002023_no_taxonomy_3300010376_Ga0126381_1000803852
GIY56757_1_Caerostris darwini
IMGVR_UViG_3300009790_004092_r__Varidnaviria_k__Bamfordvirae_p__Preplasmiviricota_c__Polintoviricetes_o__Orthopolintovirales____3300009790_Ga0115012_100138032
IMGVR_UViG_3300009790_004280_r__Varidnaviria_k__Bamfordvirae_p__Preplasmiviricota_c__Polintoviricetes_o__Orthopolintovirales____3300009790_Ga0115012_100223611
IMGVR_UViG_3300019147_000003_r__Varidnaviria_k__Bamfordvirae_p__Preplasmiviricota_c__Polintoviricetes_o__Orthopolintovirales____3300019147_Ga0193453_10000165
IMGVR_UViG_3300021980_000041_no_taxonomy_3300021980_Ga0232637_100027668
IMGVR_UViG_3300019739_000002_no_taxonomy_3300019739_Ga0194012_100000510
IMGVR_UViG_3300006001_000020_no_taxonomy_3300006001_Ga0081606_10058987
IMGVR_UViG_3300027091_000016_no_taxonomy_3300027091_Ga0209873_100025511
IMGVR_UViG_3300027620_000027_no_taxonomy_3300027620_Ga0209260_100019353
IMGVR_UViG_3300018414_000166_r__Varidnaviria_k__Bamfordvirae_p__Preplasmiviricota_c__Polintoviricetes_o__Orthopolintovirales____3300018414_Ga0194135_100156501
IMGVR_UViG_3300027181_000001_no_taxonomy_3300027181_Ga0208997_10001495
IMGVR_UViG_3300047331_000040_no_taxonomy_3300047331_Ga0495347_0010350_6974_7525
VDH96060_1_Mytilus galloprovincialis
MCK5872763_1_Methylococcales bacterium
ELU00860_1_Capitella teleta
KAG1701166_1_Nymphon striatum
IMGVR_UViG_3300037991_008141_r__Varidnaviria_k__Bamfordvirae_p__Preplasmiviricota_c__Polintoviricetes_o__Orthopolintovirales____3300037991_Ga0183737_006729_4225_4698
2.5

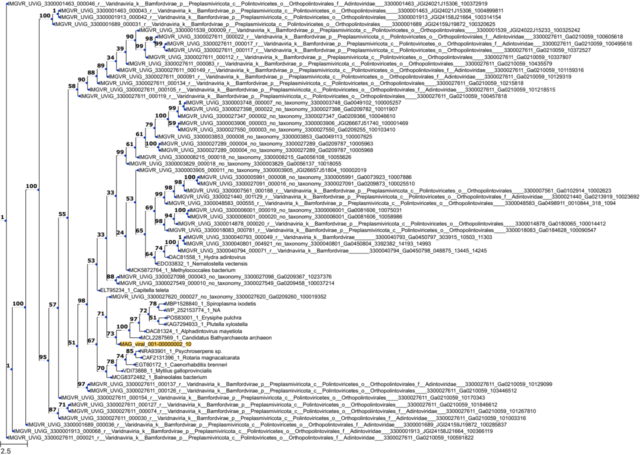
IMGVR_UViG_3300001463_000046_r__Varidnaviria_k__Bamfordvirae_p__Preplasmiviricota_c__Polintoviricetes_o__Orthopolintovirales_f__Adintoviridae___3300001463_JGI24021J15306_100372919
IMGVR_UViG_3300001463_000043_r__Varidnaviria_k__Bamfordvirae_p__Preplasmiviricota_c__Polintoviricetes_o__Orthopolintovirales____3300001463_JGI24021J15306_1004899811
IMGVR_UViG_3300001913_000042_r__Varidnaviria_k__Bamfordvirae_p__Preplasmiviricota_c__Polintoviricetes_o__Orthopolintovirales____3300001913_JGI24158J21664_100314154
IMGVR_UViG_3300001689_000031_r__Varidnaviria_k__Bamfordvirae_p__Preplasmiviricota_c__Polintoviricetes_o__Orthopolintovirales____3300001689_JGI24159J19872_100320625
IMGVR_UViG_3300001539_000009_r__Varidnaviria_k__Bamfordvirae_p__Preplasmiviricota_c__Polintoviricetes_o__Orthopolintovirales____3300001539_JGI24022J15233_100325242
IMGVR_UViG_3300027611_000022_r__Varidnaviria_k__Bamfordvirae_p__Preplasmiviricota_c__Polintoviricetes_o__Orthopolintovirales_f__Adintoviridae___3300027611_Ga0210059_100605618
IMGVR_UViG_3300027611_000017_r__Varidnaviria_k__Bamfordvirae_p__Preplasmiviricota_c__Polintoviricetes_o__Orthopolintovirales_f__Adintoviridae___3300027611_Ga0210059_100495616
IMGVR_UViG_3300027611_000117_r__Varidnaviria_k__Bamfordvirae_p__Preplasmiviricota_c__Polintoviricetes_o__Orthopolintovirales____3300027611_Ga0210059_10372527
IMGVR_UViG_3300027611_000112_r__Varidnaviria_k__Bamfordvirae_p__Preplasmiviricota_c__Polintoviricetes_o__Orthopolintovirales____3300027611_Ga0210059_10337807
IMGVR_UViG_3300027611_000083_r__Varidnaviria_k__Bamfordvirae_p__Preplasmiviricota_c__Polintoviricetes_o__Orthopolintovirales____3300027611_Ga0210059_10435579
IMGVR_UViG_3300027611_000149_r__Varidnaviria_k__Bamfordvirae_p__Preplasmiviricota_c__Polintoviricetes_o__Orthopolintovirales_f__Adintoviridae___3300027611_Ga0210059_101159316
IMGVR_UViG_3300027611_000091_r__Varidnaviria_k__Bamfordvirae_p__Preplasmiviricota_c__Polintoviricetes_o__Orthopolintovirales____3300027611_Ga0210059_10129319
IMGVR_UViG_3300027611_000134_r__Varidnaviria_k__Bamfordvirae_p__Preplasmiviricota_c__Polintoviricetes_o__Orthopolintovirales____3300027611_Ga0210059_10215818
IMGVR_UViG_3300027611_000105_r__Varidnaviria_k__Bamfordvirae_p__Preplasmiviricota_c__Polintoviricetes_o__Orthopolintovirales_f__Adintoviridae___3300027611_Ga0210059_101218515
IMGVR_UViG_3300027611_000119_r__Varidnaviria_k__Bamfordvirae_p__Preplasmiviricota_c__Polintoviricetes_o__Orthopolintovirales____3300027611_Ga0210059_100457818
IMGVR_UViG_3300003748_000007_no_taxonomy_3300003748_Ga0049102_100005257
IMGVR_UViG_3300027398_000022_no_taxonomy_3300027398_Ga0209782_10011907
IMGVR_UViG_3300027347_000002_no_taxonomy_3300027347_Ga0209366_100046610
IMGVR_UViG_3300003906_000003_no_taxonomy_3300003906_JGI26667J51740_100001469
IMGVR_UViG_3300027550_000003_no_taxonomy_3300027550_Ga0209255_100103410
IMGVR_UViG_3300003853_000008_no_taxonomy_3300003853_Ga0049113_100007625
IMGVR_UViG_3300027289_000004_no_taxonomy_3300027289_Ga0209787_10005963
IMGVR_UViG_3300027289_000004_no_taxonomy_3300027289_Ga0209787_10005968
IMGVR_UViG_3300008215_000018_no_taxonomy_3300008215_Ga0056108_10055626
IMGVR_UViG_3300003829_000018_no_taxonomy_3300003829_Ga0056137_10018055
IMGVR_UViG_3300003905_000011_no_taxonomy_3300003905_JGI26657J51804_100002019
IMGVR_UViG_3300005991_000008_no_taxonomy_3300005991_Ga0073923_10007886
IMGVR_UViG_3300027091_000016_no_taxonomy_3300027091_Ga0209873_100025510
IMGVR_UViG_3300007561_000188_r__Varidnaviria_k__Bamfordvirae_p__Preplasmiviricota_c__Polintoviricetes_o__Orthopolintovirales____3300007561_Ga0102914_10002623
IMGVR_UViG_3300021440_001129_r__Varidnaviria_k__Bamfordvirae_p__Preplasmiviricota_c__Polintoviricetes_o__Orthopolintovirales_f__Adintoviridae___3300021440_Ga0213919_10023692
IMGVR_UViG_3300048583_000555_r__Varidnaviria_k__Bamfordvirae_p__Preplasmiviricota_c__Polintoviricetes_o__Orthopolintovirales____3300048583_Ga0498911_0010844_318_1094
IMGVR_UViG_3300006001_000019_no_taxonomy_3300006001_Ga0081606_10075031
IMGVR_UViG_3300006001_000020_no_taxonomy_3300006001_Ga0081606_10058986
IMGVR_UViG_3300014878_000020_r__Varidnaviria_k__Bamfordvirae_p__Preplasmiviricota_c__Polintoviricetes_o__Orthopolintovirales____3300014878_Ga0180065_100014412
IMGVR_UViG_3300018083_000781_r__Varidnaviria_k__Bamfordvirae_p__Preplasmiviricota_c__Polintoviricetes_o__Orthopolintovirales____3300018083_Ga0184628_100090547
IMGVR_UViG_3300040793_000049_r__Varidnaviria_k__Bamfordvirae________3300040793_Ga0450797_303915_10503_11303
IMGVR_UViG_3300040801_004921_no_taxonomy_3300040801_Ga0450804_3392382_14193_14993
IMGVR_UViG_3300040794_000071_r__Varidnaviria_k__Bamfordvirae________3300040794_Ga0450798_048875_13445_14245
DAC81558_1_Hydra adintovirus
EDO33832_1_Nematostella vectensis
MCK5872764_1_Methylococcales bacterium
IMGVR_UViG_3300027098_000043_no_taxonomy_3300027098_Ga0209367_10237376
IMGVR_UViG_3300027549_000010_no_taxonomy_3300027549_Ga0209458_100037214
ELT95234_1_Capitella teleta
IMGVR_UViG_3300027620_000027_no_taxonomy_3300027620_Ga0209260_100019352
MBP1528840_1_Spiroplasma ixodetis
WP_252153774_1_NA
POS83001_1_Erysiphe pulchra
KAG7294933_1_Plutella xylostella
DAC81324_1_Alphadintovirus mayetiola
MCL2287569_1_Candidatus Bathyarchaeota archaeon
MAG_viral_001-00000002_10
NRA93901_1_Psychroserpens sp.
CAF2131396_1_Rotaria magnacalcarata
EGT60172_1_Caenorhabditis brenneri
VDI73888_1_Mytilus galloprovincialis
MCG8372482_1_Balneolales bacterium
IMGVR_UViG_3300027611_000137_r__Varidnaviria_k__Bamfordvirae_p__Preplasmiviricota_c__Polintoviricetes_o__Orthopolintovirales_f__Adintoviridae___3300027611_Ga0210059_10129099
IMGVR_UViG_3300027611_000126_r__Varidnaviria_k__Bamfordvirae_p__Preplasmiviricota_c__Polintoviricetes_o__Orthopolintovirales____3300027611_Ga0210059_103446512
IMGVR_UViG_3300027611_000154_r__Varidnaviria_k__Bamfordvirae_p__Preplasmiviricota_c__Polintoviricetes_o__Orthopolintovirales____3300027611_Ga0210059_10170343
IMGVR_UViG_3300027611_000127_r__Varidnaviria_k__Bamfordvirae_p__Preplasmiviricota_c__Polintoviricetes_o__Orthopolintovirales____3300027611_Ga0210059_101846612
IMGVR_UViG_3300027611_000074_r__Varidnaviria_k__Bamfordvirae_p__Preplasmiviricota_c__Polintoviricetes_o__Orthopolintovirales_f__Adintoviridae___3300027611_Ga0210059_101267810
IMGVR_UViG_3300027611_000030_r__Varidnaviria_k__Bamfordvirae_p__Preplasmiviricota_c__Polintoviricetes_o__Orthopolintovirales_f__Adintoviridae___3300027611_Ga0210059_101003316
IMGVR_UViG_3300001689_000036_r__Varidnaviria_k__Bamfordvirae_p__Preplasmiviricota_c__Polintoviricetes_o__Orthopolintovirales_f__Adintoviridae___3300001689_JGI24159J19872_100285837
IMGVR_UViG_3300001913_000068_r__Varidnaviria_k__Bamfordvirae_p__Preplasmiviricota_c__Polintoviricetes_o__Orthopolintovirales_f__Adintoviridae___3300001913_JGI24158J21664_100366119
IMGVR_UViG_3300027611_000021_r__Varidnaviria_k__Bamfordvirae_p__Preplasmiviricota_c__Polintoviricetes_o__Orthopolintovirales_f__Adintoviridae___3300027611_Ga0210059_100591822
1
100
90
98
99
39
99
34
89
88
90
58
1
99
100
59
79
61
100
61
99
100
98
98
99
53
100
94
69
98
24
65
1
100
74
64
33
33
88
53
100
78
72
97
51
71
100
73
67
85
74
78
70
98
97
51
57
55
100
71
87
95
1
100
1
2.5

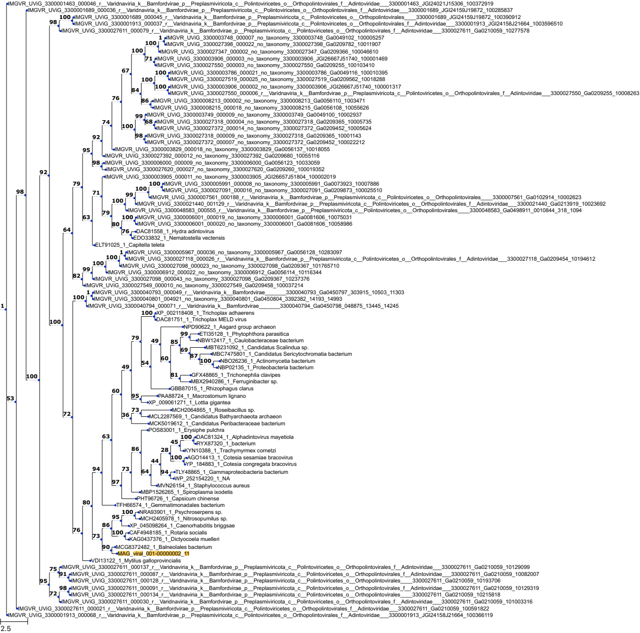
IMGVR_UViG_3300001463_000046_r__Varidnaviria_k__Bamfordvirae_p__Preplasmiviricota_c__Polintoviricetes_o__Orthopolintovirales_f__Adintoviridae___3300001463_JGI24021J15306_100372919
IMGVR_UViG_3300001689_000036_r__Varidnaviria_k__Bamfordvirae_p__Preplasmiviricota_c__Polintoviricetes_o__Orthopolintovirales_f__Adintoviridae___3300001689_JGI24159J19872_100285837
IMGVR_UViG_3300001689_000045_r__Varidnaviria_k__Bamfordvirae_p__Preplasmiviricota_c__Polintoviricetes_o__Orthopolintovirales____3300001689_JGI24159J19872_100390912
IMGVR_UViG_3300001913_000037_r__Varidnaviria_k__Bamfordvirae_p__Preplasmiviricota_c__Polintoviricetes_o__Orthopolintovirales_f__Adintoviridae___3300001913_JGI24158J21664_1003596510
IMGVR_UViG_3300027611_000079_r__Varidnaviria_k__Bamfordvirae_p__Preplasmiviricota_c__Polintoviricetes_o__Orthopolintovirales_f__Adintoviridae___3300027611_Ga0210059_10277578
IMGVR_UViG_3300003748_000007_no_taxonomy_3300003748_Ga0049102_100005257
IMGVR_UViG_3300027398_000022_no_taxonomy_3300027398_Ga0209782_10011907
IMGVR_UViG_3300027347_000002_no_taxonomy_3300027347_Ga0209366_100046610
IMGVR_UViG_3300003906_000003_no_taxonomy_3300003906_JGI26667J51740_100001469
IMGVR_UViG_3300027550_000003_no_taxonomy_3300027550_Ga0209255_100103410
IMGVR_UViG_3300003786_000021_no_taxonomy_3300003786_Ga0049116_100010395
IMGVR_UViG_3300027519_000025_no_taxonomy_3300027519_Ga0209562_10018288
IMGVR_UViG_3300003906_000002_no_taxonomy_3300003906_JGI26667J51740_100001317
IMGVR_UViG_3300027550_000006_r__Varidnaviria_k__Bamfordvirae_p__Preplasmiviricota_c__Polintoviricetes_o__Orthopolintovirales_f__Adintoviridae___3300027550_Ga0209255_10008263
IMGVR_UViG_3300008213_000002_no_taxonomy_3300008213_Ga0056110_1003471
IMGVR_UViG_3300008215_000018_no_taxonomy_3300008215_Ga0056108_10055626
IMGVR_UViG_3300003749_000009_no_taxonomy_3300003749_Ga0049100_10002937
IMGVR_UViG_3300027318_000004_no_taxonomy_3300027318_Ga0209365_10005735
IMGVR_UViG_3300027372_000014_no_taxonomy_3300027372_Ga0209452_10005624
IMGVR_UViG_3300027318_000009_no_taxonomy_3300027318_Ga0209365_10001143
IMGVR_UViG_3300027372_000007_no_taxonomy_3300027372_Ga0209452_100022212
IMGVR_UViG_3300003829_000018_no_taxonomy_3300003829_Ga0056137_10018055
IMGVR_UViG_3300027392_000012_no_taxonomy_3300027392_Ga0209680_10055116
IMGVR_UViG_3300006000_000009_no_taxonomy_3300006000_Ga0056123_10033059
IMGVR_UViG_3300027620_000027_no_taxonomy_3300027620_Ga0209260_100019352
IMGVR_UViG_3300003905_000011_no_taxonomy_3300003905_JGI26657J51804_100002019
IMGVR_UViG_3300005991_000008_no_taxonomy_3300005991_Ga0073923_10007886
IMGVR_UViG_3300027091_000016_no_taxonomy_3300027091_Ga0209873_100025510
IMGVR_UViG_3300007561_000188_r__Varidnaviria_k__Bamfordvirae_p__Preplasmiviricota_c__Polintoviricetes_o__Orthopolintovirales____3300007561_Ga0102914_10002623
IMGVR_UViG_3300021440_001129_r__Varidnaviria_k__Bamfordvirae_p__Preplasmiviricota_c__Polintoviricetes_o__Orthopolintovirales_f__Adintoviridae___3300021440_Ga0213919_10023692
IMGVR_UViG_3300048583_000555_r__Varidnaviria_k__Bamfordvirae_p__Preplasmiviricota_c__Polintoviricetes_o__Orthopolintovirales____3300048583_Ga0498911_0010844_318_1094
IMGVR_UViG_3300006001_000019_no_taxonomy_3300006001_Ga0081606_10075031
IMGVR_UViG_3300006001_000020_no_taxonomy_3300006001_Ga0081606_10058986
DAC81558_1_Hydra adintovirus
EDO33832_1_Nematostella vectensis
ELT91025_1_Capitella teleta
IMGVR_UViG_3300005967_000036_no_taxonomy_3300005967_Ga0056128_10283097
IMGVR_UViG_3300027118_000026_r__Varidnaviria_k__Bamfordvirae_p__Preplasmiviricota_c__Polintoviricetes_o__Orthopolintovirales_f__Adintoviridae___3300027118_Ga0209454_10194612
IMGVR_UViG_3300027098_000023_no_taxonomy_3300027098_Ga0209367_101765710
IMGVR_UViG_3300006912_000022_no_taxonomy_3300006912_Ga0056114_10116344
IMGVR_UViG_3300027098_000043_no_taxonomy_3300027098_Ga0209367_10237376
IMGVR_UViG_3300027549_000010_no_taxonomy_3300027549_Ga0209458_100037214
IMGVR_UViG_3300040793_000049_r__Varidnaviria_k__Bamfordvirae_______3300040793_Ga0450797_303915_10503_11303
IMGVR_UViG_3300040801_004921_no_taxonomy_3300040801_Ga0450804_3392382_14193_14993
IMGVR_UViG_3300040794_000071_r__Varidnaviria_k__Bamfordvirae_______3300040794_Ga0450798_048875_13445_14245
XP_002118408_1_Trichoplax adhaerens
DAC81751_1_Trichoplax MELD virus
NPD90622_1_Asgard group archaeon
ETI35128_1_Phytophthora parasitica
NBW12417_1_Caulobacteraceae bacterium
MBT6231092_1_Candidatus Scalindua sp.
MBC7475801_1_Candidatus Sericytochromatia bacterium
NBO26236_1_Actinomycetia bacterium
NBP02135_1_Proteobacteria bacterium
GFX48865_1_Trichonephila clavipes
MBX2940286_1_Ferruginibacter sp.
GBB87015_1_Rhizophagus clarus
PAA88724_1_Macrostomum lignano
XP_009061271_1_Lottia gigantea
MCH2064865_1_Roseibacillus sp.
MCL2287569_1_Candidatus Bathyarchaeota archaeon
MCK5019612_1_Candidatus Peribacteraceae bacterium
POS83001_1_Erysiphe pulchra
DAC81324_1_Alphadintovirus mayetiola
RYX87320_1_bacterium
KYN10388_1_Trachymyrmex cornetzi
AGO14413_1_Cotesia sesamiae bracovirus
YP_184883_1_Cotesia congregata bracovirus
TLY48865_1_Gammaproteobacteria bacterium
WP_252154220_1_NA
MVN26154_1_Staphylococcus aureus
MBP1526265_1_Spiroplasma ixodetis
PHT96726_1_Capsicum chinense
TFH66574_1_Gemmatimonadales bacterium
NRA93901_1_Psychroserpens sp.
MCH2405978_1_Nitrosopumilus sp.
XP_045098264_1_Caenorhabditis briggsae
CAF4948185_1_Rotaria socialis
KAG0437376_1_Dictyocoela muelleri
MCG8372482_1_Balneolales bacterium
MAG_viral_001-00000002_11
VDI13122_1_Mytilus galloprovincialis
IMGVR_UViG_3300027611_000137_r__Varidnaviria_k__Bamfordvirae_p__Preplasmiviricota_c__Polintoviricetes_o__Orthopolintovirales_f__Adintoviridae___3300027611_Ga0210059_10129099
IMGVR_UViG_3300027611_000087_r__Varidnaviria_k__Bamfordvirae_p__Preplasmiviricota_c__Polintoviricetes_o__Orthopolintovirales_f__Adintoviridae___3300027611_Ga0210059_10082007
IMGVR_UViG_3300027611_000128_r__Varidnaviria_k__Bamfordvirae_p__Preplasmiviricota_c__Polintoviricetes_o__Orthopolintovirales____3300027611_Ga0210059_10193706
IMGVR_UViG_3300027611_000091_r__Varidnaviria_k__Bamfordvirae_p__Preplasmiviricota_c__Polintoviricetes_o__Orthopolintovirales_f__Adintoviridae___3300027611_Ga0210059_10129319
IMGVR_UViG_3300027611_000134_r__Varidnaviria_k__Bamfordvirae_p__Preplasmiviricota_c__Polintoviricetes_o__Orthopolintovirales____3300027611_Ga0210059_10215818
IMGVR_UViG_3300027611_000030_r__Varidnaviria_k__Bamfordvirae_p__Preplasmiviricota_c__Polintoviricetes_o__Orthopolintovirales_f__Adintoviridae___3300027611_Ga0210059_101003316
IMGVR_UViG_3300027611_000021_r__Varidnaviria_k__Bamfordvirae_p__Preplasmiviricota_c__Polintoviricetes_o__Orthopolintovirales_f__Adintoviridae___3300027611_Ga0210059_100591822
IMGVR_UViG_3300001913_000068_r__Varidnaviria_k__Bamfordvirae_p__Preplasmiviricota_c__Polintoviricetes_o__Orthopolintovirales_f__Adintoviridae___3300001913_JGI24158J21664_100366119
2.5

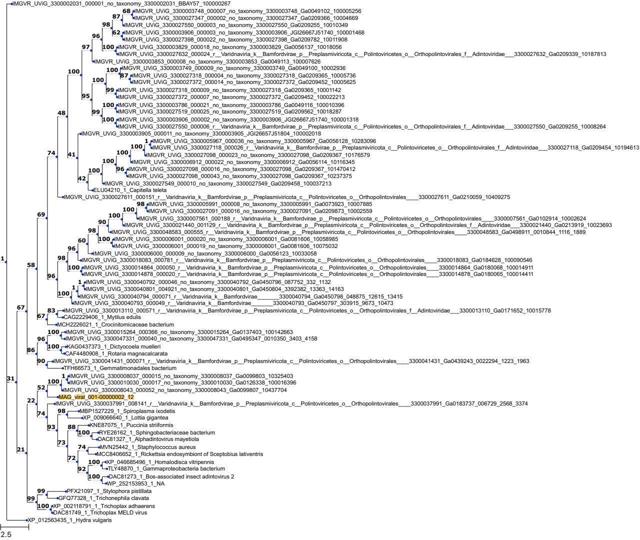
IMGVR_UViG_3300002031_000001_no_taxonomy_3300002031_BBAY57_100000267
IMGVR_UViG_3300003748_000007_no_taxonomy_3300003748_Ga0049102_100005256
IMGVR_UViG_3300027347_000002_no_taxonomy_3300027347_Ga0209366_10004669
IMGVR_UViG_3300027550_000003_no_taxonomy_3300027550_Ga0209255_10010349
IMGVR_UViG_3300003906_000003_no_taxonomy_3300003906_JGI26667J51740_100001468
IMGVR_UViG_3300027398_000022_no_taxonomy_3300027398_Ga0209782_10011908
IMGVR_UViG_3300003829_000018_no_taxonomy_3300003829_Ga0056137_10018056
IMGVR_UViG_3300027632_000024_r__Varidnaviria_k__Bamfordvirae_p__Preplasmiviricota_c__Polintoviricetes_o__Orthopolintovirales_f__Adintoviridae___3300027632_Ga0209339_10187813
IMGVR_UViG_3300003853_000008_no_taxonomy_3300003853_Ga0049113_100007626
IMGVR_UViG_3300003749_000009_no_taxonomy_3300003749_Ga0049100_10002936
IMGVR_UViG_3300027318_000004_no_taxonomy_3300027318_Ga0209365_10005736
IMGVR_UViG_3300027372_000014_no_taxonomy_3300027372_Ga0209452_10005625
IMGVR_UViG_3300027318_000009_no_taxonomy_3300027318_Ga0209365_10001142
IMGVR_UViG_3300027372_000007_no_taxonomy_3300027372_Ga0209452_100022213
IMGVR_UViG_3300003786_000021_no_taxonomy_3300003786_Ga0049116_100010396
IMGVR_UViG_3300027519_000025_no_taxonomy_3300027519_Ga0209562_10018287
IMGVR_UViG_3300003906_000002_no_taxonomy_3300003906_JGI26667J51740_100001318
IMGVR_UViG_3300027550_000006_r__Varidnaviria_k__Bamfordvirae_p__Preplasmiviricota_c__Polintoviricetes_o__Orthopolintovirales_f__Adintoviridae___3300027550_Ga0209255_10008264
IMGVR_UViG_3300003905_000011_no_taxonomy_3300003905_JGI26657J51804_100002018
IMGVR_UViG_3300005967_000036_no_taxonomy_3300005967_Ga0056128_10283096
IMGVR_UViG_3300027118_000026_r__Varidnaviria_k__Bamfordvirae_p__Preplasmiviricota_c__Polintoviricetes_o__Orthopolintovirales_f__Adintoviridae___3300027118_Ga0209454_10194613
IMGVR_UViG_3300027098_000023_no_taxonomy_3300027098_Ga0209367_10176579
IMGVR_UViG_3300006912_000022_no_taxonomy_3300006912_Ga0056114_10116345
IMGVR_UViG_3300027098_000016_no_taxonomy_3300027098_Ga0209367_101470412
IMGVR_UViG_3300027098_000043_no_taxonomy_3300027098_Ga0209367_10237375
IMGVR_UViG_3300027549_000010_no_taxonomy_3300027549_Ga0209458_100037213
ELU04210_1_Capitella teleta
IMGVR_UViG_3300027611_000151_r__Varidnaviria_k__Bamfordvirae_p__Preplasmiviricota_c__Polintoviricetes_o__Orthopolintovirales____3300027611_Ga0210059_10409275
IMGVR_UViG_3300005991_000008_no_taxonomy_3300005991_Ga0073923_10007885
IMGVR_UViG_3300027091_000016_no_taxonomy_3300027091_Ga0209873_10002559
IMGVR_UViG_3300007561_000188_r__Varidnaviria_k__Bamfordvirae_p__Preplasmiviricota_c__Polintoviricetes_o__Orthopolintovirales____3300007561_Ga0102914_10002624
IMGVR_UViG_3300021440_001129_r__Varidnaviria_k__Bamfordvirae_p__Preplasmiviricota_c__Polintoviricetes_o__Orthopolintovirales_f__Adintoviridae___3300021440_Ga0213919_10023693
IMGVR_UViG_3300048583_000555_r__Varidnaviria_k__Bamfordvirae_p__Preplasmiviricota_c__Polintoviricetes_o__Orthopolintovirales____3300048583_Ga0498911_0010844_1116_1889
IMGVR_UViG_3300006001_000020_no_taxonomy_3300006001_Ga0081606_10058985
IMGVR_UViG_3300006001_000019_no_taxonomy_3300006001_Ga0081606_10075032
IMGVR_UViG_3300006000_000009_no_taxonomy_3300006000_Ga0056123_10033058
IMGVR_UViG_3300018083_000781_r__Varidnaviria_k__Bamfordvirae_p__Preplasmiviricota_c__Polintoviricetes_o__Orthopolintovirales____3300018083_Ga0184628_100090546
IMGVR_UViG_3300014864_000050_r__Varidnaviria_k__Bamfordvirae_p__Preplasmiviricota_c__Polintoviricetes_o__Orthopolintovirales____3300014864_Ga0180068_100014911
IMGVR_UViG_3300014878_000020_r__Varidnaviria_k__Bamfordvirae_p__Preplasmiviricota_c__Polintoviricetes_o__Orthopolintovirales____3300014878_Ga0180065_100014411
IMGVR_UViG_3300040792_000046_no_taxonomy_3300040792_Ga0450796_087752_332_1132
IMGVR_UViG_3300040801_004921_no_taxonomy_3300040801_Ga0450804_3392382_13363_14163
IMGVR_UViG_3300040794_000071_r__Varidnaviria_k__Bamfordvirae_______3300040794_Ga0450798_048875_12615_13415
IMGVR_UViG_3300040793_000049_r__Varidnaviria_k__Bamfordvirae_______3300040793_Ga0450797_303915_9673_10473
IMGVR_UViG_3300013110_000571_r__Varidnaviria_k__Bamfordvirae_p__Preplasmiviricota_c__Polintoviricetes_o__Orthopolintovirales_f__Adintoviridae___3300013110_Ga0171652_10015778
CAG2229406_1_Mytilus edulis
MCH2226021_1_Crocinitomicaceae bacterium
IMGVR_UViG_3300015264_000366_no_taxonomy_3300015264_Ga0137403_100142663
IMGVR_UViG_3300047331_000040_no_taxonomy_3300047331_Ga0495347_0010350_3403_4158
KAG0437373_1_Dictyocoela muelleri
CAF4480908_1_Rotaria magnacalcarata
IMGVR_UViG_3300041431_000071_r__Varidnaviria_k__Bamfordvirae_p__Preplasmiviricota_c__Polintoviricetes_o__Orthopolintovirales____3300041431_Ga0439243_0022294_1223_1963
TFH66573_1_Gemmatimonadales bacterium
IMGVR_UViG_3300008037_000015_no_taxonomy_3300008037_Ga0099803_10325403
IMGVR_UViG_3300010030_000017_no_taxonomy_3300010030_Ga0126338_100016396
IMGVR_UViG_3300008043_000052_no_taxonomy_3300008043_Ga0099807_10437704
MAG_viral_001-00000002_12
IMGVR_UViG_3300037991_008141_r__Varidnaviria_k__Bamfordvirae_p__Preplasmiviricota_c__Polintoviricetes_o__Orthopolintovirales____3300037991_Ga0183737_006729_2568_3374
MBP1527229_1_Spiroplasma ixodetis
XP_009066640_1_Lottia gigantea
KNE87075_1_Puccinia striiformis
RYE26162_1_Sphingobacteriaceae bacterium
DAC81327_1_Alphadintovirus mayetiola
MVN25442_1_Staphylococcus aureus
MCC8406652_1_Rickettsia endosymbiont of Sceptobius lativentris
XP_046685496_1_Homalodisca vitripennis
TLY48870_1_Gammaproteobacteria bacterium
DAC81273_1_Bos-associated insect adintovirus 2
WP_252153953_1_NA
PFX21097_1_Stylophora pistillata
GFQ77328_1_Trichonephila clavata
XP_002118791_1_Trichoplax adhaerens
DAC81749_1_Trichoplax MELD virus
XP_012563435_1_Hydra vulgaris
2.5

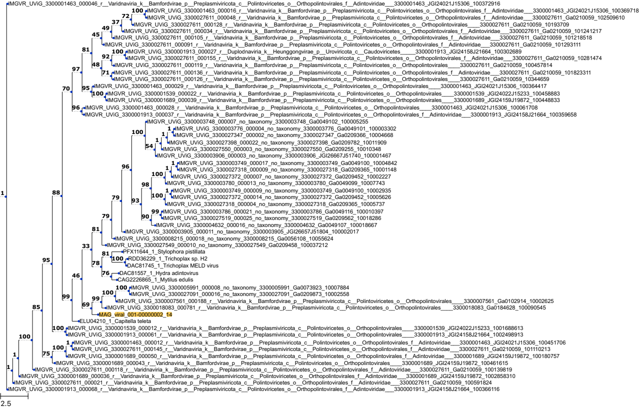

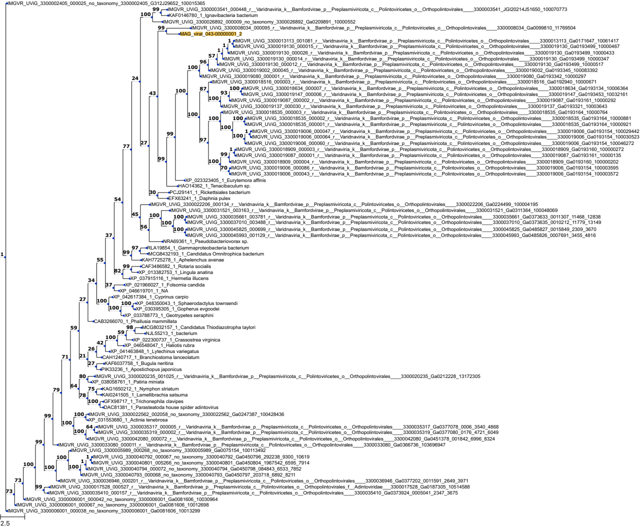
IMGVR_UViG_3300002405_000025_no_taxonomy_3300002405_G312J29652_100015365
IMGVR_UViG_3300003541_000448_r__Varidnaviria_k__Bamfordvirae_p__Preplasmiviricota_c__Polintoviricetes_o__Orthopolintovirales____3300003541_JGI20214J51650_100070773
KAF0146780_1_Ignavibacteria bacterium
IMGVR_UViG_3300026892_000009_no_taxonomy_3300026892_Ga0209891_10000552
IMGVR_UViG_3300008034_000095_r__Varidnaviria_k__Bamfordvirae_p__Preplasmiviricota_c__Polintoviricetes_o__Orthopolintovirales____3300008034_Ga0099810_11769504
MAG_viral_043-00000001_2
IMGVR_UViG_3300013113_001081_r__Varidnaviria_k__Bamfordvirae_p__Preplasmiviricota_c__Polintoviricetes_o__Orthopolintovirales____3300013113_Ga0171647_10061417
IMGVR_UViG_3300019130_000015_r__Varidnaviria_k__Bamfordvirae_p__Preplasmiviricota_c__Polintoviricetes_o__Orthopolintovirales____3300019130_Ga0193499_10000467
IMGVR_UViG_3300019130_000026_r__Varidnaviria_k__Bamfordvirae_p__Preplasmiviricota_c__Polintoviricetes_o__Orthopolintovirales____3300019130_Ga0193499_10000433
IMGVR_UViG_3300019130_000014_r__Varidnaviria_k__Bamfordvirae_p__Preplasmiviricota_c__Polintoviricetes_o__Orthopolintovirales____3300019130_Ga0193499_10000347
IMGVR_UViG_3300019130_000012_r__Varidnaviria_k__Bamfordvirae_p__Preplasmiviricota_c__Polintoviricetes_o__Orthopolintovirales____3300019130_Ga0193499_10000517
IMGVR_UViG_3300019002_000045_r__Varidnaviria_k__Bamfordvirae_p__Preplasmiviricota_c__Polintoviricetes_o__Orthopolintovirales____3300019002_Ga0193345_100083392
IMGVR_UViG_3300019080_000001_r__Varidnaviria_k__Bamfordvirae_p__Preplasmiviricota_c__Polintoviricetes_o__Orthopolintovirales____3300019080_Ga0193342_10000297
IMGVR_UViG_3300018516_000003_r__Varidnaviria_k__Bamfordvirae_p__Preplasmiviricota_c__Polintoviricetes_o__Orthopolintovirales____3300018516_Ga0192940_10000671
IMGVR_UViG_3300018634_000007_r__Varidnaviria_k__Bamfordvirae_p__Preplasmiviricota_c__Polintoviricetes_o__Orthopolintovirales____3300018634_Ga0193134_10006364
IMGVR_UViG_3300019147_000006_r__Varidnaviria_k__Bamfordvirae_p__Preplasmiviricota_c__Polintoviricetes_o__Orthopolintovirales____3300019147_Ga0193453_10032161
IMGVR_UViG_3300019087_000002_r__Varidnaviria_k__Bamfordvirae_p__Preplasmiviricota_c__Polintoviricetes_o__Orthopolintovirales____3300019087_Ga0193161_10000292
IMGVR_UViG_3300019137_000030_r__Varidnaviria_k__Bamfordvirae_p__Preplasmiviricota_c__Polintoviricetes_o__Orthopolintovirales____3300019137_Ga0193321_10003643
IMGVR_UViG_3300018535_000003_r__Varidnaviria_k__Bamfordvirae_p__Preplasmiviricota_c__Polintoviricetes_o__Orthopolintovirales____3300018535_Ga0193164_10000764
IMGVR_UViG_3300018535_000002_r__Varidnaviria_k__Bamfordvirae_p__Preplasmiviricota_c__Polintoviricetes_o__Orthopolintovirales____3300018535_Ga0193164_10000881
IMGVR_UViG_3300018535_000001_r__Varidnaviria_k__Bamfordvirae_p__Preplasmiviricota_c__Polintoviricetes_o__Orthopolintovirales____3300018535_Ga0193164_10000921
IMGVR_UViG_3300019006_000047_r__Varidnaviria_k__Bamfordvirae_p__Preplasmiviricota_c__Polintoviricetes_o__Orthopolintovirales____3300019006_Ga0193154_100029442
IMGVR_UViG_3300019006_000064_r__Varidnaviria_k__Bamfordvirae_p__Preplasmiviricota_c__Polintoviricetes_o__Orthopolintovirales____3300019006_Ga0193154_100030523
IMGVR_UViG_3300019006_000060_r__Varidnaviria_k__Bamfordvirae_p__Preplasmiviricota_c__Polintoviricetes_o__Orthopolintovirales____3300019006_Ga0193154_100046272
IMGVR_UViG_3300018909_000003_r__Varidnaviria_k__Bamfordvirae_p__Preplasmiviricota_c__Polintoviricetes_o__Orthopolintovirales____3300018909_Ga0193160_100000272
IMGVR_UViG_3300019087_000001_r__Varidnaviria_k__Bamfordvirae_p__Preplasmiviricota_c__Polintoviricetes_o__Orthopolintovirales____3300019087_Ga0193161_10000135
IMGVR_UViG_3300018909_000004_r__Varidnaviria_k__Bamfordvirae_p__Preplasmiviricota_c__Polintoviricetes_o__Orthopolintovirales____3300018909_Ga0193160_10000202
IMGVR_UViG_3300019006_000086_r__Varidnaviria_k__Bamfordvirae_p__Preplasmiviricota_c__Polintoviricetes_o__Orthopolintovirales____3300019006_Ga0193154_100003595
IMGVR_UViG_3300019006_000043_r__Varidnaviria_k__Bamfordvirae_p__Preplasmiviricota_c__Polintoviricetes_o__Orthopolintovirales____3300019006_Ga0193154_100003572
XP_023323405_1_Eurytemora affinis
HAO14362_1_Tenacibaculum sp.
PCJ29141_1_Rickettsiales bacterium
EFX63241_1_Daphnia pulex
IMGVR_UViG_3300022206_000134_r__Varidnaviria_k__Bamfordvirae_p__Preplasmiviricota_c__Polintoviricetes_o__Orthopolintovirales____3300022206_Ga0224499_100004195
IMGVR_UViG_3300031521_003163_r__Varidnaviria_k__Bamfordvirae_p__Preplasmiviricota_c__Polintoviricetes_o__Orthopolintovirales____3300031521_Ga0311364_100048069
IMGVR_UViG_3300035661_003781_r__Varidnaviria_k__Bamfordvirae_p__Preplasmiviricota_c__Polintoviricetes_o__Orthopolintovirales____3300035661_Ga0373633_0011307_11468_12838
IMGVR_UViG_3300037010_003488_r__Varidnaviria_k__Bamfordvirae_p__Preplasmiviricota_c__Polintoviricetes_o__Orthopolintovirales____3300037010_Ga0373635_0010212_11779_13149
IMGVR_UViG_3300045825_000699_r__Varidnaviria_k__Bamfordvirae_p__Preplasmiviricota_c__Polintoviricetes_o__Orthopolintovirales____3300045825_Ga0485827_0015849_2309_3670
IMGVR_UViG_3300045993_001129_r__Varidnaviria_k__Bamfordvirae_p__Preplasmiviricota_c__Polintoviricetes_o__Orthopolintovirales____3300045993_Ga0485826_0007691_3455_4816
NRA69361_1_Pseudobacteriovorax sp.
RLA19854_1_Gammaproteobacteria bacterium
MCG8432193_1_Candidatus Omnitrophica bacterium
KAH7725278_1_Aphelenchus avenae
CAF3486582_1_Rotaria socialis
XP_013382753_1_Lingula anatina
XP_037915116_1_Hermetia illucens
XP_021966027_1_Folsomia candida
XP_046619701_1_NA
XP_042617384_1_Cyprinus carpio
XP_048350043_1_Sphaerodactylus townsendi
XP_030395305_1_Gopherus evgoodei
XP_033788773_1_Geotrypetes seraphini
CAB3266070_1_Phallusia mammillata
MCG8032157_1_Candidatus Thiodiazotropha taylori
NJL55213_1_bacterium
XP_022300737_1_Crassostrea virginica
XP_046548047_1_Haliotis rubra
XP_041463848_1_Lytechinus variegatus
CAH1240717_1_Branchiostoma lanceolatum
KAF6037758_1_Bugula neritina
PIK33236_1_Apostichopus japonicus
IMGVR_UViG_3300020235_001025_r__Varidnaviria_k__Bamfordvirae_p__Preplasmiviricota_c__Polintoviricetes_o__Orthopolintovirales____3300020235_Ga0212228_13172305
XP_038058761_1_Patiria miniata
KAG1650212_1_Nymphon striatum
KAI0241505_1_Lamellibrachia satsuma
GFX98717_1_Trichonephila clavipes
DAC81381_1_Parasteatoda house spider adintovirus
IMGVR_UViG_3300022562_002058_no_taxonomy_3300022562_Ga0247387_100428436
XP_031553680_1_Actinia tenebrosa
IMGVR_UViG_3300035317_000005_r__Varidnaviria_k__Bamfordvirae_p__Preplasmiviricota_c__Polintoviricetes_o__Orthopolintovirales____3300035317_Ga0377078_0006_3540_4868
IMGVR_UViG_3300035319_000002_r__Varidnaviria_k__Bamfordvirae_p__Preplasmiviricota_c__Polintoviricetes_o__Orthopolintovirales____3300035319_Ga0377080_0176_4721_6049
IMGVR_UViG_3300042080_000072_r__Varidnaviria_k__Bamfordvirae_p__Preplasmiviricota_c__Polintoviricetes_o__Orthopolintovirales____3300042080_Ga0451378_001842_6996_8324
IMGVR_UViG_3300033080_000011_r__Varidnaviria_k__Bamfordvirae_p__Preplasmiviricota_c__Polintoviricetes_o__Orthopolintovirales____3300033080_Ga0366736_103696947
IMGVR_UViG_3300005989_000268_no_taxonomy_3300005989_Ga0075154_100113492
IMGVR_UViG_3300040792_000067_no_taxonomy_3300040792_Ga0450796_292238_9300_10619
IMGVR_UViG_3300040801_005266_no_taxonomy_3300040801_Ga0450804_1967542_6595_7914
IMGVR_UViG_3300040794_000072_no_taxonomy_3300040794_Ga0450798_084843_6533_7852
IMGVR_UViG_3300040793_000068_no_taxonomy_3300040793_Ga0450797_203718_6892_8211
IMGVR_UViG_3300036946_000201_r__Varidnaviria_k__Bamfordvirae_p__Preplasmiviricota_c__Polintoviricetes_o__Orthopolintovirales____3300036946_Ga0377202_0011591_2649_3971
IMGVR_UViG_3300017528_000527_r__Varidnaviria_k__Bamfordvirae_p__Preplasmiviricota_c__Polintoviricetes_o__Orthopolintovirales_f__Adintoviridae____3300017528_Ga0187305_10514588
IMGVR_UViG_3300035410_000157_r__Varidnaviria_k__Bamfordvirae_p__Preplasmiviricota_c__Polintoviricetes_o__Orthopolintovirales____3300035410_Ga0373924_0005041_2347_3675
IMGVR_UViG_3300006001_000042_no_taxonomy_3300006001_Ga0081606_10030964
IMGVR_UViG_3300006001_000067_no_taxonomy_3300006001_Ga0081606_10012698
IMGVR_UViG_3300006001_000038_no_taxonomy_3300006001_Ga0081606_10013299
2.5

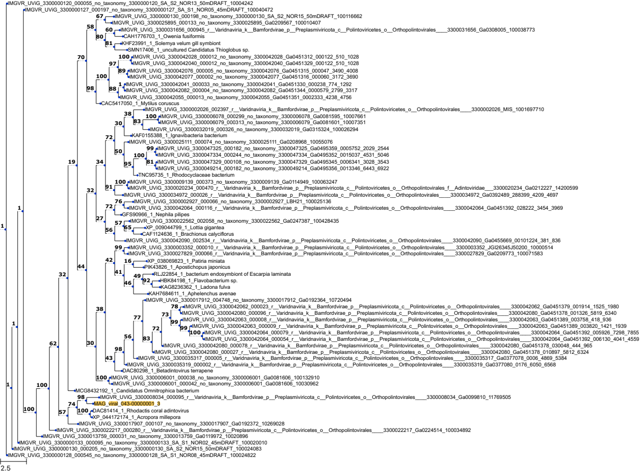

IMGVR_UViG_3300000120_000055_no_taxonomy_3300000120_SA_S2_NOR13_50mDRAFT_10004242
IMGVR_UViG_3300000127_000197_no_taxonomy_3300000127_SA_S1_NOR05_45mDRAFT_100040472
IMGVR_UViG_3300000130_000198_no_taxonomy_3300000130_SA_S2_NOR15_50mDRAFT_100116662
IMGVR_UViG_3300025895_000133_no_taxonomy_3300025895_Ga0209567_100010407
IMGVR_UViG_3300031656_000945_r__Varidnaviria_k__Bamfordvirae_p__Preplasmiviricota_c__Polintoviricetes_o__Orthopolintovirales____3300031656_Ga0308005_100038773
CAH1776703_1_Owenia fusiformis
KHF23991_1_Solemya velum gill symbiont
SMN17406_1_uncultured Candidatus Thioglobus sp.
IMGVR_UViG_3300042028_000012_no_taxonomy_3300042028_Ga0451312_000122_510_1028
IMGVR_UViG_3300042040_000012_no_taxonomy_3300042040_Ga0451329_000122_510_1028
IMGVR_UViG_3300042076_000005_no_taxonomy_3300042076_Ga0451315_000047_3490_4008
IMGVR_UViG_3300042077_000002_no_taxonomy_3300042077_Ga0451316_000060_3172_3690
IMGVR_UViG_3300042041_000003_no_taxonomy_3300042041_Ga0451330_000238_774_1292
IMGVR_UViG_3300042082_000004_no_taxonomy_3300042082_Ga0451344_000579_2799_3317
IMGVR_UViG_3300042055_000013_no_taxonomy_3300042055_Ga0451351_000233_4238_4756
CAC5417050_1_Mytilus coruscus
IMGVR_UViG_3300002026_002397_r__Varidnaviria_k__Bamfordvirae_p__Preplasmiviricota_c__Polintoviricetes_o__Orthopolintovirales____3300002026_MIS_1001697710
IMGVR_UViG_3300006078_000299_no_taxonomy_3300006078_Ga0081595_10007661
IMGVR_UViG_3300006079_000313_no_taxonomy_3300006079_Ga0081601_10007351
IMGVR_UViG_3300032019_000326_no_taxonomy_3300032019_Ga0315324_100126294
KAF0155388_1_Ignavibacteria bacterium
IMGVR_UViG_3300025111_000074_no_taxonomy_3300025111_Ga0208968_10055076
IMGVR_UViG_3300047325_000182_no_taxonomy_3300047325_Ga0495359_0005752_2029_2544
IMGVR_UViG_3300047334_000244_no_taxonomy_3300047334_Ga0495352_0015037_4531_5046
IMGVR_UViG_3300047329_000108_no_taxonomy_3300047329_Ga0495345_0006341_3028_3543
IMGVR_UViG_3300049214_000182_no_taxonomy_3300049214_Ga0495356_0013346_6443_6922
TNC95735_1_Rhodocyclaceae bacterium
IMGVR_UViG_3300009139_000373_no_taxonomy_3300009139_Ga0114949_100063247
IMGVR_UViG_3300020234_000470_r__Varidnaviria_k__Bamfordvirae_p__Preplasmiviricota_c__Polintoviricetes_o__Orthopolintovirales_f__Adintoviridae____3300020234_Ga0212227_14200599
IMGVR_UViG_3300034972_000026_r__Varidnaviria_k__Bamfordvirae_p__Preplasmiviricota_c__Polintoviricetes_o__Orthopolintovirales____3300034972_Ga0392489_288399_4209_4697
IMGVR_UViG_3300002927_000066_no_taxonomy_3300002927_LBH21_100025136
IMGVR_UViG_3300042064_000116_r__Varidnaviria_k__Bamfordvirae_p__Preplasmiviricota_c__Polintoviricetes_o__Orthopolintovirales____3300042064_Ga0451392_028222_3454_3969
GFS90966_1_Nephila pilipes
IMGVR_UViG_3300022562_002058_no_taxonomy_3300022562_Ga0247387_100428435
XP_009044799_1_Lottia gigantea
CAF1124636_1_Brachionus calyciflorus
IMGVR_UViG_3300042090_002534_r__Varidnaviria_k__Bamfordvirae_p__Preplasmiviricota_c__Polintoviricetes_o__Orthopolintovirales____3300042090_Ga0455669_00101224_381_836
IMGVR_UViG_3300003352_000010_r__Varidnaviria_k__Bamfordvirae_p__Preplasmiviricota_c__Polintoviricetes_o__Orthopolintovirales____3300003352_JGI26345J50200_10000514
IMGVR_UViG_3300027829_000066_r__Varidnaviria_k__Bamfordvirae_p__Preplasmiviricota_c__Polintoviricetes_o__Orthopolintovirales____3300027829_Ga0209773_100071583
XP_038069823_1_Patiria miniata
PIK43826_1_Apostichopus japonicus
RLJ22854_1_bacterium endosymbiont of Escarpia laminata
HBK84198_1_Flavobacterium sp.
KAG8236362_1_Ladona fulva
KAH7684611_1_Aphelenchus avenae
IMGVR_UViG_3300017912_004748_no_taxonomy_3300017912_Ga0192364_10720494
IMGVR_UViG_3300042062_000023_r__Varidnaviria_k__Bamfordvirae_p__Preplasmiviricota_c__Polintoviricetes_o__Orthopolintovirales____3300042062_Ga0451379_001914_1525_1980
IMGVR_UViG_3300042080_000096_r__Varidnaviria_k__Bamfordvirae_p__Preplasmiviricota_c__Polintoviricetes_o__Orthopolintovirales____3300042080_Ga0451378_001326_5819_6340
IMGVR_UViG_3300042063_000008_r__Varidnaviria_k__Bamfordvirae_p__Preplasmiviricota_c__Polintoviricetes_o__Orthopolintovirales____3300042063_Ga0451389_003758_418_936
IMGVR_UViG_3300042063_000009_r__Varidnaviria_k__Bamfordvirae_p__Preplasmiviricota_c__Polintoviricetes_o__Orthopolintovirales____3300042063_Ga0451389_003820_1421_1939
IMGVR_UViG_3300042064_000079_r__Varidnaviria_k__Bamfordvirae_p__Preplasmiviricota_c__Polintoviricetes_o__Orthopolintovirales____3300042064_Ga0451392_005926_7298_7855
IMGVR_UViG_3300042064_000054_r__Varidnaviria_k__Bamfordvirae_p__Preplasmiviricota_c__Polintoviricetes_o__Orthopolintovirales____3300042064_Ga0451392_006130_4041_4559
IMGVR_UViG_3300042080_000078_r__Varidnaviria_k__Bamfordvirae_p__Preplasmiviricota_c__Polintoviricetes_o__Orthopolintovirales____3300042080_Ga0451378_003048_444_965
IMGVR_UViG_3300042080_000027_r__Varidnaviria_k__Bamfordvirae_p__Preplasmiviricota_c__Polintoviricetes_o__Orthopolintovirales____3300042080_Ga0451378_010897_5812_6324
IMGVR_UViG_3300035317_000005_r__Varidnaviria_k__Bamfordvirae_p__Preplasmiviricota_c__Polintoviricetes_o__Orthopolintovirales____3300035317_Ga0377078_0006_4869_5384
IMGVR_UViG_3300035319_000002_r__Varidnaviria_k__Bamfordvirae_p__Preplasmiviricota_c__Polintoviricetes_o__Orthopolintovirales____3300035319_Ga0377080_0176_6050_6568
DAC80298_1_Betadintovirus terrapene
IMGVR_UViG_3300006001_000038_no_taxonomy_3300006001_Ga0081606_100132910
IMGVR_UViG_3300006001_000042_no_taxonomy_3300006001_Ga0081606_10030962
MCG8432192_1_Candidatus Omnitrophica bacterium
IMGVR_UViG_3300008034_000095_r__Varidnaviria_k__Bamfordvirae_p__Preplasmiviricota_c__Polintoviricetes_o__Orthopolintovirales____3300008034_Ga0099810_11769505
MAG_viral_043-00000001_3
DAC81414_1_Rhodactis coral adintovirus
XP_044172174_1_Acropora millepora
IMGVR_UViG_3300017907_000107_no_taxonomy_3300017907_Ga0192372_10269028
IMGVR_UViG_3300022217_000280_r__Varidnaviria_k__Bamfordvirae_p__Preplasmiviricota_c__Polintoviricetes_o__Orthopolintovirales____3300022217_Ga0224514_100034892
IMGVR_UViG_3300013759_000031_no_taxonomy_3300013759_Ga0119972_10020896
IMGVR_UViG_3300000133_000095_no_taxonomy_3300000133_SA_S1_NOR02_45mDRAFT_100020010
IMGVR_UViG_3300000130_000205_no_taxonomy_3300000130_SA_S2_NOR15_50mDRAFT_100024083
IMGVR_UViG_3300000128_000545_no_taxonomy_3300000128_SA_S1_NOR08_45mDRAFT_100024822
2.5

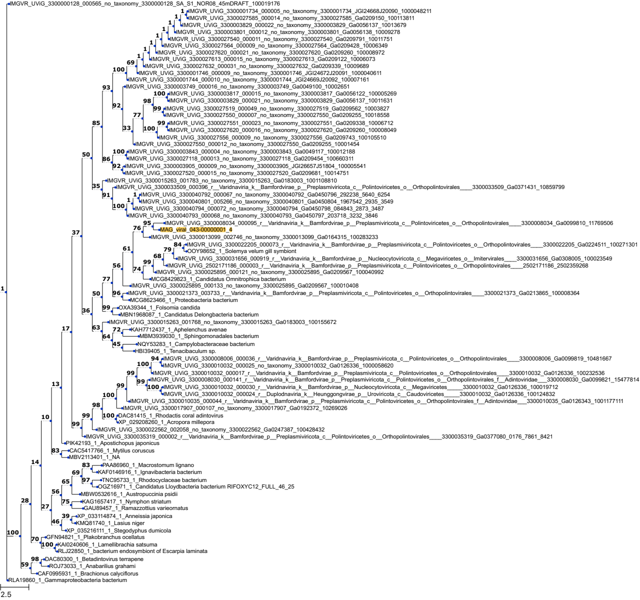
IMGVR_UViG_3300000128_000565_no_taxonomy_3300000128_SA_S1_NOR08_45mDRAFT_100019176
IMGVR_UViG_3300001734_000005_no_taxonomy_3300001734_JGI24668J20090_1000048211
IMGVR_UViG_3300027585_000014_no_taxonomy_3300027585_Ga0209150_100113811
IMGVR_UViG_3300003829_000022_no_taxonomy_3300003829_Ga0056137_10013679
IMGVR_UViG_3300003801_000012_no_taxonomy_3300003801_Ga0056138_10009278
IMGVR_UViG_3300027540_000011_no_taxonomy_3300027540_Ga0209791_10011751
IMGVR_UViG_3300027564_000009_no_taxonomy_3300027564_Ga0209428_10006349
IMGVR_UViG_3300027620_000021_no_taxonomy_3300027620_Ga0209260_100008972
IMGVR_UViG_3300027613_000015_no_taxonomy_3300027613_Ga0209122_10006073
IMGVR_UViG_3300027632_000031_no_taxonomy_3300027632_Ga0209339_10009689
IMGVR_UViG_3300001746_000009_no_taxonomy_3300001746_JGI24672J20091_1000040611
IMGVR_UViG_3300001744_000010_no_taxonomy_3300001744_JGI24669J20092_100007161
IMGVR_UViG_3300003749_000016_no_taxonomy_3300003749_Ga0049100_10002651
IMGVR_UViG_3300003817_000015_no_taxonomy_3300003817_Ga0056122_100005269
IMGVR_UViG_3300003829_000021_no_taxonomy_3300003829_Ga0056137_10011631
IMGVR_UViG_3300027519_000049_no_taxonomy_3300027519_Ga0209562_10003827
IMGVR_UViG_3300027550_000007_no_taxonomy_3300027550_Ga0209255_10018558
IMGVR_UViG_3300027551_000023_no_taxonomy_3300027551_Ga0209338_10006712
IMGVR_UViG_3300027620_000016_no_taxonomy_3300027620_Ga0209260_100008049
IMGVR_UViG_3300027556_000009_no_taxonomy_3300027556_Ga0209743_100105510
IMGVR_UViG_3300027550_000012_no_taxonomy_3300027550_Ga0209255_10001454
IMGVR_UViG_3300003843_000004_no_taxonomy_3300003843_Ga0049117_100012188
IMGVR_UViG_3300027118_000013_no_taxonomy_3300027118_Ga0209454_100660311
IMGVR_UViG_3300003905_000009_no_taxonomy_3300003905_JGI26657J51804_100005541
IMGVR_UViG_3300027520_000015_no_taxonomy_3300027520_Ga0209681_10014751
IMGVR_UViG_3300015263_001783_no_taxonomy_3300015263_Ga0183003_1001108810
IMGVR_UViG_3300033509_000396_r__Varidnaviria_k__Bamfordvirae_p__Preplasmiviricota_c__Polintoviricetes_o__Orthopolintovirales____3300033509_Ga0371431_10859799
IMGVR_UViG_3300040792_000067_no_taxonomy_3300040792_Ga0450796_292238_5640_6254
IMGVR_UViG_3300040801_005266_no_taxonomy_3300040801_Ga0450804_1967542_2935_3549
IMGVR_UViG_3300040794_000072_no_taxonomy_3300040794_Ga0450798_084843_2873_3487
IMGVR_UViG_3300040793_000068_no_taxonomy_3300040793_Ga0450797_203718_3232_3846
IMGVR_UViG_3300008034_000095_r__Varidnaviria_k__Bamfordvirae_p__Preplasmiviricota_c__Polintoviricetes_o__Orthopolintovirales____3300008034_Ga0099810_11769506
MAG_viral_043-00000001_4
IMGVR_UViG_3300013099_002746_no_taxonomy_3300013099_Ga0164315_100283233
IMGVR_UViG_3300022205_000073_r__Varidnaviria_k__Bamfordvirae_p__Preplasmiviricota_c__Polintoviricetes_o__Orthopolintovirales____3300022205_Ga0224511_100271301
OOY98652_1_Solemya velum gill symbiont
IMGVR_UViG_3300031656_000919_r__Varidnaviria_k__Bamfordvirae_p__Nucleocytoviricota_c__Megaviricetes_o__Imitervirales____3300031656_Ga0308005_100023549
IMGVR_UViG_2502171186_000003_r__Varidnaviria_k__Bamfordvirae_p__Preplasmiviricota_c__Polintoviricetes_o__Orthopolintovirales____2502171186_2502359268
IMGVR_UViG_3300025895_000121_no_taxonomy_3300025895_Ga0209567_100040992
MCG8429823_1_Candidatus Omnitrophica bacterium
IMGVR_UViG_3300025895_000133_no_taxonomy_3300025895_Ga0209567_100010408
IMGVR_UViG_3300021373_003733_r__Varidnaviria_k__Bamfordvirae_p__Preplasmiviricota_c__Polintoviricetes_o__Orthopolintovirales____3300021373_Ga0213865_100008364
MCG8623466_1_Proteobacteria bacterium
OXA39344_1_Folsomia candida
MBN1968087_1_Candidatus Delongbacteria bacterium
IMGVR_UViG_3300015263_001768_no_taxonomy_3300015263_Ga0183003_100155672
KAH7712437_1_Aphelenchus avenae
MBM3939030_1_Sphingomonadales bacterium
NQY53283_1_Campylobacteraceae bacterium
HBI39405_1_Tenacibaculum sp.
IMGVR_UViG_3300008006_000036_r__Varidnaviria_k__Bamfordvirae_p__Preplasmiviricota_c__Polintoviricetes_o__Orthopolintovirales____3300008006_Ga0099819_10481667
IMGVR_UViG_3300010032_000025_no_taxonomy_3300010032_Ga0126336_1000058620
IMGVR_UViG_3300010032_000017_r__Varidnaviria_k__Bamfordvirae_p__Preplasmiviricota_c__Polintoviricetes_o__Orthopolintovirales____3300010032_Ga0126336_100232536
IMGVR_UViG_3300008030_000141_r__Varidnaviria_k__Bamfordvirae_p__Preplasmiviricota_c__Polintoviricetes_o__Orthopolintovirales_f__Adintoviridae____3300008030_Ga0099821_15477814
IMGVR_UViG_3300010032_000030_r__Varidnaviria_k__Bamfordvirae_p__Nucleocytoviricota_c__Megaviricetes____3300010032_Ga0126336_100019712
IMGVR_UViG_3300010032_000024_r__Duplodnaviria_k__Heunggongvirae_p__Uroviricota_c__Caudoviricetes____3300010032_Ga0126336_100124832
IMGVR_UViG_3300010035_000044_r__Varidnaviria_k__Bamfordvirae_p__Preplasmiviricota_c__Polintoviricetes_o__Orthopolintovirales_f__Adintoviridae____3300010035_Ga0126343_1001177111
IMGVR_UViG_3300017907_000107_no_taxonomy_3300017907_Ga0192372_10269026
DAC81415_1_Rhodactis coral adintovirus
XP_029208260_1_Acropora millepora
IMGVR_UViG_3300022562_002058_no_taxonomy_3300022562_Ga0247387_100428432
IMGVR_UViG_3300035319_000002_r__Varidnaviria_k__Bamfordvirae_p__Preplasmiviricota_c__Polintoviricetes_o__Orthopolintovirales____3300035319_Ga0377080_0176_7861_8421
PIK42193_1_Apostichopus japonicus
CAC5417766_1_Mytilus coruscus
MBV2113401_1_NA
PAA86960_1_Macrostomum lignano
KAF0146916_1_Ignavibacteria bacterium
TNC95733_1_Rhodocyclaceae bacterium
OGZ16971_1_Candidatus Lloydbacteria bacterium RIFOXYC12_FULL_46_25
MBW0532616_1_Austropuccinia psidii
KAG1657417_1_Nymphon striatum
GAU89457_1_Ramazzottius varieornatus
XP_033114874_1_Anneissia japonica
KMQ81740_1_Lasius niger
XP_035216111_1_Stegodyphus dumicola
GFN94821_1_Plakobranchus ocellatus
KAI0240606_1_Lamellibrachia satsuma
RLJ22850_1_bacterium endosymbiont of Escarpia laminata
DAC80300_1_Betadintovirus terrapene
ROJ73033_1_Anabarilius grahami
CAF0995931_1_Brachionus calyciflorus
RLA19860_1_Gammaproteobacteria bacterium
2.5

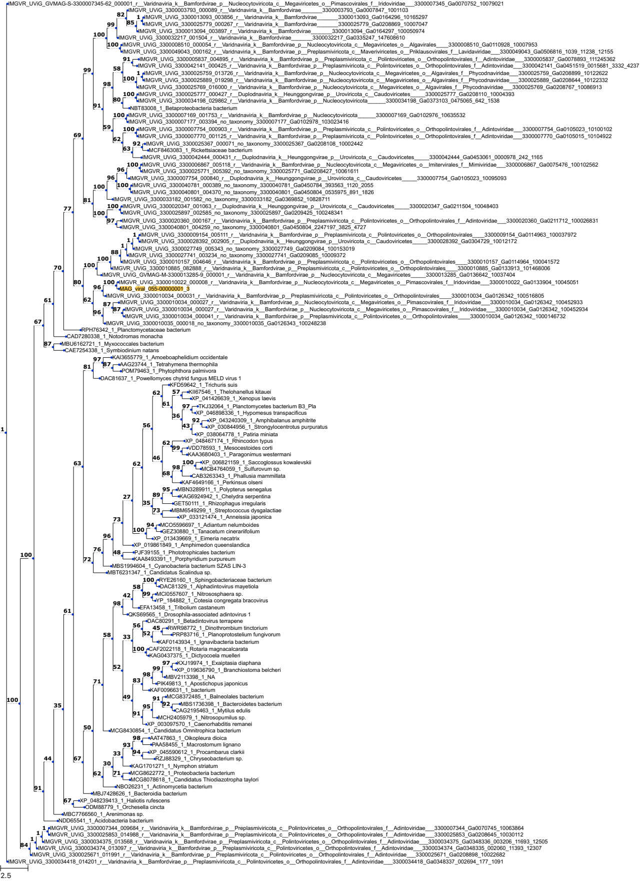
IMGVR_UViG_GVMAG-S-3300007345-62_000001_r__Varidnaviria_k__Bamfordvirae_p__Nucleocytoviricota_c__Megaviricetes_o__Pimascovirales_f__Iridoviridae___3300007345_Ga0070752_10079021
IMGVR_UViG_3300003793_000089_r__Varidnaviria_k__Bamfordvirae_______3300003793_Ga0007847_1001103
IMGVR_UViG_3300013093_003856_r__Varidnaviria_k__Bamfordvirae_______3300013093_Ga0164296_10165297
IMGVR_UViG_3300025779_000267_r__Varidnaviria_k__Bamfordvirae_______3300025779_Ga0208869_10007047
IMGVR_UViG_3300013094_003897_r__Varidnaviria_k__Bamfordvirae_______3300013094_Ga0164297_100050974
IMGVR_UViG_3300032217_001504_r__Varidnaviria_k__Bamfordvirae_______3300032217_Ga0335247_147608610
IMGVR_UViG_3300008510_000054_r__Varidnaviria_k__Bamfordvirae_p__Nucleocytoviricota_c__Megaviricetes_o__Algavirales_____3300008510_Ga0110928_10007953
IMGVR_UViG_3300049043_000162_r__Varidnaviria_k__Bamfordvirae_p__Preplasmiviricota_c__Maveriviricetes_o__Priklausovirales_f__Lavidaviridae___3300049043_Ga0506816_1039_11238_12155
IMGVR_UViG_3300005837_004895_r__Varidnaviria_k__Bamfordvirae_p__Preplasmiviricota_c__Polintoviricetes_o__Orthopolintovirales_f__Adintoviridae___3300005837_Ga0078893_111245362
IMGVR_UViG_3300042141_000425_r__Varidnaviria_k__Bamfordvirae_p__Preplasmiviricota_c__Polintoviricetes_o__Orthopolintovirales_f__Adintoviridae___3300042141_Ga0451519_0015681_3332_4237
IMGVR_UViG_3300025759_013726_r__Varidnaviria_k__Bamfordvirae_p__Nucleocytoviricota_c__Megaviricetes_o__Algavirales_f__Phycodnaviridae___3300025759_Ga0208899_10122622
IMGVR_UViG_3300025889_019298_r__Varidnaviria_k__Bamfordvirae_p__Nucleocytoviricota_c__Megaviricetes_o__Algavirales_f__Phycodnaviridae___3300025889_Ga0208644_10122332
IMGVR_UViG_3300025769_016000_r__Varidnaviria_k__Bamfordvirae_p__Nucleocytoviricota_c__Megaviricetes_o__Algavirales_f__Phycodnaviridae___3300025769_Ga0208767_10086913
IMGVR_UViG_3300025777_000427_r__Duplodnaviria_k__Heunggongvirae_p__Uroviricota_c__Caudoviricetes_____3300025777_Ga0208110_10004393
IMGVR_UViG_3300034198_029862_r__Varidnaviria_k__Bamfordvirae_p__Nucleocytoviricota_______3300034198_Ga0373103_0475065_642_1538
NBT83008_1_Betaproteobacteria bacterium
IMGVR_UViG_3300007169_001753_r__Varidnaviria_k__Bamfordvirae_p__Nucleocytoviricota______3300007169_Ga0102976_10635532
IMGVR_UViG_3300007177_003394_no_taxonomy_3300007177_Ga0102978_103023416
IMGVR_UViG_3300007754_000903_r__Varidnaviria_k__Bamfordvirae_p__Preplasmiviricota_c__Polintoviricetes_o__Orthopolintovirales_f__Adintoviridae___3300007754_Ga0105023_10100102
IMGVR_UViG_3300007770_001125_r__Varidnaviria_k__Bamfordvirae_p__Preplasmiviricota_c__Polintoviricetes_o__Orthopolintovirales_f__Adintoviridae___3300007770_Ga0105015_10104922
IMGVR_UViG_3300025367_000071_no_taxonomy_3300025367_Ga0208108_10002442
MCF8463083_1_Rickettsiaceae bacterium
IMGVR_UViG_3300042444_000431_r__Duplodnaviria_k__Heunggongvirae_p__Uroviricota_c__Caudoviricetes_____3300042444_Ga0453061_0000978_242_1165
IMGVR_UViG_3300006867_005118_r__Varidnaviria_k__Bamfordvirae_p__Nucleocytoviricota_c__Megaviricetes_o__Imitervirales_f__Mimiviridae___3300006867_Ga0075476_100102562
IMGVR_UViG_3300025771_005392_no_taxonomy_3300025771_Ga0208427_10061611
IMGVR_UViG_3300007754_000840_r__Duplodnaviria_k__Heunggongvirae_p__Uroviricota_c__Caudoviricetes_____3300007754_Ga0105023_10095093
IMGVR_UViG_3300040781_000389_no_taxonomy_3300040781_Ga0450784_393563_1120_2055
IMGVR_UViG_3300040801_004370_no_taxonomy_3300040801_Ga0450804_0535975_891_1826
IMGVR_UViG_3300033182_001582_no_taxonomy_3300033182_Ga0369852_10828711
IMGVR_UViG_3300020347_001063_r__Duplodnaviria_k__Heunggongvirae_p__Uroviricota_c__Caudoviricetes_____3300020347_Ga0211504_10048403
IMGVR_UViG_3300025897_002585_no_taxonomy_3300025897_Ga0209425_100248341
IMGVR_UViG_3300020360_000167_r__Varidnaviria_k__Bamfordvirae_p__Preplasmiviricota_c__Polintoviricetes_o__Orthopolintovirales_f__Adintoviridae___3300020360_Ga0211712_100026831
IMGVR_UViG_3300040801_004259_no_taxonomy_3300040801_Ga0450804_2247197_3825_4727
IMGVR_UViG_3300009154_005111_r__Varidnaviria_k__Bamfordvirae_p__Preplasmiviricota_c__Polintoviricetes_o__Orthopolintovirales_____3300009154_Ga0114963_100037972
IMGVR_UViG_3300028392_002905_r__Duplodnaviria_k__Heunggongvirae_p__Uroviricota_c__Caudoviricetes_____3300028392_Ga0304729_10012172
IMGVR_UViG_3300027749_005343_no_taxonomy_3300027749_Ga0209084_100153019
IMGVR_UViG_3300027741_003234_no_taxonomy_3300027741_Ga0209085_10009372
IMGVR_UViG_3300010157_004646_r__Varidnaviria_k__Bamfordvirae_p__Preplasmiviricota_c__Polintoviricetes_o__Orthopolintovirales_____3300010157_Ga0114964_100041572
IMGVR_UViG_3300010885_082888_r__Varidnaviria_k__Bamfordvirae_p__Preplasmiviricota_c__Polintoviricetes_o__Orthopolintovirales_____3300010885_Ga0133913_101468006
IMGVR_UViG_GVMAG-M-3300013285-9_000001_r__Varidnaviria_k__Bamfordvirae_p__Nucleocytoviricota_c__Megaviricetes_____3300013285_Ga0136642_10037404
IMGVR_UViG_3300010022_000008_r__Varidnaviria_k__Bamfordvirae_p__Nucleocytoviricota_c__Megaviricetes_o__Pimascovirales_f__Iridoviridae___3300010022_Ga0133904_10045051
MAG_viral_055-00000001_3
IMGVR_UViG_3300010034_000031_r__Varidnaviria_k__Bamfordvirae_p__Preplasmiviricota_c__Polintoviricetes_o__Orthopolintovirales_____3300010034_Ga0126342_100516805
IMGVR_UViG_3300010034_000027_r__Varidnaviria_k__Bamfordvirae_p__Nucleocytoviricota_c__Megaviricetes_o__Pimascovirales_f__Iridoviridae___3300010034_Ga0126342_100452933
IMGVR_UViG_3300010034_000027_r__Varidnaviria_k__Bamfordvirae_p__Nucleocytoviricota_c__Megaviricetes_o__Pimascovirales_f__Iridoviridae___3300010034_Ga0126342_100452934
IMGVR_UViG_3300010034_000041_r__Varidnaviria_k__Bamfordvirae_p__Preplasmiviricota_c__Polintoviricetes_o__Orthopolintovirales_____3300010034_Ga0126342_1000146732
IMGVR_UViG_3300010035_000018_no_taxonomy_3300010035_Ga0126343_100248238
RPH76342_1_Planctomycetaceae bacterium
CAD7280338_1_Notodromas monacha
MBU6162721_1_Myxococcales bacterium
CAE7254338_1_Symbiodinium natans
KAI3655779_1_Amoeboaphelidium occidentale
AAG23744_1_Tetrahymena thermophila
POM79463_1_Phytophthora palmivora
DAC81637_1_Powellomyces chytrid fungus MELD virus 1
KFD59642_1_Trichuris suis
KII67546_1_Thelohanellus kitauei
XP_041426639_1_Xenopus laevis
TKJ32064_1_Planctomycetes bacterium B3_Pla
XP_046898336_1_Hypomesus transpacificus
XP_043240309_1_Amphibalanus amphitrite
XP_030844956_1_Strongylocentrotus purpuratus
XP_038064778_1_Patiria miniata
XP_048467174_1_Rhincodon typus
VDD78593_1_Mesocestoides corti
KAA3680403_1_Paragonimus westermani
XP_006821159_1_Saccoglossus kowalevskii
MCB4764059_1_Sulfurovum sp.
CAB3263343_1_Phallusia mammillata
KAF4649166_1_Perkinsus olseni
MBN3289911_1_Polypterus senegalus
KAG6924942_1_Chelydra serpentina
GET50111_1_Rhizophagus irregularis
MBM6549299_1_Streptococcus dysgalactiae
XP_033121474_1_Anneissia japonica
MCO5596697_1_Adiantum nelumboides
GEZ30880_1_Tanacetum cinerariifolium
XP_013439669_1_Eimeria necatrix
XP_019861849_1_Amphimedon queenslandica
PJF39155_1_Phototrophicales bacterium
KAA8493391_1_Porphyridium purpureum
MBS1994604_1_Cyanobacteria bacterium SZAS LIN-3
MBT6231347_1_Candidatus Scalindua sp.
RYE26160_1_Sphingobacteriaceae bacterium
DAC81329_1_Alphadintovirus mayetiola
MCI0557607_1_Nitrososphaera sp.
YP_184882_1_Cotesia congregata bracovirus
EFA13458_1_Tribolium castaneum
QKS69565_1_Drosophila-associated adintovirus 1
DAC80291_1_Betadintovirus terrapene
RWR98772_1_Dinothrombium tinctorium
PRP83716_1_Planoprotostelium fungivorum
KAF0143934_1_Ignavibacteria bacterium
CAF2022118_1_Rotaria magnacalcarata
KAG0437375_1_Dictyocoela muelleri
KXJ19974_1_Exaiptasia diaphana
XP_019636790_1_Branchiostoma belcheri
MBV2113398_1_NA
PIK49813_1_Apostichopus japonicus
KAF0096631_1_bacterium
MCG8372485_1_Balneolales bacterium
MBS1736398_1_Bacteroidetes bacterium
CAG2195463_1_Mytilus edulis
MCH2405979_1_Nitrosopumilus sp.
XP_003097570_1_Caenorhabditis remanei
MCG8430854_1_Candidatus Omnitrophica bacterium
AAT47863_1_Oikopleura dioica
PAA58455_1_Macrostomum lignano
XP_045590612_1_Procambarus clarkii
RZJ88329_1_Chryseobacterium sp.
KAG1701271_1_Nymphon striatum
MCG8622772_1_Proteobacteria bacterium
MCG8078618_1_Candidatus Thiodiazotropha taylori
NBO26231_1_Actinomycetia bacterium
MBJ7428626_1_Bacteroidia bacterium
XP_048239413_1_Haliotis rufescens
ODM88779_1_Orchesella cincta
MBC7766560_1_Arenimonas sp.
NDD65541_1_Acidobacteria bacterium
IMGVR_UViG_3300007344_009684_r__Varidnaviria_k__Bamfordvirae_p__Preplasmiviricota_c__Polintoviricetes_o__Orthopolintovirales_f__Adintoviridae___3300007344_Ga0070745_10063864
IMGVR_UViG_3300025853_014988_r__Varidnaviria_k__Bamfordvirae_p__Preplasmiviricota_c__Polintoviricetes_o__Orthopolintovirales_f__Adintoviridae___3300025853_Ga0208645_10030112
IMGVR_UViG_3300034375_013568_r__Varidnaviria_k__Bamfordvirae_p__Preplasmiviricota_c__Polintoviricetes_o__Orthopolintovirales_f__Adintoviridae___3300034375_Ga0348336_003206_11693_12505
IMGVR_UViG_3300034374_013097_r__Varidnaviria_k__Bamfordvirae_p__Preplasmiviricota_c__Polintoviricetes_o__Orthopolintovirales_f__Adintoviridae___3300034374_Ga0348335_002060_11393_12307
IMGVR_UViG_3300025671_011991_r__Varidnaviria_k__Bamfordvirae_p__Preplasmiviricota_c__Polintoviricetes_o__Orthopolintovirales_f__Adintoviridae___3300025671_Ga0208898_10022682
IMGVR_UViG_3300034418_014201_r__Varidnaviria_k__Bamfordvirae_p__Preplasmiviricota_c__Polintoviricetes_o__Orthopolintovirales_f__Adintoviridae___3300034418_Ga0348337_002694_177_1091
2.5

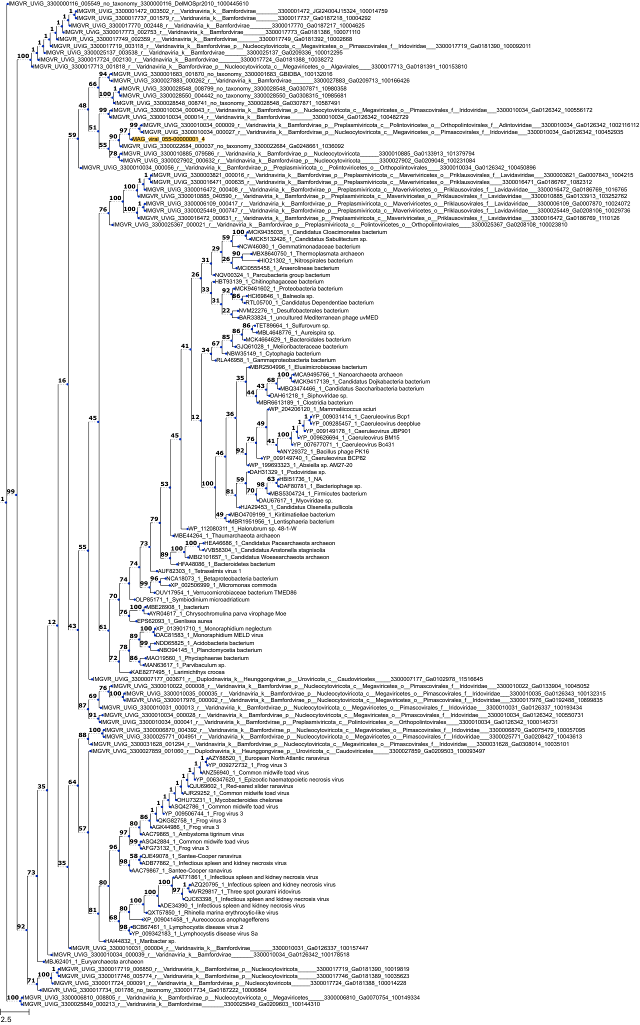
IMGVR_UViG_3300000116_005549_no_taxonomy_3300000116_DelMOSpr2010_1000445610
IMGVR_UViG_3300001472_003502_r__Varidnaviria_k__Bamfordvirae________3300001472_JGI24004J15324_100014759
IMGVR_UViG_3300017737_001579_r__Varidnaviria_k__Bamfordvirae________3300017737_Ga0187218_10004292
IMGVR_UViG_3300017770_002448_r__Varidnaviria_k__Bamfordvirae________3300017770_Ga0187217_10004625
IMGVR_UViG_3300017773_002753_r__Varidnaviria_k__Bamfordvirae________3300017773_Ga0181386_100071110
IMGVR_UViG_3300017749_002359_r__Varidnaviria_k__Bamfordvirae________3300017749_Ga0181392_10002668
IMGVR_UViG_3300017719_003118_r__Varidnaviria_k__Bamfordvirae_p__Nucleocytoviricota_c__Megaviricetes_o__Pimascovirales_f__Iridoviridae___3300017719_Ga0181390_100092011
IMGVR_UViG_3300025137_003538_r__Varidnaviria_k__Bamfordvirae________3300025137_Ga0209336_100012295
IMGVR_UViG_3300017724_002130_r__Varidnaviria_k__Bamfordvirae________3300017724_Ga0181388_10038272
IMGVR_UViG_3300017713_001818_r__Varidnaviria_k__Bamfordvirae_p__Nucleocytoviricota_c__Megaviricetes_o__Algavirales____3300017713_Ga0181391_100153810
IMGVR_UViG_3300001683_001870_no_taxonomy_3300001683_GBIDBA_100132016
IMGVR_UViG_3300027883_000262_r__Varidnaviria_k__Bamfordvirae________3300027883_Ga0209713_100166426
IMGVR_UViG_3300028548_008799_no_taxonomy_3300028548_Ga0307871_10980358
IMGVR_UViG_3300028550_004442_no_taxonomy_3300028550_Ga0308315_10985681
IMGVR_UViG_3300028548_008741_no_taxonomy_3300028548_Ga0307871_10587491
IMGVR_UViG_3300010034_000043_r__Varidnaviria_k__Bamfordvirae_p__Nucleocytoviricota_c__Megaviricetes_o__Pimascovirales_f__Iridoviridae___3300010034_Ga0126342_100556172
IMGVR_UViG_3300010034_000014_r__Varidnaviria_k__Bamfordvirae________3300010034_Ga0126342_100482729
IMGVR_UViG_3300010034_000009_r__Varidnaviria_k__Bamfordvirae_p__Preplasmiviricota_c__Polintoviricetes_o__Orthopolintovirales_f__Adintoviridae___3300010034_Ga0126342_1002116112
IMGVR_UViG_3300010034_000027_r__Varidnaviria_k__Bamfordvirae_p__Nucleocytoviricota_c__Megaviricetes_o__Pimascovirales_f__Iridoviridae___3300010034_Ga0126342_100452935
MAG_viral_055-00000001_4
IMGVR_UViG_3300022684_000037_no_taxonomy_3300022684_Ga0248661_1036092
IMGVR_UViG_3300010885_079586_r__Varidnaviria_k__Bamfordvirae_p__Nucleocytoviricota______3300010885_Ga0133913_101379794
IMGVR_UViG_3300027902_000632_r__Varidnaviria_k__Bamfordvirae_p__Nucleocytoviricota______3300027902_Ga0209048_100231084
IMGVR_UViG_3300010034_000056_r__Varidnaviria_k__Bamfordvirae_p__Preplasmiviricota_c__Polintoviricetes_o__Orthopolintovirales____3300010034_Ga0126342_100450896
IMGVR_UViG_3300003821_000016_r__Varidnaviria_k__Bamfordvirae_p__Preplasmiviricota_c__Maveriviricetes_o__Priklausovirales_f__Lavidaviridae___3300003821_Ga0007843_1004215
IMGVR_UViG_3300016471_000635_r__Varidnaviria_k__Bamfordvirae_p__Preplasmiviricota_c__Maveriviricetes_o__Priklausovirales____3300016471_Ga0186767_1082312
IMGVR_UViG_3300016472_000408_r__Varidnaviria_k__Bamfordvirae_p__Preplasmiviricota_c__Maveriviricetes_o__Priklausovirales_f__Lavidaviridae___3300016472_Ga0186769_1016765
IMGVR_UViG_3300010885_040590_r__Varidnaviria_k__Bamfordvirae_p__Preplasmiviricota_c__Maveriviricetes_o__Priklausovirales_f__Lavidaviridae___3300010885_Ga0133913_103252762
IMGVR_UViG_3300006109_000417_r__Varidnaviria_k__Bamfordvirae_p__Preplasmiviricota_c__Maveriviricetes_o__Priklausovirales_f__Lavidaviridae___3300006109_Ga0007870_10024072
IMGVR_UViG_3300025449_000747_r__Varidnaviria_k__Bamfordvirae_p__Preplasmiviricota_c__Maveriviricetes_o__Priklausovirales_f__Lavidaviridae___3300025449_Ga0208106_10029736
IMGVR_UViG_3300016472_000631_r__Varidnaviria_k__Bamfordvirae_p__Preplasmiviricota_c__Maveriviricetes_o__Priklausovirales_f__Lavidaviridae___3300016472_Ga0186769_1110126
IMGVR_UViG_3300025367_000021_r__Varidnaviria_k__Bamfordvirae_p__Preplasmiviricota_c__Polintoviricetes_o__Orthopolintovirales____3300025367_Ga0208108_100023810
MCK9435035_1_Candidatus Cloacimonetes bacterium
MCK5132426_1_Candidatus Sabulitectum sp.
NCW46080_1_Gemmatimonadaceae bacterium
MBX8640750_1_Thermoplasmata archaeon
HIO21302_1_Nitrospirales bacterium
MCI0555458_1_Anaerolineae bacterium
NQV00324_1_Parcubacteria group bacterium
HBT93139_1_Chitinophagaceae bacterium
MCK9461602_1_Proteobacteria bacterium
HCI69846_1_Balneola sp.
RTL05700_1_Candidatus Dependentiae bacterium
NVM22276_1_Desulfobacterales bacterium
BAR33824_1_uncultured Mediterranean phage uvMED
TET89664_1_Sulfurovum sp.
MBL4648776_1_Aureispira sp.
MCK4664629_1_Bacteroidales bacterium
GJQ61028_1_Melioribacteraceae bacterium
NBW35149_1_Cytophagia bacterium
RLA46958_1_Gammaproteobacteria bacterium
MBR2504996_1_Elusimicrobiaceae bacterium
MCA9495766_1_Nanoarchaeota archaeon
MCK9417139_1_Candidatus Dojkabacteria bacterium
MBQ3474466_1_Candidatus Saccharibacteria bacterium
DAH61218_1_Siphoviridae sp.
MBR6613189_1_Clostridia bacterium
WP_204206120_1_Mammaliicoccus sciuri
YP_009031414_1_Caeruleovirus Bcp1
YP_009285457_1_Caeruleovirus deepblue
YP_009149178_1_Caeruleovirus JBP901
YP_009626694_1_Caeruleovirus BM15
YP_007677071_1_Caeruleovirus Bc431
ANY29372_1_Bacillus phage PK16
YP_009149740_1_Caeruleovirus BCP82
WP_199693323_1_Absiella sp. AM27-20
DAH31329_1_Podoviridae sp.
HBI51736_1_NA
DAF80781_1_Bacteriophage sp.
MBS5304724_1_Firmicutes bacterium
DAU67617_1_Myoviridae sp.
HJA29453_1_Candidatus Olsenella pullicola
MBO4709199_1_Kiritimatiellae bacterium
MBR1951956_1_Lentisphaeria bacterium
WP_112080311_1_Halorubrum sp. 48-1-W
MBE44264_1_Thaumarchaeota archaeon
HEA46686_1_Candidatus Pacearchaeota archaeon
VVB58304_1_Candidatus Anstonella stagnisolia
MBI2101657_1_Candidatus Woesearchaeota archaeon
HFA48086_1_Bacteroidetes bacterium
AUF82303_1_Tetraselmis virus 1
NCA18073_1_Betaproteobacteria bacterium
XP_002506999_1_Micromonas commoda
OUV17954_1_Verrucomicrobiaceae bacterium TMED86
OLP85171_1_Symbiodinium microadriaticum
MBE28908_1_bacterium
AYR04617_1_Chrysochromulina parva virophage Moe
EPS62093_1_Genlisea aurea
XP_013901710_1_Monoraphidium neglectum
DAC81583_1_Monoraphidium MELD virus
NDD65825_1_Acidobacteria bacterium
NBO94145_1_Planctomycetia bacterium
MAO19560_1_Phycisphaerae bacterium
MAN63617_1_Parvibaculum sp.
KAE8277495_1_Larimichthys crocea
IMGVR_UViG_3300007177_003671_r__Duplodnaviria_k__Heunggongvirae_p__Uroviricota_c__Caudoviricetes_____3300007177_Ga0102978_11516645
IMGVR_UViG_3300010022_000008_r__Varidnaviria_k__Bamfordvirae_p__Nucleocytoviricota_c__Megaviricetes_o__Pimascovirales_f__Iridoviridae___3300010022_Ga0133904_10045052
IMGVR_UViG_3300010035_000035_r__Varidnaviria_k__Bamfordvirae_p__Nucleocytoviricota_c__Megaviricetes_o__Pimascovirales_f__Iridoviridae___3300010035_Ga0126343_100132315
IMGVR_UViG_3300017976_000002_r__Varidnaviria_k__Bamfordvirae_p__Nucleocytoviricota_c__Megaviricetes_o__Pimascovirales_f__Iridoviridae___3300017976_Ga0192488_10899835
IMGVR_UViG_3300010031_000013_r__Varidnaviria_k__Bamfordvirae_p__Nucleocytoviricota_c__Megaviricetes_o__Pimascovirales_f__Iridoviridae___3300010031_Ga0126337_100193434
IMGVR_UViG_3300010034_000028_r__Varidnaviria_k__Bamfordvirae_p__Nucleocytoviricota_c__Megaviricetes_o__Pimascovirales_f__Iridoviridae___3300010034_Ga0126342_100550731
IMGVR_UViG_3300010034_000041_r__Varidnaviria_k__Bamfordvirae_p__Preplasmiviricota_c__Polintoviricetes_o__Orthopolintovirales____3300010034_Ga0126342_1000146731
IMGVR_UViG_3300006870_004392_r__Varidnaviria_k__Bamfordvirae_p__Nucleocytoviricota_c__Megaviricetes_o__Pimascovirales_f__Iridoviridae___3300006870_Ga0075479_100057095
IMGVR_UViG_3300025771_004951_r__Varidnaviria_k__Bamfordvirae_p__Nucleocytoviricota_c__Megaviricetes_o__Pimascovirales_f__Iridoviridae___3300025771_Ga0208427_10043613
IMGVR_UViG_3300031628_001294_r__Varidnaviria_k__Bamfordvirae_p__Nucleocytoviricota_c__Megaviricetes_o__Pimascovirales_f__Iridoviridae___3300031628_Ga0308014_10035101
IMGVR_UViG_3300027859_001060_r__Duplodnaviria_k__Heunggongvirae_p__Uroviricota_c__Caudoviricetes_____3300027859_Ga0209503_100093497
AZY88520_1_European North Atlantic ranavirus
YP_009272732_1_Frog virus 3
ANZ56940_1_Common midwife toad virus
YP_006347620_1_Epizootic haematopoietic necrosis virus
QJU69602_1_Red-eared slider ranavirus
AJR29252_1_Common midwife toad virus
OHU73231_1_Mycobacteroides chelonae
ASQ42786_1_Common midwife toad virus
YP_009506744_1_Frog virus 3
QKG82758_1_Frog virus 3
AGK44986_1_Frog virus 3
AAC79865_1_Ambystoma tigrinum virus
ASQ42884_1_Common midwife toad virus
AFG73132_1_Frog virus 3
QJE49078_1_Santee-Cooper ranavirus
ADB77862_1_Infectious spleen and kidney necrosis virus
AAC79867_1_Santee-Cooper ranavirus
AAT71861_1_Infectious spleen and kidney necrosis virus
AZQ20795_1_Infectious spleen and kidney necrosis virus
AVR29817_1_Three spot gourami iridovirus
QJC63398_1_Infectious spleen and kidney necrosis virus
ADE34390_1_Infectious spleen and kidney necrosis virus
QXT57850_1_Rhinella marina erythrocytic-like virus
XP_009041458_1_Aureococcus anophagefferens
BCB67461_1_Lymphocystis disease virus 2
YP_009342183_1_Lymphocystis disease virus Sa
HAI44832_1_Maribacter sp.
IMGVR_UViG_3300010031_000004_r__Varidnaviria_k__Bamfordvirae________3300010031_Ga0126337_100157447
IMGVR_UViG_3300010034_000039_r__Varidnaviria_k__Bamfordvirae________3300010034_Ga0126342_100178518
MBJ62401_1_Euryarchaeota archaeon
IMGVR_UViG_3300017719_006850_r__Varidnaviria_k__Bamfordvirae_p__Nucleocytoviricota______3300017719_Ga0181390_10019819
IMGVR_UViG_3300017746_005774_r__Varidnaviria_k__Bamfordvirae_p__Nucleocytoviricota______3300017746_Ga0181389_10035623
IMGVR_UViG_3300017724_000091_r__Varidnaviria_k__Bamfordvirae_p__Nucleocytoviricota______3300017724_Ga0181388_100014228
IMGVR_UViG_3300017734_001786_no_taxonomy_3300017734_Ga0187222_10006864
IMGVR_UViG_3300006810_008805_r__Varidnaviria_k__Bamfordvirae_p__Nucleocytoviricota_c__Megaviricetes_____3300006810_Ga0070754_100149334
IMGVR_UViG_3300025849_000213_r__Varidnaviria_k__Bamfordvirae________3300025849_Ga0209603_100144310
2.5

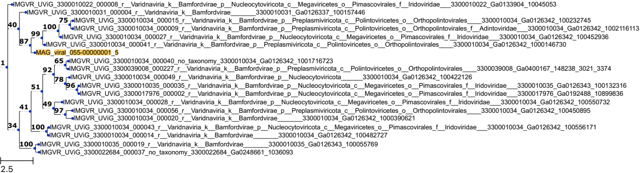
IMGVR_UViG_3300010022_000008_r__Varidnaviria_k__Bamfordvirae_p__Nucleocytoviricota_c__Megaviricetes_o__Pimascovirales_f__Iridoviridae___3300010022_Ga0133904_10045053
IMGVR_UViG_3300010031_000004_r__Varidnaviria_k__Bamfordvirae_______3300010031_Ga0126337_100157446
IMGVR_UViG_3300010034_000015_r__Varidnaviria_k__Bamfordvirae_p__Preplasmiviricota_c__Polintoviricetes_o__Orthopolintovirales_____3300010034_Ga0126342_100232745
IMGVR_UViG_3300010034_000009_r__Varidnaviria_k__Bamfordvirae_p__Preplasmiviricota_c__Polintoviricetes_o__Orthopolintovirales_f__Adintoviridae___3300010034_Ga0126342_1002116113
IMGVR_UViG_3300010034_000027_r__Varidnaviria_k__Bamfordvirae_p__Nucleocytoviricota_c__Megaviricetes_o__Pimascovirales_f__Iridoviridae___3300010034_Ga0126342_100452936
IMGVR_UViG_3300010034_000041_r__Varidnaviria_k__Bamfordvirae_p__Preplasmiviricota_c__Polintoviricetes_o__Orthopolintovirales_____3300010034_Ga0126342_1000146730
MAG_viral_055-00000001_5
IMGVR_UViG_3300010034_000040_no_taxonomy_3300010034_Ga0126342_1001716723
IMGVR_UViG_3300039008_000227_r__Varidnaviria_k__Bamfordvirae_p__Preplasmiviricota_c__Polintoviricetes_o__Orthopolintovirales_____3300039008_Ga0400167_148238_3021_3374
IMGVR_UViG_3300010034_000049_r__Varidnaviria_k__Bamfordvirae_p__Nucleocytoviricota_______3300010034_Ga0126342_100422126
IMGVR_UViG_3300010035_000035_r__Varidnaviria_k__Bamfordvirae_p__Nucleocytoviricota_c__Megaviricetes_o__Pimascovirales_f__Iridoviridae___3300010035_Ga0126343_100132316
IMGVR_UViG_3300017976_000002_r__Varidnaviria_k__Bamfordvirae_p__Nucleocytoviricota_c__Megaviricetes_o__Pimascovirales_f__Iridoviridae___3300017976_Ga0192488_10899836
IMGVR_UViG_3300010034_000028_r__Varidnaviria_k__Bamfordvirae_p__Nucleocytoviricota_c__Megaviricetes_o__Pimascovirales_f__Iridoviridae___3300010034_Ga0126342_100550732
IMGVR_UViG_3300010034_000056_r__Varidnaviria_k__Bamfordvirae_p__Preplasmiviricota_c__Polintoviricetes_o__Orthopolintovirales_____3300010034_Ga0126342_100450895
IMGVR_UViG_3300010034_000020_r__Varidnaviria_k__Bamfordvirae_______3300010034_Ga0126342_1000390621
IMGVR_UViG_3300010034_000043_r__Varidnaviria_k__Bamfordvirae_p__Nucleocytoviricota_c__Megaviricetes_o__Pimascovirales_f__Iridoviridae___3300010034_Ga0126342_100556171
IMGVR_UViG_3300010034_000014_r__Varidnaviria_k__Bamfordvirae_______3300010034_Ga0126342_100482727
IMGVR_UViG_3300010035_000019_r__Varidnaviria_k__Bamfordvirae_______3300010035_Ga0126343_100055769
IMGVR_UViG_3300022684_000037_no_taxonomy_3300022684_Ga0248661_1036093
40
75
100
99
87
1
65
92
78
96
51
49
97
41
34
100
100
2.5

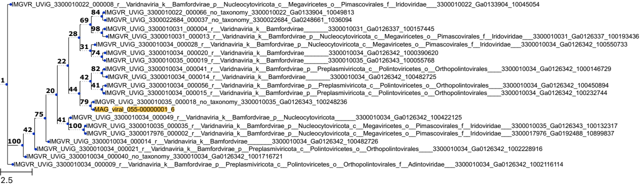
IMGVR_UViG_3300010022_000008_r__Varidnaviria_k__Bamfordvirae_p__Nucleocytoviricota_c__Megaviricetes_o__Pimascovirales_f__Iridoviridae___3300010022_Ga0133904_10045054
IMGVR_UViG_3300010022_000066_no_taxonomy_3300010022_Ga0133904_10049813
IMGVR_UViG_3300022684_000037_no_taxonomy_3300022684_Ga0248661_1036094
IMGVR_UViG_3300010031_000004_r__Varidnaviria_k__Bamfordvirae_______3300010031_Ga0126337_100157445
IMGVR_UViG_3300010031_000013_r__Varidnaviria_k__Bamfordvirae_p__Nucleocytoviricota_c__Megaviricetes_o__Pimascovirales_f__Iridoviridae___3300010031_Ga0126337_100193436
IMGVR_UViG_3300010034_000028_r__Varidnaviria_k__Bamfordvirae_p__Nucleocytoviricota_c__Megaviricetes_o__Pimascovirales_f__Iridoviridae___3300010034_Ga0126342_100550733
IMGVR_UViG_3300010034_000020_r__Varidnaviria_k__Bamfordvirae_______3300010034_Ga0126342_1000390620
IMGVR_UViG_3300010035_000019_r__Varidnaviria_k__Bamfordvirae_______3300010035_Ga0126343_100055768
IMGVR_UViG_3300010034_000041_r__Varidnaviria_k__Bamfordvirae_p__Preplasmiviricota_c__Polintoviricetes_o__Orthopolintovirales____3300010034_Ga0126342_1000146729
IMGVR_UViG_3300010034_000014_r__Varidnaviria_k__Bamfordvirae_______3300010034_Ga0126342_100482725
IMGVR_UViG_3300010034_000056_r__Varidnaviria_k__Bamfordvirae_p__Preplasmiviricota_c__Polintoviricetes_o__Orthopolintovirales____3300010034_Ga0126342_100450894
IMGVR_UViG_3300010034_000015_r__Varidnaviria_k__Bamfordvirae_p__Preplasmiviricota_c__Polintoviricetes_o__Orthopolintovirales____3300010034_Ga0126342_100232744
IMGVR_UViG_3300010035_000018_no_taxonomy_3300010035_Ga0126343_100248236
MAG_viral_055-00000001_6
IMGVR_UViG_3300010034_000049_r__Varidnaviria_k__Bamfordvirae_p__Nucleocytoviricota______3300010034_Ga0126342_100422125
IMGVR_UViG_3300010035_000035_r__Varidnaviria_k__Bamfordvirae_p__Nucleocytoviricota_c__Megaviricetes_o__Pimascovirales_f__Iridoviridae___3300010035_Ga0126343_100132317
IMGVR_UViG_3300017976_000002_r__Varidnaviria_k__Bamfordvirae_p__Nucleocytoviricota_c__Megaviricetes_o__Pimascovirales_f__Iridoviridae___3300017976_Ga0192488_10899837
IMGVR_UViG_3300010034_000014_r__Varidnaviria_k__Bamfordvirae_______3300010034_Ga0126342_100482726
IMGVR_UViG_3300010034_000021_r__Varidnaviria_k__Bamfordvirae_p__Preplasmiviricota_c__Polintoviricetes_o__Orthopolintovirales____3300010034_Ga0126342_1002228916
IMGVR_UViG_3300010034_000040_no_taxonomy_3300010034_Ga0126342_1001716721
IMGVR_UViG_3300010034_000009_r__Varidnaviria_k__Bamfordvirae_p__Preplasmiviricota_c__Polintoviricetes_o__Orthopolintovirales_f__Adintoviridae___3300010034_Ga0126342_1002116114
1
100
42
75
20
41
22
28
69
84
98
31
74
44
42
82
41
79
100
2.5

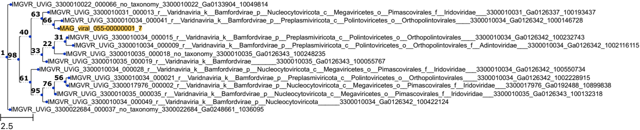
IMGVR_UViG_3300010022_000066_no_taxonomy_3300010022_Ga0133904_10049814
IMGVR_UViG_3300010031_000013_r__Varidnaviria_k__Bamfordvirae_p__Nucleocytoviricota_c__Megaviricetes_o__Pimascovirales_f__Iridoviridae___3300010031_Ga0126337_100193437
IMGVR_UViG_3300010034_000041_r__Varidnaviria_k__Bamfordvirae_p__Preplasmiviricota_c__Polintoviricetes_o__Orthopolintovirales____3300010034_Ga0126342_1000146728
MAG_viral_055-00000001_7
IMGVR_UViG_3300010034_000015_r__Varidnaviria_k__Bamfordvirae_p__Preplasmiviricota_c__Polintoviricetes_o__Orthopolintovirales____3300010034_Ga0126342_100232743
IMGVR_UViG_3300010034_000009_r__Varidnaviria_k__Bamfordvirae_p__Preplasmiviricota_c__Polintoviricetes_o__Orthopolintovirales_f__Adintoviridae___3300010034_Ga0126342_1002116115
IMGVR_UViG_3300010035_000018_no_taxonomy_3300010035_Ga0126343_100248235
IMGVR_UViG_3300010035_000019_r__Varidnaviria_k__Bamfordvirae_______3300010035_Ga0126343_100055767
IMGVR_UViG_3300010034_000028_r__Varidnaviria_k__Bamfordvirae_p__Nucleocytoviricota_c__Megaviricetes_o__Pimascovirales_f__Iridoviridae___3300010034_Ga0126342_100550734
IMGVR_UViG_3300010034_000021_r__Varidnaviria_k__Bamfordvirae_p__Preplasmiviricota_c__Polintoviricetes_o__Orthopolintovirales____3300010034_Ga0126342_1002228915
IMGVR_UViG_3300017976_000002_r__Varidnaviria_k__Bamfordvirae_p__Nucleocytoviricota_c__Megaviricetes_o__Pimascovirales_f__Iridoviridae___3300017976_Ga0192488_10899838
IMGVR_UViG_3300010035_000035_r__Varidnaviria_k__Bamfordvirae_p__Nucleocytoviricota_c__Megaviricetes_o__Pimascovirales_f__Iridoviridae___3300010035_Ga0126343_100132318
IMGVR_UViG_3300010034_000049_r__Varidnaviria_k__Bamfordvirae_p__Nucleocytoviricota______3300010034_Ga0126342_100422124
IMGVR_UViG_3300022684_000037_no_taxonomy_3300022684_Ga0248661_1036095
63
66
40
31
22
33
1
98
61
56
76
95
2.5

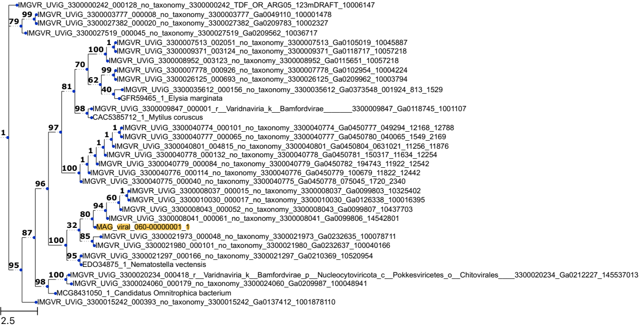
IMGVR_UViG_3300000242_000128_no_taxonomy_3300000242_TDF_OR_ARG05_123mDRAFT_10006147
IMGVR_UViG_3300003777_000008_no_taxonomy_3300003777_Ga0049110_100001478
IMGVR_UViG_3300027382_000020_no_taxonomy_3300027382_Ga0209783_10002327
IMGVR_UViG_3300027519_000045_no_taxonomy_3300027519_Ga0209562_10036717
IMGVR_UViG_3300007513_002051_no_taxonomy_3300007513_Ga0105019_10045887
IMGVR_UViG_3300009371_003124_no_taxonomy_3300009371_Ga0118717_10057218
IMGVR_UViG_3300008952_003123_no_taxonomy_3300008952_Ga0115651_10057218
IMGVR_UViG_3300007778_000926_no_taxonomy_3300007778_Ga0102954_10004224
IMGVR_UViG_3300026125_000693_no_taxonomy_3300026125_Ga0209962_10003794
IMGVR_UViG_3300035612_000156_no_taxonomy_3300035612_Ga0373548_001924_813_1529
GFR59465_1_Elysia marginata
IMGVR_UViG_3300009847_000001_r__Varidnaviria_k__Bamfordvirae________3300009847_Ga0118745_1001107
CAC5385712_1_Mytilus coruscus
IMGVR_UViG_3300040774_000101_no_taxonomy_3300040774_Ga0450777_049294_12168_12788
IMGVR_UViG_3300040777_000065_no_taxonomy_3300040777_Ga0450780_040065_1549_2169
IMGVR_UViG_3300040801_004815_no_taxonomy_3300040801_Ga0450804_0631021_11256_11876
IMGVR_UViG_3300040778_000132_no_taxonomy_3300040778_Ga0450781_150317_11634_12254
IMGVR_UViG_3300040779_000084_no_taxonomy_3300040779_Ga0450782_194743_11922_12542
IMGVR_UViG_3300040776_000114_no_taxonomy_3300040776_Ga0450779_100679_11822_12442
IMGVR_UViG_3300040775_000040_no_taxonomy_3300040775_Ga0450778_075045_1720_2340
IMGVR_UViG_3300008037_000015_no_taxonomy_3300008037_Ga0099803_10325402
IMGVR_UViG_3300010030_000017_no_taxonomy_3300010030_Ga0126338_100016395
IMGVR_UViG_3300008043_000052_no_taxonomy_3300008043_Ga0099807_10437703
IMGVR_UViG_3300008041_000061_no_taxonomy_3300008041_Ga0099806_14542801
MAG_viral_060-00000001_1
IMGVR_UViG_3300021973_000048_no_taxonomy_3300021973_Ga0232635_100078711
IMGVR_UViG_3300021980_000101_no_taxonomy_3300021980_Ga0232637_100040166
IMGVR_UViG_3300021297_000166_no_taxonomy_3300021297_Ga0210369_10520954
EDO34875_1_Nematostella vectensis
IMGVR_UViG_3300020234_000418_r__Varidnaviria_k__Bamfordvirae_p__Nucleocytoviricota_c__Pokkesviricetes_o__Chitovirales____3300020234_Ga0212227_145537013
IMGVR_UViG_3300024060_000179_no_taxonomy_3300024060_Ga0209987_100048941
MCG8431050_1_Candidatus Omnitrophica bacterium
IMGVR_UViG_3300015242_000393_no_taxonomy_3300015242_Ga0137412_1001878110
2.5

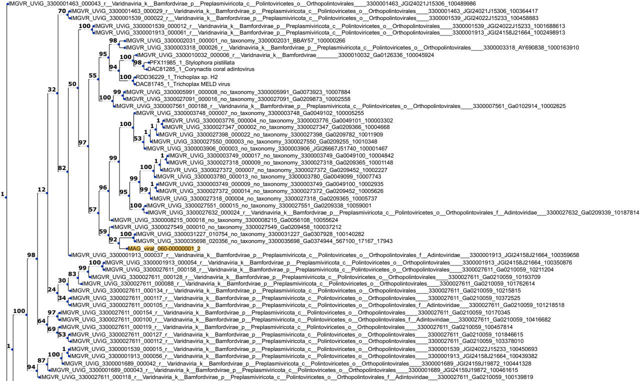
IMGVR_UViG_3300001463_000043_r__Varidnaviria_k__Bamfordvirae_p__Preplasmiviricota_c__Polintoviricetes_o__Orthopolintovirales____3300001463_JGI24021J15306_100489986
IMGVR_UViG_3300001463_000029_r__Varidnaviria_k__Bamfordvirae_p__Preplasmiviricota_c__Polintoviricetes_o__Orthopolintovirales____3300001463_JGI24021J15306_100364417
IMGVR_UViG_3300001539_000022_r__Varidnaviria_k__Bamfordvirae_p__Preplasmiviricota_c__Polintoviricetes_o__Orthopolintovirales____3300001539_JGI24022J15233_100458883
IMGVR_UViG_3300001539_000012_r__Varidnaviria_k__Bamfordvirae_p__Preplasmiviricota_c__Polintoviricetes_o__Orthopolintovirales____3300001539_JGI24022J15233_1001688613
IMGVR_UViG_3300001913_000061_r__Varidnaviria_k__Bamfordvirae_p__Preplasmiviricota_c__Polintoviricetes_o__Orthopolintovirales____3300001913_JGI24158J21664_1002498913
IMGVR_UViG_3300002031_000001_no_taxonomy_3300002031_BBAY57_100000266
IMGVR_UViG_3300003318_000026_r__Varidnaviria_k__Bamfordvirae_p__Preplasmiviricota_c__Polintoviricetes_o__Orthopolintovirales____3300003318_AY690838_1000163910
IMGVR_UViG_3300010032_000006_r__Varidnaviria_k__Bamfordvirae_______3300010032_Ga0126336_100045924
PFX11985_1_Stylophora pistillata
DAC81285_1_Corynactis coral adintovirus
RDD36229_1_Trichoplax sp. H2
DAC81745_1_Trichoplax MELD virus
IMGVR_UViG_3300005991_000008_no_taxonomy_3300005991_Ga0073923_10007884
IMGVR_UViG_3300027091_000016_no_taxonomy_3300027091_Ga0209873_10002558
IMGVR_UViG_3300007561_000188_r__Varidnaviria_k__Bamfordvirae_p__Preplasmiviricota_c__Polintoviricetes_o__Orthopolintovirales____3300007561_Ga0102914_10002625
IMGVR_UViG_3300003748_000007_no_taxonomy_3300003748_Ga0049102_100005255
IMGVR_UViG_3300003776_000004_no_taxonomy_3300003776_Ga0049101_100003302
IMGVR_UViG_3300027347_000002_no_taxonomy_3300027347_Ga0209366_10004668
IMGVR_UViG_3300027398_000022_no_taxonomy_3300027398_Ga0209782_10011909
IMGVR_UViG_3300027550_000003_no_taxonomy_3300027550_Ga0209255_10010348
IMGVR_UViG_3300003906_000003_no_taxonomy_3300003906_JGI26667J51740_100001467
IMGVR_UViG_3300003749_000017_no_taxonomy_3300003749_Ga0049100_10004842
IMGVR_UViG_3300027318_000009_no_taxonomy_3300027318_Ga0209365_10001148
IMGVR_UViG_3300027372_000007_no_taxonomy_3300027372_Ga0209452_10002227
IMGVR_UViG_3300003780_000013_no_taxonomy_3300003780_Ga0049099_10007743
IMGVR_UViG_3300003749_000009_no_taxonomy_3300003749_Ga0049100_10002935
IMGVR_UViG_3300027372_000014_no_taxonomy_3300027372_Ga0209452_10005626
IMGVR_UViG_3300027318_000004_no_taxonomy_3300027318_Ga0209365_10005737
IMGVR_UViG_3300027551_000015_no_taxonomy_3300027551_Ga0209338_10059001
IMGVR_UViG_3300027632_000024_r__Varidnaviria_k__Bamfordvirae_p__Preplasmiviricota_c__Polintoviricetes_o__Orthopolintovirales_f__Adintoviridae___3300027632_Ga0209339_10187814
IMGVR_UViG_3300008215_000018_no_taxonomy_3300008215_Ga0056108_10055624
IMGVR_UViG_3300027549_000010_no_taxonomy_3300027549_Ga0209458_100037212
IMGVR_UViG_3300031227_010754_no_taxonomy_3300031227_Ga0307928_100140282
IMGVR_UViG_3300035698_020356_no_taxonomy_3300035698_Ga0374944_567100_17167_17943
MAG_viral_060-00000001_2
IMGVR_UViG_3300001913_000037_r__Varidnaviria_k__Bamfordvirae_p__Preplasmiviricota_c__Polintoviricetes_o__Orthopolintovirales_f__Adintoviridae___3300001913_JGI24158J21664_100359658
IMGVR_UViG_3300001913_000054_r__Varidnaviria_k__Bamfordvirae_p__Preplasmiviricota_c__Polintoviricetes_o__Orthopolintovirales____3300001913_JGI24158J21664_100350876
IMGVR_UViG_3300027611_000158_r__Varidnaviria_k__Bamfordvirae_p__Preplasmiviricota_c__Polintoviricetes_o__Orthopolintovirales____3300027611_Ga0210059_10211204
IMGVR_UViG_3300027611_000128_r__Varidnaviria_k__Bamfordvirae_p__Preplasmiviricota_c__Polintoviricetes_o__Orthopolintovirales____3300027611_Ga0210059_10193709
IMGVR_UViG_3300027611_000088_r__Varidnaviria_k__Bamfordvirae_p__Preplasmiviricota_c__Polintoviricetes_o__Orthopolintovirales____3300027611_Ga0210059_101762614
IMGVR_UViG_3300027611_000134_r__Varidnaviria_k__Bamfordvirae_p__Preplasmiviricota_c__Polintoviricetes_o__Orthopolintovirales____3300027611_Ga0210059_10215815
IMGVR_UViG_3300027611_000117_r__Varidnaviria_k__Bamfordvirae_p__Preplasmiviricota_c__Polintoviricetes_o__Orthopolintovirales____3300027611_Ga0210059_10372525
IMGVR_UViG_3300027611_000105_r__Varidnaviria_k__Bamfordvirae_p__Preplasmiviricota_c__Polintoviricetes_o__Orthopolintovirales_f__Adintoviridae___3300027611_Ga0210059_101218518
IMGVR_UViG_3300027611_000154_r__Varidnaviria_k__Bamfordvirae_p__Preplasmiviricota_c__Polintoviricetes_o__Orthopolintovirales____3300027611_Ga0210059_10170345
IMGVR_UViG_3300027611_000100_r__Varidnaviria_k__Bamfordvirae_p__Preplasmiviricota_c__Polintoviricetes_o__Orthopolintovirales_f__Adintoviridae___3300027611_Ga0210059_10416682
IMGVR_UViG_3300027611_000119_r__Varidnaviria_k__Bamfordvirae_p__Preplasmiviricota_c__Polintoviricetes_o__Orthopolintovirales____3300027611_Ga0210059_100457814
IMGVR_UViG_3300027611_000127_r__Varidnaviria_k__Bamfordvirae_p__Preplasmiviricota_c__Polintoviricetes_o__Orthopolintovirales____3300027611_Ga0210059_101846615
IMGVR_UViG_3300027611_000112_r__Varidnaviria_k__Bamfordvirae_p__Preplasmiviricota_c__Polintoviricetes_o__Orthopolintovirales____3300027611_Ga0210059_103378010
IMGVR_UViG_3300001539_000015_r__Varidnaviria_k__Bamfordvirae_p__Preplasmiviricota_c__Polintoviricetes_o__Orthopolintovirales____3300001539_JGI24022J15233_100450693
IMGVR_UViG_3300001913_000056_r__Varidnaviria_k__Bamfordvirae_p__Preplasmiviricota_c__Polintoviricetes_o__Orthopolintovirales____3300001913_JGI24158J21664_100439382
IMGVR_UViG_3300001689_000042_r__Varidnaviria_k__Bamfordvirae_p__Preplasmiviricota_c__Polintoviricetes_o__Orthopolintovirales____3300001689_JGI24159J19872_100441328
IMGVR_UViG_3300001689_000043_r__Varidnaviria_k__Bamfordvirae_p__Preplasmiviricota_c__Polintoviricetes_o__Orthopolintovirales____3300001689_JGI24159J19872_100461615
IMGVR_UViG_3300027611_000118_r__Varidnaviria_k__Bamfordvirae_p__Preplasmiviricota_c__Polintoviricetes_o__Orthopolintovirales_f__Adintoviridae___3300027611_Ga0210059_100139819
IMGVR_UViG_3300001689_000031_r__Varidnaviria_k__Bamfordvirae_p__Preplasmiviricota_c__Polintoviricetes_o__Orthopolintovirales____3300001689_JGI24159J19872_1003206210
IMGVR_UViG_3300001913_000042_r__Varidnaviria_k__Bamfordvirae_p__Preplasmiviricota_c__Polintoviricetes_o__Orthopolintovirales____3300001913_JGI24158J21664_100314159
2.5

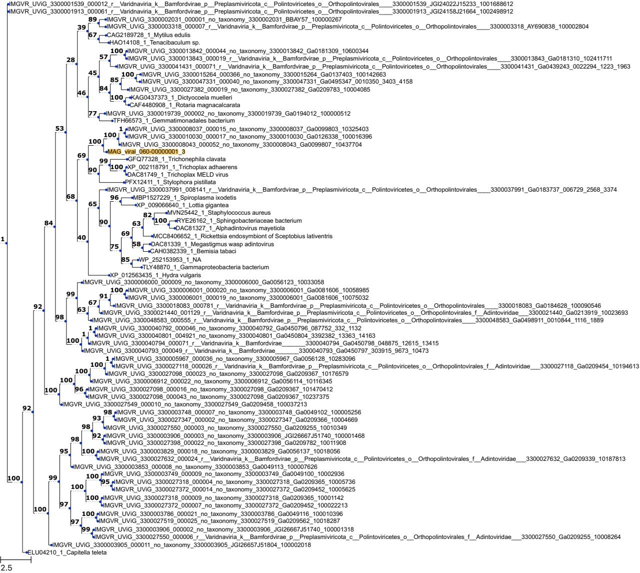
IMGVR_UViG_3300001539_000012_r__Varidnaviria_k__Bamfordvirae_p__Preplasmiviricota_c__Polintoviricetes_o__Orthopolintovirales____3300001539_JGI24022J15233_1001688612
IMGVR_UViG_3300001913_000061_r__Varidnaviria_k__Bamfordvirae_p__Preplasmiviricota_c__Polintoviricetes_o__Orthopolintovirales____3300001913_JGI24158J21664_1002498912
IMGVR_UViG_3300002031_000001_no_taxonomy_3300002031_BBAY57_100000267
IMGVR_UViG_3300003318_000007_r__Varidnaviria_k__Bamfordvirae_p__Preplasmiviricota_c__Polintoviricetes_o__Orthopolintovirales____3300003318_AY690838_100002804
CAG2189728_1_Mytilus edulis
HAO14108_1_Tenacibaculum sp.
IMGVR_UViG_3300013842_000044_no_taxonomy_3300013842_Ga0181309_10600344
IMGVR_UViG_3300013843_000019_r__Varidnaviria_k__Bamfordvirae_p__Preplasmiviricota_c__Polintoviricetes_o__Orthopolintovirales____3300013843_Ga0181310_102411711
IMGVR_UViG_3300041431_000071_r__Varidnaviria_k__Bamfordvirae_p__Preplasmiviricota_c__Polintoviricetes_o__Orthopolintovirales____3300041431_Ga0439243_0022294_1223_1963
IMGVR_UViG_3300015264_000366_no_taxonomy_3300015264_Ga0137403_100142663
IMGVR_UViG_3300047331_000040_no_taxonomy_3300047331_Ga0495347_0010350_3403_4158
IMGVR_UViG_3300027382_000019_no_taxonomy_3300027382_Ga0209783_10004085
KAG0437373_1_Dictyocoela muelleri
CAF4480908_1_Rotaria magnacalcarata
IMGVR_UViG_3300019739_000002_no_taxonomy_3300019739_Ga0194012_100000512
TFH66573_1_Gemmatimonadales bacterium
IMGVR_UViG_3300008037_000015_no_taxonomy_3300008037_Ga0099803_10325403
IMGVR_UViG_3300010030_000017_no_taxonomy_3300010030_Ga0126338_100016396
IMGVR_UViG_3300008043_000052_no_taxonomy_3300008043_Ga0099807_10437704
MAG_viral_060-00000001_3
GFQ77328_1_Trichonephila clavata
XP_002118791_1_Trichoplax adhaerens
DAC81749_1_Trichoplax MELD virus
PFX12411_1_Stylophora pistillata
IMGVR_UViG_3300037991_008141_r__Varidnaviria_k__Bamfordvirae_p__Preplasmiviricota_c__Polintoviricetes_o__Orthopolintovirales____3300037991_Ga0183737_006729_2568_3374
MBP1527229_1_Spiroplasma ixodetis
XP_009066640_1_Lottia gigantea
MVN25442_1_Staphylococcus aureus
RYE26162_1_Sphingobacteriaceae bacterium
DAC81327_1_Alphadintovirus mayetiola
MCC8406652_1_Rickettsia endosymbiont of Sceptobius lativentris
DAC81339_1_Megastigmus wasp adintovirus
CAH0382339_1_Bemisia tabaci
WP_252153953_1_NA
TLY48870_1_Gammaproteobacteria bacterium
XP_012563435_1_Hydra vulgaris
IMGVR_UViG_3300006000_000009_no_taxonomy_3300006000_Ga0056123_10033058
IMGVR_UViG_3300006001_000020_no_taxonomy_3300006001_Ga0081606_10058985
IMGVR_UViG_3300006001_000019_no_taxonomy_3300006001_Ga0081606_10075032
IMGVR_UViG_3300018083_000781_r__Varidnaviria_k__Bamfordvirae_p__Preplasmiviricota_c__Polintoviricetes_o__Orthopolintovirales____3300018083_Ga0184628_100090546
IMGVR_UViG_3300021440_001129_r__Varidnaviria_k__Bamfordvirae_p__Preplasmiviricota_c__Polintoviricetes_o__Orthopolintovirales_f__Adintoviridae___3300021440_Ga0213919_10023693
IMGVR_UViG_3300048583_000555_r__Varidnaviria_k__Bamfordvirae_p__Preplasmiviricota_c__Polintoviricetes_o__Orthopolintovirales____3300048583_Ga0498911_0010844_1116_1889
IMGVR_UViG_3300040792_000046_no_taxonomy_3300040792_Ga0450796_087752_332_1132
IMGVR_UViG_3300040801_004921_no_taxonomy_3300040801_Ga0450804_3392382_13363_14163
IMGVR_UViG_3300040794_000071_r__Varidnaviria_k__Bamfordvirae________3300040794_Ga0450798_048875_12615_13415
IMGVR_UViG_3300040793_000049_r__Varidnaviria_k__Bamfordvirae________3300040793_Ga0450797_303915_9673_10473
IMGVR_UViG_3300005967_000036_no_taxonomy_3300005967_Ga0056128_10283096
IMGVR_UViG_3300027118_000026_r__Varidnaviria_k__Bamfordvirae_p__Preplasmiviricota_c__Polintoviricetes_o__Orthopolintovirales_f__Adintoviridae___3300027118_Ga0209454_10194613
IMGVR_UViG_3300027098_000023_no_taxonomy_3300027098_Ga0209367_10176579
IMGVR_UViG_3300006912_000022_no_taxonomy_3300006912_Ga0056114_10116345
IMGVR_UViG_3300027098_000016_no_taxonomy_3300027098_Ga0209367_101470412
IMGVR_UViG_3300027098_000043_no_taxonomy_3300027098_Ga0209367_10237375
IMGVR_UViG_3300027549_000010_no_taxonomy_3300027549_Ga0209458_100037213
IMGVR_UViG_3300003748_000007_no_taxonomy_3300003748_Ga0049102_100005256
IMGVR_UViG_3300027347_000002_no_taxonomy_3300027347_Ga0209366_10004669
IMGVR_UViG_3300027550_000003_no_taxonomy_3300027550_Ga0209255_10010349
IMGVR_UViG_3300003906_000003_no_taxonomy_3300003906_JGI26667J51740_100001468
IMGVR_UViG_3300027398_000022_no_taxonomy_3300027398_Ga0209782_10011908
IMGVR_UViG_3300003829_000018_no_taxonomy_3300003829_Ga0056137_10018056
IMGVR_UViG_3300027632_000024_r__Varidnaviria_k__Bamfordvirae_p__Preplasmiviricota_c__Polintoviricetes_o__Orthopolintovirales_f__Adintoviridae___3300027632_Ga0209339_10187813
IMGVR_UViG_3300003853_000008_no_taxonomy_3300003853_Ga0049113_10007626
IMGVR_UViG_3300003749_000009_no_taxonomy_3300003749_Ga0049100_10002936
IMGVR_UViG_3300027318_000004_no_taxonomy_3300027318_Ga0209365_10005736
IMGVR_UViG_3300027372_000014_no_taxonomy_3300027372_Ga0209452_10005625
IMGVR_UViG_3300027318_000009_no_taxonomy_3300027318_Ga0209365_10001142
IMGVR_UViG_3300027372_000007_no_taxonomy_3300027372_Ga0209452_100022213
IMGVR_UViG_3300003786_000021_no_taxonomy_3300003786_Ga0049116_100010396
IMGVR_UViG_3300027519_000025_no_taxonomy_3300027519_Ga0209562_10018287
IMGVR_UViG_3300003906_000002_no_taxonomy_3300003906_JGI26667J51740_100001318
IMGVR_UViG_3300027550_000006_r__Varidnaviria_k__Bamfordvirae_p__Preplasmiviricota_c__Polintoviricetes_o__Orthopolintovirales_f__Adintoviridae___3300027550_Ga0209255_10008264
IMGVR_UViG_3300003905_000011_no_taxonomy_3300003905_JGI26657J51804_100002018
ELU04210_1_Capitella teleta
2.5

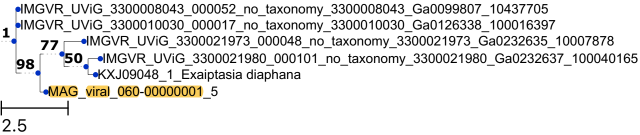
IMGVR_UViG_3300008043_000052_no_taxonomy_3300008043_Ga0099807_10437705
IMGVR_UViG_3300010030_000017_no_taxonomy_3300010030_Ga0126338_100016397
1
77
IMGVR_UViG_3300021973_000048_no_taxonomy_3300021973_Ga0232635_10007878
50
IMGVR_UViG_3300021980_000101_no_taxonomy_3300021980_Ga0232637_100040165
98
KXJ09048_1_Exaiptasia diaphana
MAG_viral_060-00000001_5
2.5

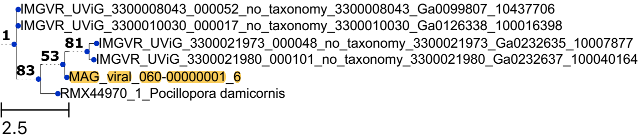

IMGVR_UViG_3300008043_000052_no_taxonomy_3300008043_Ga0099807_10437706
IMGVR_UViG_3300010030_000017_no_taxonomy_3300010030_Ga0126338_100016398
1
81
IMGVR_UViG_3300021973_000048_no_taxonomy_3300021973_Ga0232635_10007877
53
IMGVR_UViG_3300021980_000101_no_taxonomy_3300021980_Ga0232637_100040164
83
MAG_viral_060-00000001_6
RMX44970_1_Pocillopora damicornis
2.5

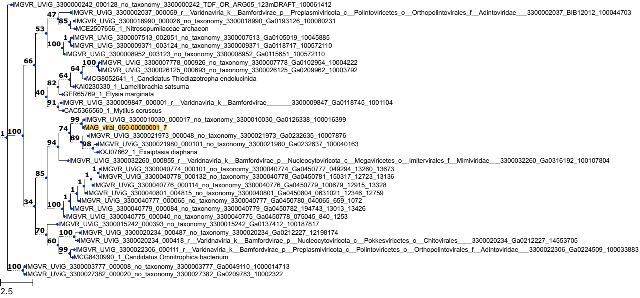
IMGVR_UViG_3300000242_000128_no_taxonomy_3300000242_TDF_OR_ARG05_123mDRAFT_100061412
IMGVR_UViG_3300002037_000059_r__Varidnaviria_k__Bamfordvirae_p__Preplasmiviricota_c__Polintoviricetes_o__Orthopolintovirales_f__Adintoviridae___3300002037_BIB12012_100044703
IMGVR_UViG_3300018990_000026_no_taxonomy_3300018990_Ga0193126_100080231
MCE2507656_1_Nitrosopumilaceae archaeon
IMGVR_UViG_3300007513_002051_no_taxonomy_3300007513_Ga0105019_10045885
IMGVR_UViG_3300009371_003124_no_taxonomy_3300009371_Ga0118717_100572110
IMGVR_UViG_3300008952_003123_no_taxonomy_3300008952_Ga0115651_100572110
IMGVR_UViG_3300007778_000926_no_taxonomy_3300007778_Ga0102954_10004222
IMGVR_UViG_3300026125_000693_no_taxonomy_3300026125_Ga0209962_10003792
MCG8052641_1_Candidatus Thiodiazotropha endolucinida
KAI0230330_1_Lamellibrachia satsuma
GFR65769_1_Elysia marginata
IMGVR_UViG_3300009847_000001_r__Varidnaviria_k__Bamfordvirae_______3300009847_Ga0118745_1001104
CAC5366560_1_Mytilus coruscus
IMGVR_UViG_3300010030_000017_no_taxonomy_3300010030_Ga0126338_100016399
MAG_viral_060-00000001_7
IMGVR_UViG_3300021973_000048_no_taxonomy_3300021973_Ga0232635_10007876
IMGVR_UViG_3300021980_000101_no_taxonomy_3300021980_Ga0232637_100040163
KXJ07862_1_Exaiptasia diaphana
IMGVR_UViG_3300032260_000855_r__Varidnaviria_k__Bamfordvirae_p__Nucleocytoviricota_c__Megaviricetes_o__Imitervirales_f__Mimiviridae___3300032260_Ga0316192_100107804
IMGVR_UViG_3300040774_000101_no_taxonomy_3300040774_Ga0450777_049294_13260_13673
IMGVR_UViG_3300040778_000132_no_taxonomy_3300040778_Ga0450781_150317_12723_13136
IMGVR_UViG_3300040776_000114_no_taxonomy_3300040776_Ga0450779_100679_12915_13328
IMGVR_UViG_3300040801_004815_no_taxonomy_3300040801_Ga0450804_0631021_12346_12759
IMGVR_UViG_3300040777_000065_no_taxonomy_3300040777_Ga0450780_040065_659_1072
IMGVR_UViG_3300040779_000084_no_taxonomy_3300040779_Ga0450782_194743_13013_13426
IMGVR_UViG_3300040775_000040_no_taxonomy_3300040775_Ga0450778_075045_840_1253
IMGVR_UViG_3300015242_000393_no_taxonomy_3300015242_Ga0137412_100187817
IMGVR_UViG_3300020234_000487_no_taxonomy_3300020234_Ga0212227_12198174
IMGVR_UViG_3300020234_000418_r__Varidnaviria_k__Bamfordvirae_p__Nucleocytoviricota_c__Pokkesviricetes_o__Chitovirales____3300020234_Ga0212227_14553705
IMGVR_UViG_3300022306_000111_r__Varidnaviria_k__Bamfordvirae_p__Preplasmiviricota_c__Polintoviricetes_o__Orthopolintovirales_f__Adintoviridae___3300022306_Ga0224509_100033883
MCG8430990_1_Candidatus Omnitrophica bacterium
IMGVR_UViG_3300003777_000008_no_taxonomy_3300003777_Ga0049110_1000014713
IMGVR_UViG_3300027382_000020_no_taxonomy_3300027382_Ga0209783_10002322
2.5

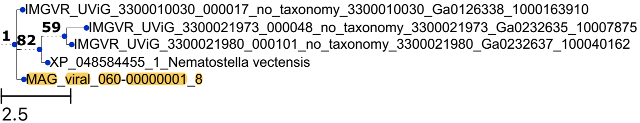
IMGVR_UViG_3300010030_000017_no_taxonomy_3300010030_Ga0126338_1000163910
59
IMGVR_UViG_3300021973_000048_no_taxonomy_3300021973_Ga0232635_10007875
1
82
IMGVR_UViG_3300021980_000101_no_taxonomy_3300021980_Ga0232637_100040162
XP_048584455_1_Nematostella vectensis
MAG_viral_060-00000001_8
2.5
